# Identification of MAZ as a novel transcription factor regulating erythropoiesis

**DOI:** 10.1101/2020.05.10.087254

**Authors:** Darya Deen, Falk Butter, Michelle L. Holland, Vasiliki Samara, Jacqueline A. Sloane-Stanley, Helena Ayyub, Matthias Mann, David Garrick, Douglas Vernimmen

## Abstract

Erythropoiesis requires a combination of ubiquitous and tissue-specific transcription factors. Here, through DNA affinity purification followed by mass spectrometry, we have identified the widely expressed protein MAZ (Myc-associated zinc finger) as a transcription factor that binds to the promoter of the erythroid-specific human α-globin gene. Genome-wide mapping in primary human erythroid cells revealed that MAZ also occupies active promoters as well as GATA1-bound enhancer elements of key erythroid genes. Consistent with an important role during erythropoiesis, knockdown of MAZ in primary human erythroid cells impairs erythroid differentiation, and genetic variants in the *MAZ* locus are associated with clinically important human erythroid traits. Taken together, these findings reveal the Zinc-finger transcription factor MAZ to be a previously unrecognised regulator of the erythroid differentiation program.

## INTRODUCTION

Over the last three decades, studies of the α- and β-globin gene clusters have contributed to our understanding of some of the fundamental principles of mammalian gene regulation, including RNA stability, termination of transcription, and the identification of remote regulatory elements^1^. A complex network of DNA sequence elements, chromatin accessibility, histone modifications, and transcription factor (TF) occupancy orchestrates expression of globin genes, which are exclusively expressed in erythroid cells ^2^. Both proximal regulatory regions (promoters) and distal regulatory elements (enhancers) are required for the initiation of α-globin (*HBA*) and β-globin (*HBB*) gene expression in erythroid cells. When active, these regulatory elements are characterised by the presence of DNase I hypersensitivity sites (DHS) and active histone modifications. A small group of lineage-restricted TFs including GATA binding protein 1 (GATA1), T cell acute lymphocytic leukemia 1 protein (TAL1), and Erythroid Krüppel-like factor (EKLF; henceforth referred to as KLF1) act as erythroid ‘master regulators ^3^, by binding to both promoters and enhancers of the globin genes as well as to other genes important for erythropoiesis, thereby significantly contributing to their expression [reviewed in ^2,4,5^].

Genetic and biochemical dissection of gene regulation programs has revealed that transcription of cell type-specific genes is usually achieved through synergy and cooperation between ubiquitously expressed TFs and tissue- or developmental stage-specific TFs ^6,7^. Although this concept is well established, the mechanisms of interactions and relationships between tissue-specific and ubiquitous TFs are generally lacking. Moreover, the broad range of phenotypic abnormalities associated with mutations/alterations in ubiquitously expressed regulators preclude dissecting their roles in specific processes and determining their contribution to disease states. Here, we have carried out an unbiased screen for proteins that bind to adjacent GC-rich motifs in the promoter of the duplicated α-globin genes (HBA2 and HBA1) in human erythroid cells, combining electrophoretic mobility shift assays (EMSAs), DNA affinity purification, and mass spectrometry. This screen identified <u>M</u>yc-<u>A</u>ssociated <u>Z</u>inc-finger protein (MAZ) as a direct binder of the promoter region of the human α-globin gene *in vitro* and *in vivo*. Knockdown of MAZ in primary human erythroid cells led to both reduction of α-globin expression and a block of erythroid differentiation. Moreover, variants in the MAZ gene itself and its promoter region are associated with clinically important human erythroid traits. By ChIP-seq in primary human erythroblasts we showed that MAZ is enriched at transcription start sites (TSSs) of transcriptionally active genes and distal regulatory elements where it recognises a canonical G_3_(C/A)G_4_ binding motif. Erythroid-specific MAZ signal is enriched at promoters and enhancers of genes associated with erythropoietic disorders. We found that MAZ erythroid-specific binding sites frequently colocalize with GATA1, a key regulator of erythropoiesis, particularly at enhancer elements, suggesting functional synergy between these two transcription factors. Together our findings have identified MAZ as an important regulator of the erythroid differentiation program.

## RESULTS

### A novel DNA-protein complex occupies the distal promoter region of the human HBA genes

Transcriptional activation of the HBA genes during erythroid differentiation is associated with localised relaxation of chromatin structure, which is observed experimentally as a DHS immediately upstream (promoter region) of the adult HBA genes (chr16: 172k-176k, Suppl. Figure 1A) specifically in erythroid cells^8^. In order to map chromatin accessibility within this erythroid-specific DHS at a higher resolution than provided by DNase- and ATAC-seq approaches, intact nuclei isolated from cells expressing *α*-globin (the erythroid cell line K562) and not expressing *α*-globin (EBV-transformed B lymphoblast cell line) were digested with low concentrations of selected restriction enzymes. The extent of digestion reflects accessibility of the specific recognition sites and was assessed by Southern blotting (Suppl. Figure 1B). This assay revealed that the region of erythroid-specific sensitivity extends from −220 bp (*Fok*I site) to +35 bp (*Nco*I site) relative to the TSS of the HBA genes (Suppl. Figure 1B).

**Figure 1.**
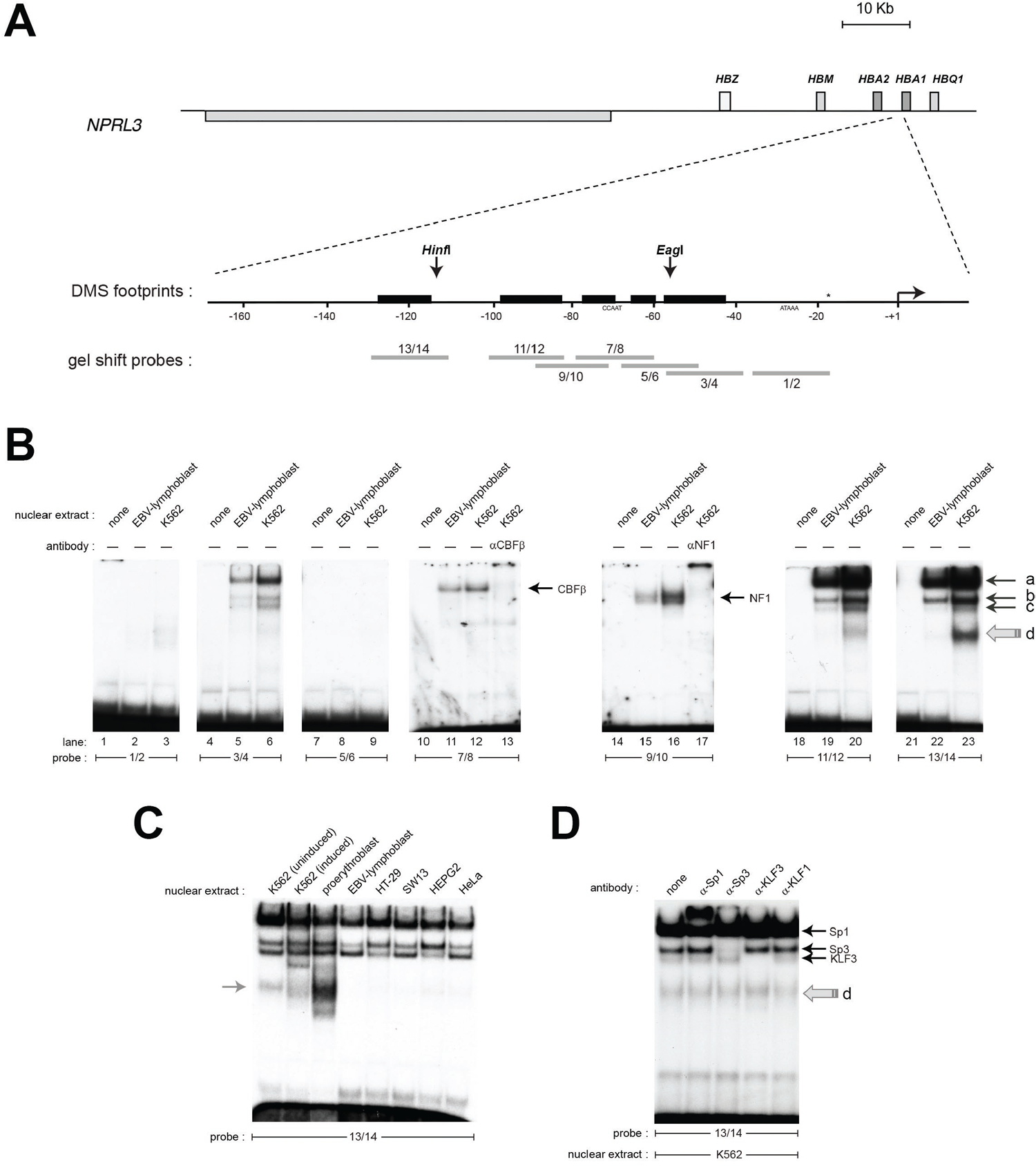
An erythroid-enriched complex binds a distal element of the HBA promoter. **(A)** Structure of the HBA core and distal promoter elements. The TSS is depicted as an angled arrow, and the locations of the EMSA probes are shown as grey bars. *In vivo* DMS footprints detected in K562 and erythroblasts ^9^ are represented by the black boxes. The nucleotide located 18 bp downstream of the TATA box (*) differs between the HBA1 (C) and HBA2 (G) promoters. **(B)** EMSA and supershift assays using K562 and EBV nuclear extracts with the indicated probes. **(C)** The probe 13/14 binds an additional protein enriched in erythroid cells. **(D)** Sp/X-KLF-family antibodies cause retardation or disruption of the bands (a), (b) and (c), but not (d).

In order to characterise protein binding across this hypersensitive region, we carried out electrophoretic mobility shift assays (EMSAs) using a series of overlapping oligonucleotide probes to compare *in vitro* binding proteins in nuclear extracts prepared from the same erythroid (K562) and non-erythroid (EBV) cells. Oligonucleotide probes were specifically placed at positions suggestive of protein binding by *in vivo* dimethyl-sulfate (DMS) footprinting assays reported previously (Figure 1A, Table 1) ^9^.

**Table 1.**
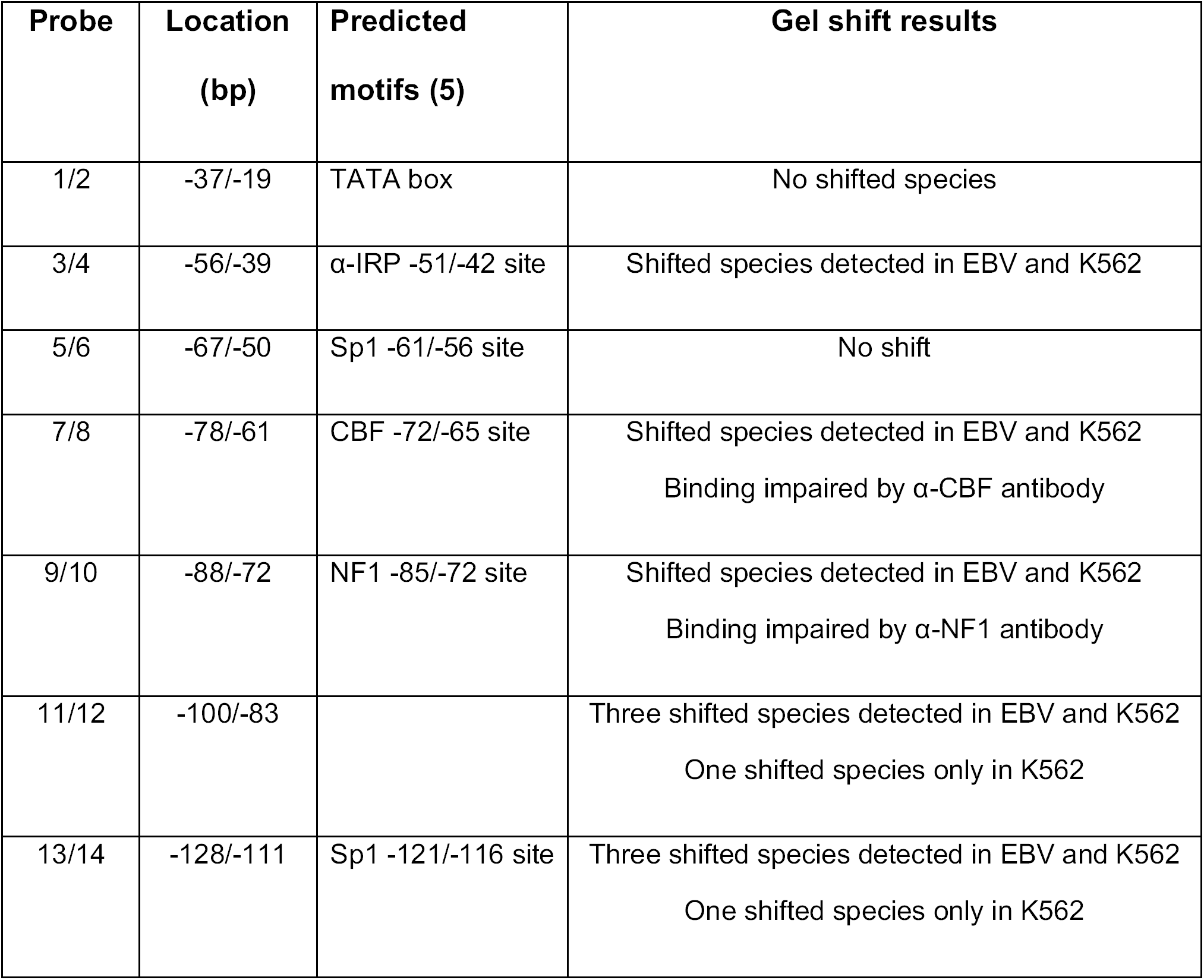
EMSA Results

We observed band shifts for five out of seven probes used (Figure 1B). Previous motif analysis had suggested potential binding motifs for SP1 (−121/-116, −61/-56), CBF (− 72/-65), Nuclear Factor I (NF1) (−85/-72), and *α*-inverted repeat protein (*α*-IRP) (−51/- 42) ^9^ (Table 1). Gel supershift assays, in which nuclear extracts were pre-incubated with anti-CBF or anti-NF1 antibodies, affect the retarded bands at probes 7/8 and 9/10 respectively, confirming that these sites are indeed bound by CBF and NF1 (Figure 1B, lanes 13 and 17, respectively). Interestingly, complex binding patterns comprised of four shifted bands were observed at two neighbouring GC boxes situated at −100/- 83 (probe 11/12) and −128/-111 (probe 13/14). These GC boxes have a highly similar sequence (Suppl. Figure 2A) and were able to cross-compete with each other in competition assays (Suppl. Figure 2B), suggesting that they are likely to be bound by the same proteins. Interestingly, while the most heavily retarded bands (labelled a, b and c in Figure 1B) were detected in both K562 and EBV nuclear extracts, the faster migrating band (d) was observed only in K562 cells. This factor bound more strongly to probe 13/14 than to 11/12 (Figure 1B, compare lanes 20 and 23), a finding that was confirmed in competition assays (Suppl. Figure 2B). Gel shifts carried out with nuclear extracts from other cell types revealed that band (d) was the most abundant complex in primary human erythroblasts but was either not detected or weak in a range of non-erythroid cells types (HT-29, SW13, HepG2, and HeLa) (Figure 1C).

**Figure 2.**
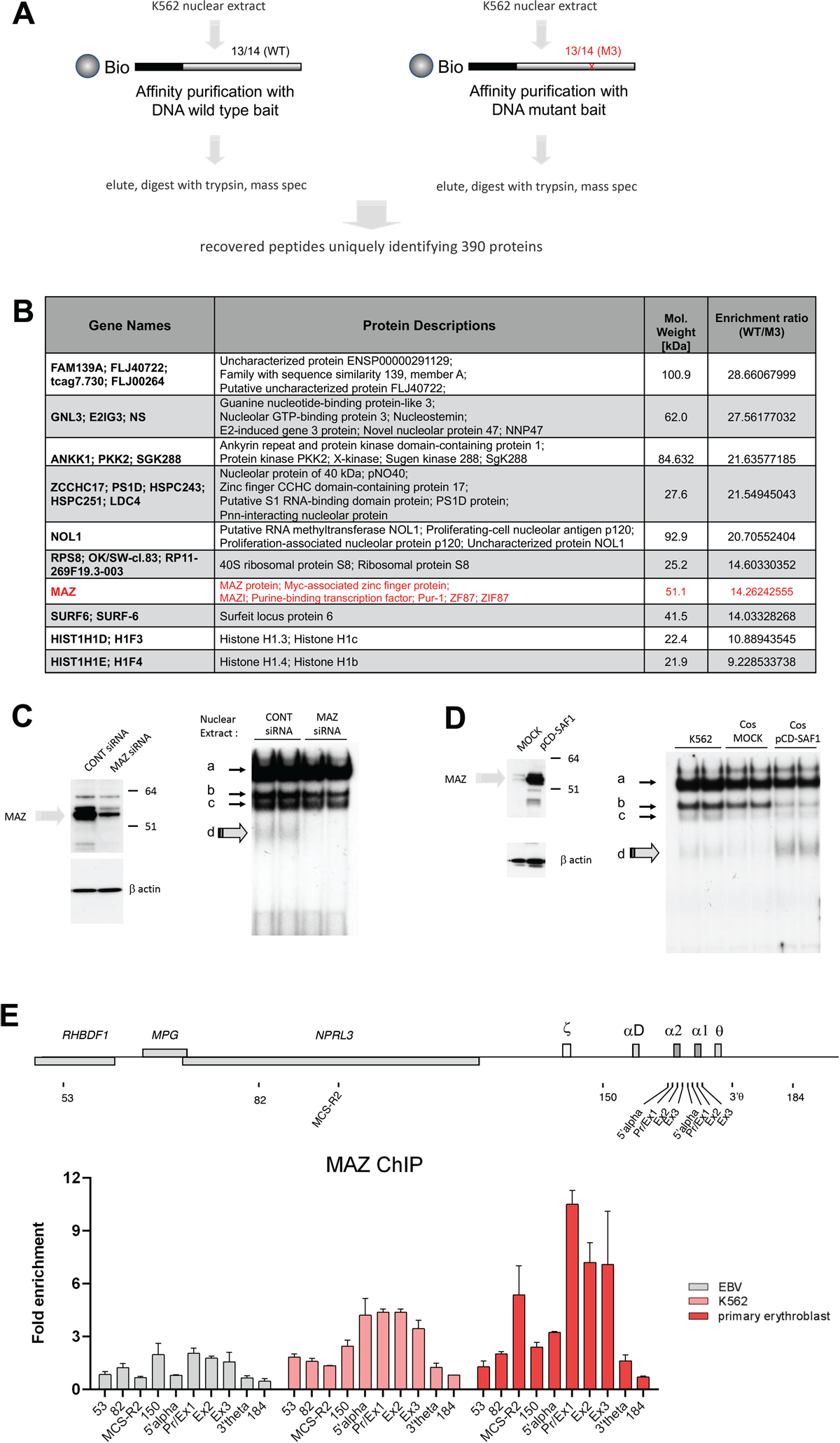
Identification of MAZ by mass spectrometry. **(A)** Schematic representation of the affinity purification screen. **(B)** The 10 proteins identified by mass spectrometry which were most enriched for binding to the wild type probe relative to the mutant. MAZ is highlighted in red. **(C)** and **(D)** EMSA showing that the intensity of band (d) is dependent on MAZ expression using knockdown (C) or overexpression (D) experiments. Knockdown experiments (siRNA) in K562 showed a strong reduction of MAZ protein levels and reduction of complex (d) in EMSA assays. Ectopic expression of MAZ (SAF1) in COS7 cells strongly enhanced complex (d) in EMSA assays. **(E)** MAZ is recruited to the active *α*-globin promoter *in vivo*. Analysis of MAZ binding at the *α*-globin locus in EBV-lymphoblasts, K562 cells and human primary erythroid cells by ChIP-qPCR. The y axis represents enrichment over the input DNA, normalised to a control sequence in the human 18S gene. The x axis indicates the Taqman probes used. The position of probes within the *α*-globin cluster are indicated on the heading map. The α-globin genes themselves are covered by three probes (Pr/Ex1, Ex2, Ex3). Error bars correspond to one SEM from two independent ChIPs.

The probe 13/14 (covering −128/-111 region) contains a predicted binding motif for the Krüppel-like zinc-finger transcription factor Sp1 ^9^ (Table1). We found that the binding of all four proteins to probe 13/14 was sensitive to the presence of the zinc-chelating agent EDTA (Suppl. Figure 3A), consistent with Zn-finger-dependent binding. In contrast, binding of the non-Zn-finger protein CBF to probe 7/8 was not affected by EDTA (Suppl. Figure 3A). To identify the protein responsible for these bands on probe 13/14, we carried out gel supershift reactions using antibodies against candidate Zn-finger proteins (Sp1, Sp3, Krüppel-like factor 1 (EKLF), and Krüppel-like factor 3 (BKLF)). These experiments revealed bands (a), (b) and (c) to be Sp1, Sp3 and KLF3, respectively (Figure 1D). These findings are consistent with the detection of proteins (a), (b) and (c) in both erythroid and non-erythroid nuclear extracts (Figure 1C), since Sp1, Sp3 and BKLF are all widely-expressed TFs and have previously been shown to regulate *α*-globin expression ^10,11^. In contrast, the binding of protein (d) to probe 13/14 was not strongly affected by any of the antibodies tested. Taken together, these experiments have revealed an erythroid-enriched complex at neighbouring sites in the −128/-90 region upstream of HBA genes.

**Figure 3.**
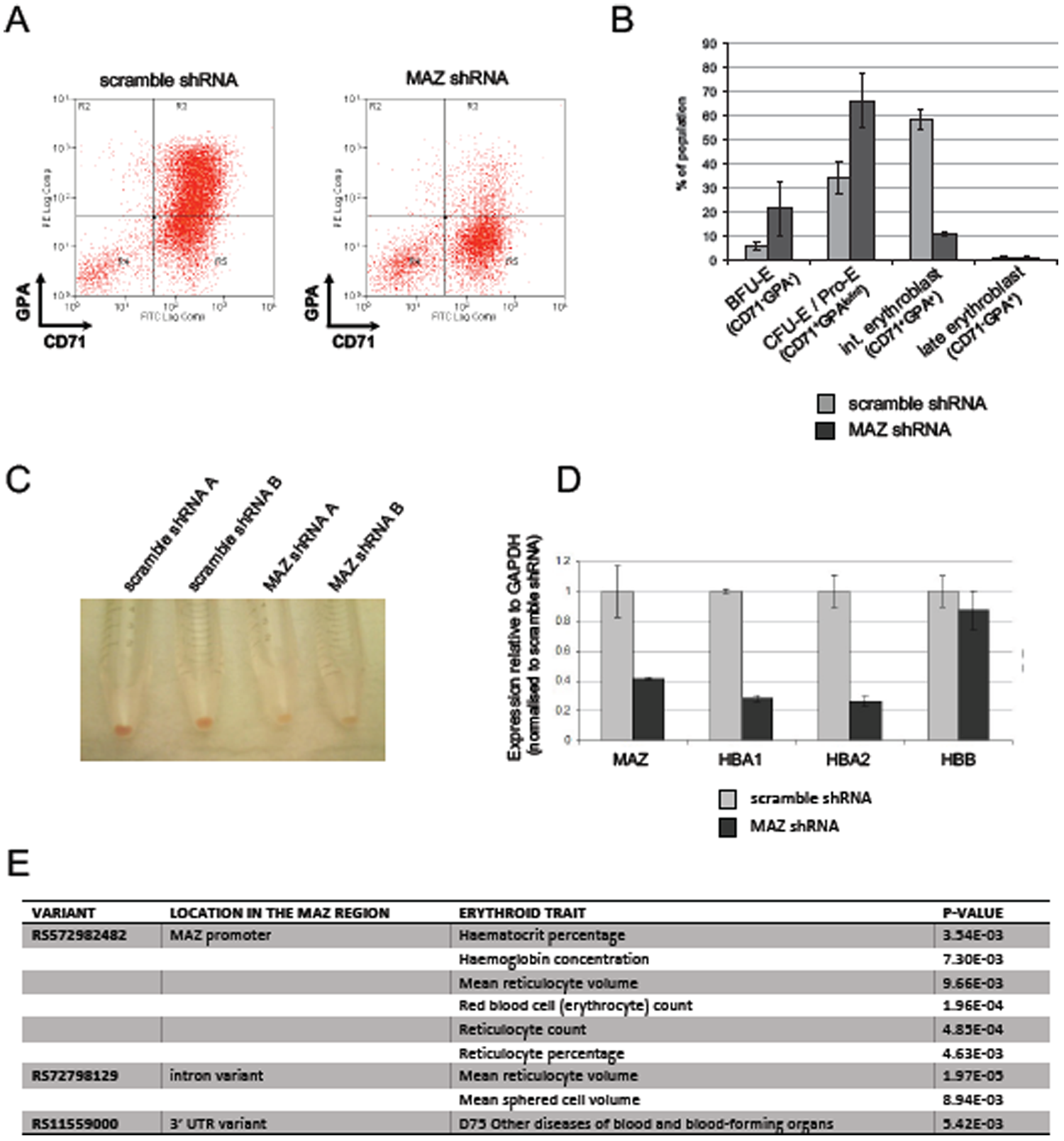
MAZ Knockdown impairs erythropoiesis. **(A)** Flow cytometry analysis of CD71 and GPA expression at day 15 of representative primary erythroblast differentiation cultures infected with lentivirus expressing scramble shRNA or MAZ shRNA. In scramble shRNA cultures, cells progress towards a CD71^+^GPA^+^ phenotype whereas shRNA against MAZ blocked differentiation at CD71 single positive cells. **(B)** Quantitation of flow cytometry staining as demonstrated in (A). Shown is the mean and standard deviation from two independent differentiation cultures. **(C)** Cell pellets from primary cultures at d15 of differentiation. Cultures with the MAZ shRNA are more pale, indicating less extensive hemoglobinisation. **(D)** Real-time RT-PCR analysis of expression of MAZ, HBA1, HBA2 and HBB after 15 d of primary erythroid differentiation cultures. Shown is the mean and standard deviation from two independent differentiation cultures. For each gene, expression (relative to the GAPDH) is normalized to the mean value observed with scramble shRNA. **(E)** Variants around the MAZ locus significantly associated with clinical erythroid traits (p-value <10^−2^).

### A mass spectrometry-based screen identifies MYC-associated Zn-finger protein MAZ as a factor binding at the *α*-globin promoter

In order to identify the factor responsible for band (d), we carried out a protein affinity purification screen, using probe 13/14 as the bait (Figure 2A). As a negative control, we designed a mutant version of probe 13/14 (M3) in which a central guanidine was changed to thymidine, preventing formation of band (d) while also depleting binding of Sp1, Sp3 and BKLF (Suppl. Figure 3B). To carry out this screen, K562 nuclear extracts were incubated with desthiobiotin-modified DNA probes, protein-DNA complexes were purified using streptavidin beads, and eluted proteins were subjected to mass spectrometry (Figure 2A)^12^. Of the 390 proteins identified (Supplemental Table 1), one of the most interesting candidates to emerge was the Myc-associated Zn-finger protein MAZ, a C2H2-type Zinc finger protein which has been previously shown to bind *in vitro* to a G-rich consensus motif (G_3_AG_3_) ^13-15^ (Figure 2B).

In order to investigate whether MAZ is indeed the protein responsible for band (d), we performed siRNA-mediated knockdown (KD) of MAZ in K562 cells. Depletion of MAZ protein was associated with reduction of band (d) in the gel shift assays (Figure 2C). Conversely, expression of exogenous MAZ protein in COS7 cells was associated with strong enrichment of band (d) (Figure 2D). Binding of the over-expressed MAZ protein resulted in a depletion of the Sp3 gel shift, indicating competition for binding under these *in vitro* conditions. Altogether, these experiments confirm that MAZ is indeed the protein responsible for band (d) and indicate that MAZ binds to neighbouring GC boxes at −100/-83 and −128/-111 within the human *α*-globin promoter *in vitro*.

In order to confirm binding by MAZ at the active HBA promoter *in vivo*, we carried out ChIP-qPCR experiments using a previously described series of qPCR amplicons throughout the HBA locus^10^. Whereas binding was absent or low in non-expressing cells (EBV lymphoblast), strong enrichment of MAZ was observed at the HBA promoter and gene body in cells expressing HBA, with the highest enrichment observed in primary erythroid cells (Figure 2E). In these primary cells, MAZ was also enriched at the important distal regulatory element of the *α*-globin locus (MCS–R2). We further investigated the dynamics of MAZ recruitment to the *α*-globin genes during erythroid maturation in primary erythroblast differentiation cultures (Suppl. Figure 4A-C). During differentiation, expression of the adult globin genes HBB and HBA is strongly upregulated, peaking in intermediate and late stage erythroblasts (Suppl. Figure 4D, see also ^16^). Concomitantly with the strong upregulation of α-globin expression, MAZ was dynamically recruited to the HBA promoter and genes in these cultures (Suppl. Figure 4E). Altogether, our results indicate that MAZ binds to the active human HBA locus both *in vitro* and *in vivo*.

**Figure 4.**
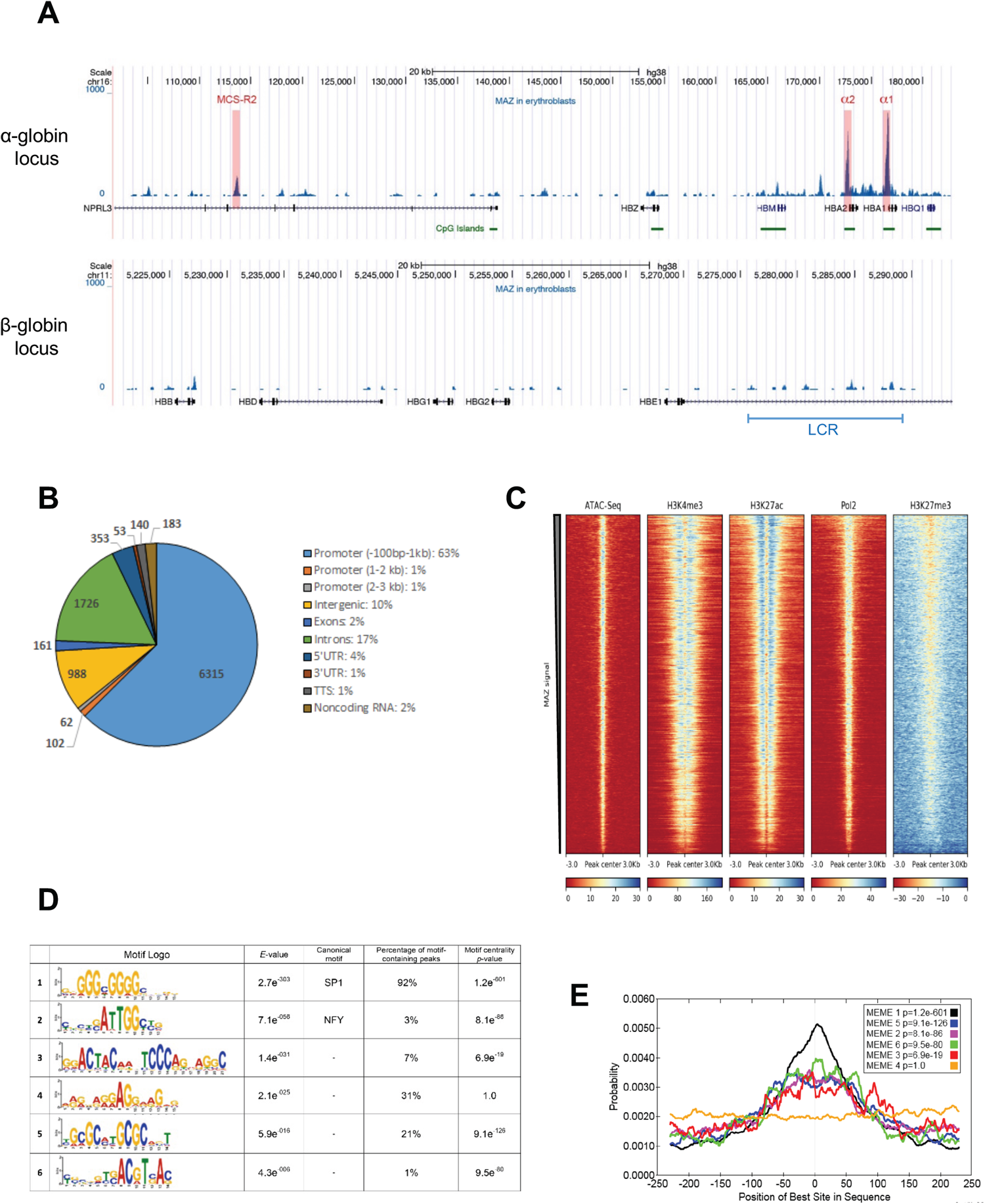
MAZ is enriched at active TSS and binds to DNA through a (G)_3_C(G)_4_ consensus site. **(A)** MAZ ChIP-seq enrichment profiles for the α- and β-globin loci. **(B)** Genomic distribution of MAZ binding sites showing the number of peaks overlapping the category of genomic element indicated. **(C)** Heatmap plots of ATAC-seq, H3K4me3, H3K27ac, Pol II and H3K27me3 signal centred on MAZ peaks in erythroblasts cells (sorted according to decreasing MAZ ChIP signal). **(D)** Characteristics of DNA motifs significantly enriched (E<0.01) in MAZ peaks. **(E)** Motif localisation curves relative to the MAZ peak centres. The numbers of the enriched motifs correspond to the logos in (D).

### MAZ is required for erythroid differentiation and is associated with clinical erythroid-related traits

To investigate the functional significance of MAZ during erythroid differentiation, we used shRNA to deplete MAZ expression in differentiating primary human erythroid cultures. FACS analysis revealed an impaired differentiation of primary erythroid cultures infected with the MAZ shRNA lentivirus, with cells accumulating at the CFUe and pro-erythroblast (CD71^+^GPA^lo^) stages, and failing to differentiate towards intermediate erythroblasts (CD71^+^GPA^+^) (Figure 3A, B). This impaired differentiation could also be observed as a deficiency of haemoglobinisation (Figure 3C). Knockdown of MAZ expression was further associated with downregulation of expression of *α*-globin (Figure 3D). While this downregulation of *α*-globin is likely to also arise indirectly due to the impaired differentiation, expression of *α*-globin was more affected than *β*-globin (where no binding of MAZ was observed in erythroid cells, see below), consistent with MAZ playing a direct regulatory role at the *α*-globin locus.

Given these findings suggesting a previously unknown for MAZ during erythroid differentiation, we further investigated whether human genetic variants in the *MAZ* gene are associated with clinically important erythroid traits. We analysed the GeneATLAS database that reports associations between 778 traits and millions of DNA variants ^17^. Out of 25 erythroid-related traits in the GeneATLAS (Suppl. Table 2), eight traits (32%) were associated with three variants in the MAZ gene or promoter region (rs11559000, rs572982482, rs72798129) with *p*-value 10^−2^-10^−5^ (Figure 3E). Taken together, these findings indicate that MAZ is an important factor in the erythroid differentiation program and suggest that this locus contributes to clinically relevant erythroid traits.

### MAZ occupies TSS and binds directly to DNA through G_3_(C/A)G_4_ motif

Following from our findings at the HBA locus, we expanded our analysis by investigating binding of MAZ genome-wide in primary human erythroid cells by carrying out ChIP-seq (Suppl. Figure 5A-B). As previously observed by ChIP-qPCR (Figure 2E), MAZ was strongly enriched at both HBA2/1 promoters as well as, to a weaker extent, at the MCS-2 remote regulatory element (Figure 4A, top). In contrast, very low binding of MAZ was detected at the promoter and regulatory elements (locus control regions; LCR) of the *β*-globin gene cluster (Figure 4A, bottom). Genome-wide, we identified 10 088 MAZ binding sites in primary erythroid cells (Suppl. Figure 5C). While the majority of MAZ binding sites (65%) were located at promoter regions (Figure 4B), the average MAZ enrichment was comparable for peaks in the promoter regions, intergenic, and genic regions (Suppl. Figure 5D). Overall, MAZ was present at least at one TSS of 28*%* (5466/19646) of protein-coding genes. Comparison with published ChIP- and ATAC-seq datasets from erythroid cells^8,18-20^ (Suppl. Table 6) revealed that MAZ binding sites are located in regions of high chromatin accessibility (as detected by ATAC-seq) that are enriched for activating histone modifications (H3K4me3, H3K27ac) and Pol II but depleted of the repressive histone mark H3K27me3 (Figure 4C). Taken together, these results indicate that MAZ is enriched around TSS of a large proportion of transcriptionally active genes in erythroid cells.

**Figure 5.**
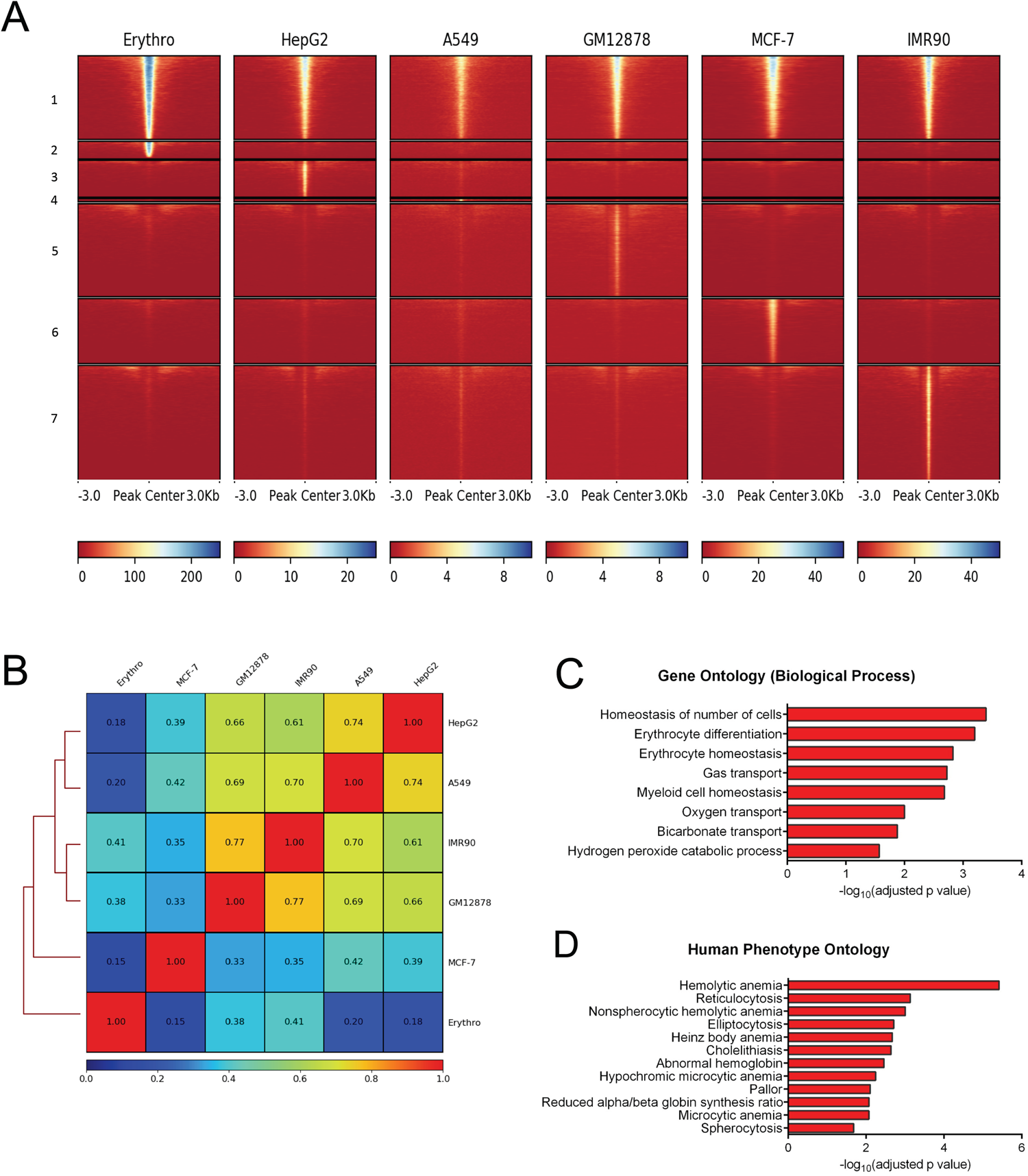
Analysis of erythroid-specific MAZ signal. **(A)** Heatmap plots of MAZ ChIP-seq datasets from six cell types centered on MAZ common and cell-line-specific peaksets. Numbers on left indicate the common peaks (1), and peaks specific to erythroblasts (2), HepG2 (3), A549 (4), GM12878 (5), MCF-7 (6) and IMR90 (7) cells. (**B)** Pearson correlation analysis of MAZ ChIP-seq datasets from the six cell types. **(C)** Gene ontology (by biological function) of genes with erythroid-specific MAZ signal. **(D)** Human Phenotype Ontology of genes with erythroid-specific MAZ signal.

In order to identify motifs contributing directly to MAZ binding, we used the MEME software suite ^21^ to discover *de novo* enriched sequence motifs among a training set consisting of the 500 highest ranked MAZ peaks ^22^. Overall, six enriched motifs were detected in the MAZ erythroblast training dataset (E-value <0.01) (Figure 4D). The most significant motif found, G_3_(C/A)G_4_, is contained within the 13/14 probe derived from the HBA promoter used to identify MAZ in the MS screen, and was similar to the published canonical MAZ motif G_3_AG_3_ ^23^. This canonical motif was present in 92% of all MAZ peaks (see methods and Figure 4D), and was the only enriched motif with a narrow unimodal central enrichment in the peaks and a large maximum site probability (Figure 4E), suggesting that the vast majority of MAZ genomic binding in erythroblasts is due to direct DNA binding of MAZ to this canonical DNA sequence motif. In agreement with these observations, mutations within this core G_3_CG_4_ motif present in the 13/14 probe prevented MAZ binding *in vitro*, while mutations outside this core motif had little effect (Suppl. Figure 6). Furthermore, the 11/12 probe, which exhibits lower affinity for MAZ (Figure 1B and Sup Fig. 2C), contains a less-perfect version of this core motif than the 13/14 probe (Suppl. Figure 6). The second most abundant motif detected in our MAZ training set was a GGAGGA-containing motif. This motif was present in 31% of MAZ peaks, but did not exhibit central localisation, suggesting it may contribute to MAZ binding in a cooperative manner. The enrichment of this motif is consistent with the previous reports demonstrating binding by MAZ to GGA repeats with a high propensity to form G4-quadruplexes ^24-27^.

**Figure 6.**
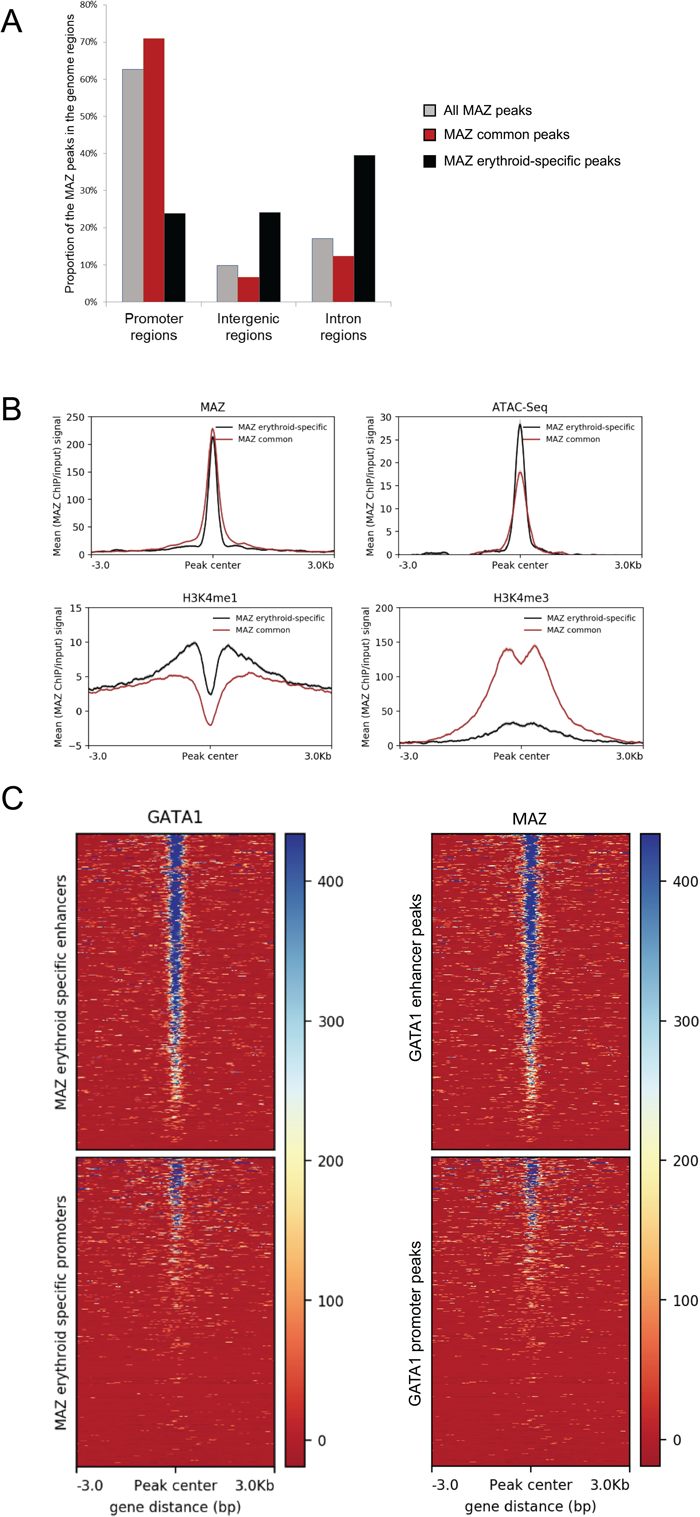
Erythroid-specific MAZ signal is enriched on GATA1-bound enhancers. **(A)** Genome distribution of MAZ common and erythroid-specific binding sites. **(B)** Average H3K4me1, H3K4me3, MAZ and ATAC-seq signal in primary erythroid cells plotted against MAZ erythroid-specific peakset (black) and MAZ common peakset (red) where signal from promoters have been excluded (non-TSS set). **(C)** Left panel: Heatmap plot of GATA1 ChIP-seq centred on MAZ erythroid peaks overlapping enhancers (top) and promoters (bottom). Right panel: Heatmap plot of MAZ ChIP-seq centred on GATA1 peaks overlapping enhancers (top) and promoters (bottom).

### Erythroid-specific MAZ binding sites are associated with the promoters of key erythropoiesis genes

To gain further insight into the specific role of MAZ during erythroid differentiation, we compared the genome-wide profile of MAZ binding in primary erythroblasts with MAZ ChIP-seq profiles from five human non-erythroid cell lines (HepG2, GM12848, MCF-7, IMR90, A549) generated by the ENCODE consortium ^28,29^ (Suppl. Table 6). Comparison of MAZ peaks identified in these six cell types revealed that, as expected for a housekeeping transcription factor, a large proportion of MAZ peaks are shared between at least two different cell types (so called “common” peaks, (n=8308) (Figure 5A). However, between 8 and 40% of the MAZ peaks observed in a given cell line were not observed in any other cell type (Figure 5A, Suppl Table 3), suggesting that as well as regulating housekeeping functions, MAZ also plays an important role in the control of cell-type restricted gene expression programs. Consistent with this, Pearson correlation coefficients for these datasets indicated that MAZ exhibited globally distinct binding profiles in the different cell types (correlation coefficient < 0.85 for all pairwise comparisons) (Figure 5B). In particular, 18% of MAZ peaks observed in erythroid cells were not shared with other cell types (1780 peaks, termed “erythroid-specific peaks”). Comparison with gene expression profiles ^20^ revealed that MAZ binds 41% (218/528) of promoters of genes with erythroid-specific expression. Gene ontology analysis revealed that genes with a TSS bound by MAZ in an erythroid-specific manner were significantly associated with erythroid differentiation and blood haemostasis functions (Figure 5C) as well as phenotypes associated with haematological diseases (Figure 5D). We also investigated MAZ binding at genomic loci which have previously been linked to clinical erythroid phenotypes in GWAS studies. Out of 31 genes associated with erythropoiesis ^30^, MAZ peaks were present on the promoters of 21 of them (Suppl. Table 4). Interestingly, among these genes with promoters bound by MAZ in an erythroid-specific manner were the TFs GATA1 and KLF1, master regulators of erythropoiesis (Suppl. Table 4 and Suppl. Figure 7). Taken together, these findings indicate that MAZ binds to the promoters of genes with key roles in erythroid differentiation and homeostasis.

### Erythroid-specific MAZ signal is enriched on GATA1-bound enhancers

Interestingly, the common and erythroid-specific peaks of MAZ displayed distinct localisation relative to genomic features (Figure 6A). While the majority (71%) of common peaks were localised at promoter regions, only 24% of the MAZ erythroid specific peaks were located at promoters. In contrast to common MAZ peaks, erythroid-specific MAZ peaks were enriched at intergenic (24%) and intronic (40%) sites, suggesting that MAZ may play a particularly important role at erythroid-specific distal regulatory elements. Comparison with our previously published catalogue of erythroid enhancers ^20^ confirmed that 27% of the erythroid-specific MAZ peaks coincided with known enhancer elements, compared with only 7% of the “common” peaks. Consistent with this observation, we observed enrichment of the enhancer-associated histone modification H3K4me1 on erythroid-specific non-TSS MAZ peaks, but not on common non-TSS MAZ peaks, with H3K4me3 following the reverse trend (Figure 6B). Taken together, these findings suggest that erythroid specific binding of MAZ is particularly important at distal enhancer elements.

Ubiquitous and lineage specific factors often work together at proximal and distal regulatory sequences. As such, we explored the association between MAZ and the erythroid master regulator GATA1 in primary erythroid cells. Overall, 48% (854/1778) of erythroid-specific MAZ binding sites overlapped with GATA1 binding sites, consistent with a functional cooperative interaction between these two factors. Interestingly, this co-association between MAZ and GATA was particularly prominent within erythroid enhancer regions, with GATA1 signal being particularly elevated at erythroid MAZ peaks within enhancers and conversely, MAZ signal being stronger at GATA1 enhancer peaks (Figure 6C). Taken together, these findings are consistent with a particular functional cooperativity between MAZ and GATA1 at erythroid enhancer elements.

## DISCUSSION

In this study, we have identified the ubiquitously-expressed Zn-finger protein MAZ as a factor binding to neighbouring GC-rich sites within the human *α*-globin promoters. ChIP and ChIP-seq experiments subsequently confirmed *in vivo* binding of MAZ to the active HBA genes as well as to promoters and enhancers of other key genes within the erythroid differentiation pathway and to genomic loci linked to clinical erythroid phenotypes. Moreover, erythroid-specific binding of MAZ was enriched at distal regulatory enhancer elements, where it frequently co-associated with the erythroid master-regulator GATA1 and knockdown of MAZ impaired erythroid differentiation in *ex vivo* primary cultures. We further showed that genetic variants within the MAZ locus were associated with approximately one third of human phenotypic erythroid traits. Taken together, these findings reveal a previously unrecognised role for MAZ in the erythroid differentiation program.

MAZ has not previously been shown to play an important role in the erythroid lineage. Germline deletion of MAZ in mice results in perinatal lethality ^31^, as might be expected for a ubiquitously expressed TF. MAZ knockdown in a primary erythroid differentiation culture system strongly affects HBA expression whereas expression of HBB was less affected. This is consistent with our ChIP-seq data showing binding of MAZ to the HBA promoters and distal regulatory elements and support a direct role for MAZ at this locus. Since our experiments indicate impaired differentiation of primary erythroid cultures infected with the MAZ shRNA lentivirus, inducible knockdown or knockout approaches will be needed to precisely determine the role of MAZ at specific stages of erythroid maturation.

Through integration of MAZ binding profiles in different cell types, we observed both common and erythroid-specific MAZ binding sites. Erythroid-specific MAZ binding is observed at the promoters of *GATA1* and *KLF1*, encoding master-regulators of the erythroid lineage. MAZ binding is also enriched at distal regulatory elements, which are considered to be primary determinants of tissue-specific gene expression programs^18,32^. Importantly, it has been shown that the activity of erythroid enhancer elements cannot be predicted based on the binding of the master regulators of erythroid differentiation, GATA1 and TAL1, alone ^18^. In contrast, combinatorial co-occupancy of enhancers by both lineage-specific and ubiquitously expressed transcription factors is a more reliable indicator of enhancer activity and cell-specific expression^18,32,33^. Our findings suggest that binding of MAZ together with lineage-restricted factors such as GATA1 might be an important step in the activation or maintenance of many erythroid enhancer elements.

The vast majority (92%) of both shared and erythroid-specific MAZ peaks contain the canonical G_3_CG_4_ motif, indicating that MAZ binds DNA directly in both cases. Consistent with previous reports ^24-27^, our analysis of MAZ whole genome binding also indicated that a likely G4-forming motif (GGAGGA) contributes to MAZ binding, although our data suggests that this motif makes only a secondary contribution to MAZ binding affinity.

At present, it is unclear how binding of MAZ to its target sites is regulated. In particular, while MAZ is a ubiquitously expressed protein whose levels do not change dramatically during erythroid differentiation (data not shown), we observed differential *in vitro* binding by MAZ to the 13/14 probe in gel shift assays, as well as a highly dynamic recruitment of MAZ to the HBA locus in primary erythroid cultures, suggesting that its binding is not primarily regulated at the level of expression. Western blot with α-MAZ antibody detected several bands in K562 cells, which were sensitive to MAZ shRNA (Figure 2C), suggesting the possibility that MAZ is subject to post-translational modifications in these cells. Interestingly, it has been shown in other cell types that phosphorylation at Ser480 (by casein kinase II) ^34^ or at Ser187 and Thr386 (by protein kinase A), increases the DNA-binding and transcriptional activation properties of MAZ^35^. MAZ can also be phosphorylated at Thr71 downstream of the Mitogen-activated protein (MAP) kinase signalling pathway, and this phosphorylation is critical for activation of MAZ in response to inflammatory cytokines^36^. In the future it will be important to characterise how post-translational modifications regulate MAZ activity at the HBA locus and during erythroid differentiation, as well as the upstream signalling pathways involved. Interestingly, our data demonstrate that the sites bound by MAZ within the HBA promoter can also be bound *in vitro* by other KLFs, including SP1, SP3 and BKLF, suggesting the possibility of a crosstalk between these factors in regulating the expression of α-globin. Indeed, complex cross-regulatory interactions involving MAZ and Sp1 have also been reported at other promoters ^37,38^. We also found that these factors compete for the same binding site in our *in vitro* conditions. It is of interest that the expression of SP1 decreases markedly during erythroid differentiation ^39^ and that this decline correlates well with the dynamic recruitment of MAZ to the HBA promoters that is observed in primary erythroid differentiation cultures. These observations suggest that MAZ may bind to the HBA promoters in a commensurate manner when SP1 levels decline during erythroid differentiation, and that this replacement of SP1 by MAZ during the late stages of erythroid differentiation correlates with the maximal activation of the HBA genes. In the future it will be important to explore this potential crosstalk between SP1 and MAZ at the HBA promoters and how it plays out at the genome-wide level during erythroid differentiation.

Overall, this study serves as an example of how a precise molecular characterisation of protein binding at a single regulatory element can be combined with an unbiased mass-spectrometry screen to identify new trans-acting regulators. This approach has revealed the Zn-finger transcription factor MAZ as a previously-unrecognised regulator of the erythroid differentiation program, which may be important for human erythroid phenotypic traits.

## METHODS

### Cell lines and primary erythroid cells

Primary erythroid cells and newly generated EBV-infected B-lymphoblasts were obtained as previously described ^39^. Cell lines (K562, HT29, HeLa, SW13 and COS7) were cultured in RPMI 1640 supplemented with 10% FBS. For siRNA-mediated knockdown of MAZ, K562 cells were transfected with MAZ SMARTPool siRNA or Non-targeting control pool siRNA (Dharmacon) using Lipofectamine according to the manufacturer’s instructions, and cells were harvested after 4 days. For overexpression of MAZ, COS7 cells were transfected with the plasmid pcDSAF-1, which expresses full length MAZ (SAF-1) cDNA under the control of the CMV promoter ^40^. Cells were transfected using Lipofectamine according to the manufacturer’s instructions and were harvested after 30 h.

### Electrophoretic mobility-shift assay (EMSA)

Nuclear extracts were performed as described ^41^ except that nuclei were incubated in buffer C for 30 min. Protein concentration was determined using the Qubit Protein Assay Kit (ThermoFisher). Oligonucleotide probes (Table 1) were designed to have an additional 5’GG overhang on each end after annealing and were labelled by filling in with the large Klenow fragment of DNA Polymerase I in the presence of [α-32P]-dCTP as described by the manufacturer (New England Biolabs). For gel shift reactions, 5 μg of nuclear extract was incubated with 1 ng radiolabelled probe (>10,000 cpm) in buffer [10 mM HEPES (pH 7.8), 50 mM potassium glutamate, 5 mM MgCl_2_, 1 mM EDTA, 1 mM dithiothreitol, 0.5 μg of poly(dI-dC), 10 μg of bovine serum albumin, 1 mM ZnSO_4_, 5% glycerol] for 30 min on ice. For competition experiments, unlabelled competitor oligonucleotides were added to the binding reactions at 100 X molar excess. For gel supershift experiments, antibodies were incubated with nuclear extract for 20 min on ice prior to the addition of the radiolabelled probe. Antibodies used in supershift experiments were as described in previous ChIP studies ^10^. Following binding, samples were subjected to electrophoresis at 4°C for 2.5 h at 12 V/cm on a native polyacrylamide gel (6% [19:1] *bis*:acrylamide in 0.5 X Tris-borate-EDTA). The gels were dried and analysed using a Storm 860 Molecular Imager (Molecular Dynamics Ltd).

### shRNA knockdown in primary erythroid cultures

Lentiviral constructs (pLKO.1) expressing MAZ-specific (clone TRCN0000015343) or non-targeting scrambled shRNAs were obtained from Dharmacon. These constructs express the shRNA from the human U6 promoter and the Puro^r^ gene from the hPGK promoter. Lentiviral preparations were prepared using standard techniques by co-transfection of 293T cells with the lentiviral construct together with pMDLg/pRRE and pMD2.G using Fugene (Roche). Culture supernatants were harvested at 72 h after transfection and concentrated by ultracentrifugation using standard techniques. For lentiviral-mediated knockdown experiments, primary erythroid differentiation cultures were carried out essentially as described in Leberbauer et al., ^42^ with the following modifications. Peripheral blood mononuclear cells were cultured for 8 days in erythroblast expansion medium (StemSpan™ SFEM medium (Stem Cell Technologies) supplemented with Epo (2 U/mL), IGF-1 (40 ng/mL), SCF (100 ng/mL) (all cytokines from Peprotech), dexamethasone (1 μM, Sigma) and cholesterol-rich lipids (40 μg/mL, Sigma)). Cells were infected with concentrated lentivirus for 24 h in wells pre-coated with Retronectin (0.2 mg/ml; Takara) in expansion medium supplemented with protamine sulfate (10 μg/ml). The cells were seeded in fresh complete medium supplemented with cyclosporine A (1 mg/ml) for 24 h prior to selection with puromycin (1 μg/ml) for 48 h. After removing Puromycin, cells were cultured for further 3 days in expansion medium containing cyclosporine prior to harvesting for FACS and RNA (total 15 days in culture).

### Mass Spectrometry

For proteomics analysis, the DNA baits corresponding to oligonucleotide 13/14 WT and 13/14 M3 with a TT/AA-overhang were annealed, phosphorylated, ligated and purified as previously described^12^. The desthiobiotinylated oligonucleotides were coupled to Streptavidin Dynabeads MyOne C1 (Life Technologies) for 60 min at room temperature and excess oligonucleotides removed by washing. The oligonucleotide coupled streptavidin beads were incubated with 250 μg of K562 nuclear extract diluted in PBB buffer (150 mM NaCl, 50 mM Tris–HCl pH 8.0, 10 mM MgCl_2_, 0.5 % Igepal CA630, Complete Protease Inhibitor without EDTA (Roche)) for 2 hours at 4°C under slight agitation. Non-bound proteins were removed by washing three times with PBB buffer and the bound DNA-protein complexes liberated from the streptavidin beads with 200 μl of a 16 mM biotin/50 mM ABC (pH 8.0) solution. The complexes were precipitated with pure ethanol overnight at room temperature. The pellet was resuspended in 50 μl 8M urea and first digested with LysC (Wako) for 3 hours and subsequently diluted to 250 μl with 50mM ABC buffer and digested with 200 ng trypsin overnight, both at room temperature. The tryptic peptides were loaded onto an SCX StageTip^43^ and stored until the MS measurement.

The tryptic peptides were fractionated on an in-house packed 25 cm microcapillary column (Reprosil 3.0 μm; Dr. Maisch GmbH) in a 125 minute gradient from 5 to 60 % acetonitrile. The MS measurement was performed on an LTQ Orbitrap XL (Thermo) with a top5 DDA method using CID fragmentation. The MS data was processed with MaxQuant^44^ version 1.0.12.5 using the human Uniprot database.

### Expression analysis

RNA was extracted with Tri reagent (Sigma) according to the manufacturer’s instructions. RNAs were DNaseI treated (Ambion) and cDNA was generated with SuperScript III (Invitrogen) as previously described ^45^.

### ChIP-qPCR and ChIP-Seq assay

ChIP was performed as previously described ^46^. Briefly, chromatin was first cross-linked with ethylene glycol bis(succinimidyl succinate) (EGS) in PBS at a final concentration of 2 mM for 60 min at RT. Formaldehyde (CH_2_O) was then added at a final concentration of 1 % for 15 min at RT and samples were sonicated over 20 min (10 × 30 s pulses) at 4°C to cleave genomic DNA (Bioruptor, Diagenode). Anti-MAZ antibody was from Bethyl Laboratories (A301-652A). Real-Time PCR was performed using primers and probes (5’FAM-3’TAMRA) for the human α-globin locus described previously ^10^. Each ChIP was performed as two independent experiments and quality was assessed by qPCR. The ChIP-seq libraries were prepared using the New England Biolabs NEBNext® ChIP-Seq Library Prep Reagent Set for Illumina according to the manufacturer’s protocol and starting with between 6 and 40 ng of captured DNA. All libraries received 12 cycles of PCR amplification in the final step before PCR clean-up using Ampure beads. Sequencing was carried out on the HiSeq 2500 using Illumina HiSeq Rapid Cluster Kit v2 - Paired-End for 100 cycles. RTA software version used was Illumina RTA 1.17.20 and the Pipeline software version was bcl2fastq-1.8.3.

### ChIP-Seq analysis

#### Alignment

Reads were aligned using the hisat2 aligner ^47^ version 2.0.3 with the --no-spliced-alignment option, but otherwise default parameters, to a splicing-unaware index of the human GRCh38 genome.

#### The strength of enrichment in the IP

was assessed plotting “fingerprints” using deeptools ^48^ and by calculating normalized strand cross-correlation coefficient (NSC) and relative strand cross-correlation coefficient (RSC) metrics using Phantompeakqualtools ^49^.

#### Normalisation of the ChIP-Seq signal

Reads of the ChIP-Seq samples were first normalised to the input and then scaled to 1x sequencing depth using deepTools v. 3.1.3 ^50^.

#### Peak calling

Peak calling was performed using MACS2 2.1.1 with the minimum FDR (q-value) cutoff of 0.01 ^51^. The top 75% fold enriched peaks were selected for further analysis. For ENCODE datasets optimal IDR thresholded peaks provided by the ENCODE consortium have been used.

#### Peak annotation

was made with UCSC RefSeq gene annotation (GRCh38 genome version) using HOMER suite 4.8 ^52^. For these analyses we define promoters as regions located within a distance of −3000 bp upstream and 100 bp downstream of the TSS.

#### Coverage analysis and ChIP heatmap plots and profiles

were performed using DeepTools suites. General genome arithmetics was performed using BEDTools v. 2.27.1 ^53^. The set of erythroid-specific and housekeeping genes and erythroid enhancers was based on ^20^.

#### Correlation of the datasets

A clustered heatmap of correlation coefficients of bigwig signal was computed using the Pearson method; the bam signal combined from the replicates was plotted over 10 kb bins with DeepTools.

#### Defining erythroid-specific MAZ signal

The peaks called on MAZ datasets were overlapped with the peaks from five MAZ ENCODE datasets with the minimum overlap of 1 bp using BEDTools. The MAZ peaks without the overlap in any of the regions were assigned to “erythroid-specific peakset”.

#### Motif Analysis

To find the sequence motifs enriched in MAZ peaks, genomic sequences (from −50 bp to +50 bp around the centres of the top 500 MAZ peaks ranked on their q-values) were extracted (hg38) and used as input for MEME *de novo* motif discovery ^54^ with *E*-value cutoff 0.01 and motif size defined as 6-30 bp.

To calculate the presence of the motifs in a given peak, we used FIMO with the above defined motifs at a *p*-value threshold of 10−4 ^55^. Motif central enrichment analysis was performed using CentriMo ^22^ on regions from −250bp to +250 bp relative to the peak summits.

#### Functional analysis of genes with erythroid-specific MAZ enrichment

g:Profiler ^56^ was employed to conduct Gene Ontology (GO) Biological Function and HP (Human Phenotype) gene annotations. Fisher’s exact test was used to retrieve significantly enriched GO terms for genes marked with erythroid-specific MAZ signal. Functional categories are defined as those containing at least five genes and a minimum enrichment score of 1.3 (*p*-value < 0.05). GeneATLAS database ^17^ was used for mining MAZ variants associated with erythroid-specific traits (p-value < 10^−2^).

## Supporting information

Deen et al Suppl material

## Data sources, protocols and analysis

Sources of ChIP-seq data are shown in Suppl. Table 6.

## DECLARATIONS

### Authors’ contributions

D.G. and D.V. developed the hypothesis; F.B., M.L.H., V.S., J.A.S.S., H.A., and D.G. performed experiments and/or collected data; D.D., F.B., M.M. D.G. and D.V. analyzed data; interpreted data; and D.D., D.G., and D.V. wrote the manuscript, which was revised and approved by all authors. All authors read and approved the final manuscript.

## Acknowledgements

We would like to thank Doug Higgs, Robert Beagrie, Michele Goodhardt and Philipp Voigt for critically reading the manuscript. High-throughput sequencing was provided by Edinburgh Genomics (http://genomics.ed.ac.uk).

## Ethics approval and consent to participate

Study protocols received ethical approval from Oxford University Ethical Review Panel. The data were analysed anonymously.

