## Supplementary material for "Identification of MAZ as a novel transcription factor regulating erythropoiesis": Deen et al Suppl material

### ADDITIONAL MATERIAL

**Suppl Fig 1. Erythroid-specific hypersensitivity regions DNAse-Seq. (A)** Structure of the α-globin locus. Chromosomal position (hg38 genome build) is shown above the genes. The locus consists of the embryonic ζ gene (HBZ), the duplicated foetal/adult α genes (HBA2 and HBA1) together with two flanking pseudogenes (HBM and HBQ1). The upstream, widely-expressed gene, NPRL3 is transcribed from the opposite strand to that of the HBA genes and contains the remote regulatory elements of the α-globin locus (MCS -R1 to -R3) (indicated by red dots). Below is shown the ATAC-seq signal in human erythroblasts (taken from [8]) indicating open chromatin structure at promoter and distal regulatory elements (shaded regions). **(B)** Fine-mapping of the promoter region of the HBA genes. Nuclei from EBV and K562 were exposed to the restriction enzymes *Bfa*I, *BsrF*I, *Fok*I, *Hinf*I, *Eag*I, *Nco*I, *Mse*I and *Hind*III as indicated. DNA was subsequently purified, limit-digested with *Pst*I and analysed by Southern blot using the 3' *Hind*III/*Pst*I segment of the HBA1 gene as a probe (blue rectangle).

**Suppl Figure 2. Both probes 11/12 and 13/14 bind the (d) complex. (A)** Comparison between probe 11/12 and probe 13/14. The predicted core MAZ-binding motif is highlighted. **(B)** Gel-shift assays showing that both probes bind the (d) complex. The affinity of the (d) complex is stronger for the 13/14 probed as demonstrated by using unlabelled oligonucleotides as competitor (100-fold molar excess over radiolabelled probe).

**Suppl Figure 3. Characterisation of DNA binding of unknown protein by EMSA. (A)** Increased concentration of EDTA abolished the protein/DNA complex for bands (a), (b), (c), and (d) with the probe 13/14, but not for the probe 7/8, which is bound by a non-Zinc finger protein, CBFβ (see Figure 1B). **(B)** A single nucleotide mutation (G->T) abolishes binding of the (d) complex to probe 13/14. The sequence of the wild type and mutant (M3) probes are shown above, with the mutated nucleotide indicated in red. EMSA were carried out using K562 nuclear extracts and either wild type or mutant probes together with unlabelled competitor oligonucleotides (100-fold molar excess) as indicated.

**Suppl Figure 4. Dynamics of MAZ recruitment during erythroid differentiation. (A)** Diagram representing the expression of the transferrin receptor (CD71) and glycophorin A (GPA) during erythroblast differentiation in Phase II culture conditions. **(B)** Flow cytometry analysis of CD71 and GPA during Phase II of a representative primary erythroblast differentiation culture. Cells initially acquire expression of CD71 alone (d6) before becoming double positive for both CD71 and GPA (d9-12). At later stages in culture, expression of CD71 decreases (d15). **(C)** Quantitation of flow cytometry staining as demonstrated in (B). Shown is the mean and standard deviation from two independent differentiation cultures. **(D)** Real-time RT-PCR analysis of expression of α- (HBA2) and β- (HBB) globin at the indicated time points during Phase II of primary erythroid differentiation cultures. Expression (relative to GAPDH) shows the mean and range from two independent differentiation cultures. **(E)** ChIP-qPCR analysis of MAZ binding at the α-globin genes at the indicated time points during Phase II of primary erythroid differentiation cultures. The y axis represents enrichment over the input DNA, normalised to a control sequence in the human 18S gene. The x axis indicates the Taqman probes used. The positions of probes at the HBA Promoter/Exon1 and HBA Ex2, and probe 82 (used as a negative control) are indicated in Figure 2E. The promoter of the MYC gene, which has been previously shown to be bound by MAZ [57] was used as a positive control for the ChIP.

**Suppl Figure 5. MAZ ChIP-Seq quality control metrics. (A)** Fingerprint of MAZ ChIP-Seq dataset compared to input showing an enrichment of DNA fragments for MAZ. **(B)** Cross-correlation plot profile for MAZ ChIP-Seq dataset indicating the normalized strand coefficient (NSC) and the ratio between the fragment-length peak and the read-length peak (relative strand correlation, RSC) for assessing signal-to-noise ratios. **(C)** Distribution of MAZ ChIP-seq signal relative to the called MAZ peaks. **(D)** The average relative MAZ signal around the MAZ peaks located at the promoters, intergenic and genic regions.

**Suppl Figure 6. DNA binding specificities of MAZ.**

EMSA assays were carried out with the indicated probes and K562 nuclear extract. The consensus $G_3CG_4$ MAZ-binding motif is shown in bold. Variation from the wild type 13/14 probe in mutant probes is indicated in red. Variation from the $G_3CG_4$ consensus in probe 11/12 is indicated in blue.

**Suppl. Figure 7. MAZ occupies loci coding for erythroid transcription factors GATA1 and KLF1.** MAZ ChIP-seq enrichment profile at GATA1 and KLF1.

**Table S1. Results of affinity-purification screen.** Detected proteins with their gene names, description and unique identifiers (Uniprot and Ensembl). Pfam annotations (protein domains) were mapped to the protein entries. Table contains identified number of MS/MS spectra for proteins bound to WT and the M3 oligonucleotide (from two independent replicate experiments). Ratio for enrichment between the two oligonucleotide baits is calculated based on protein intensity and significance from MaxQuant analysis [44].

**Table S2.** Erythroid-related traits used to search in the GeneATLAS database for MAZ variants.

**Table S3.** Tissue specific MAZ peaks in ENCODE human cell lines and human primary erythroid cells.

**Table S4.** Location of MAZ peaks within genomic loci associated with erythroid traits and erythropoiesis.

**Table S5.** Sequences of oligonucleotides used in this study

**Table S6.** Datasets used in the study

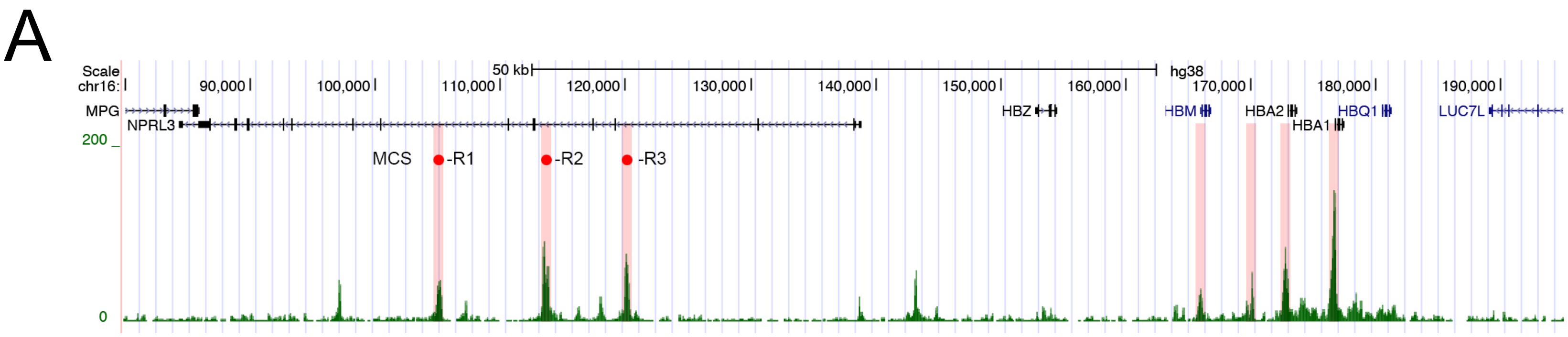
A
Scale
chr16:
50 kb
hg38
90,000
100,000
110,000
120,000
130,000
140,000
150,000
160,000
170,000
180,000
190,000
MPG
NPRL3
HBZ
HBM
HBA2
HBA1
HBQ1
LUC7L
200
0
MCS
-R1
-R2
-R3

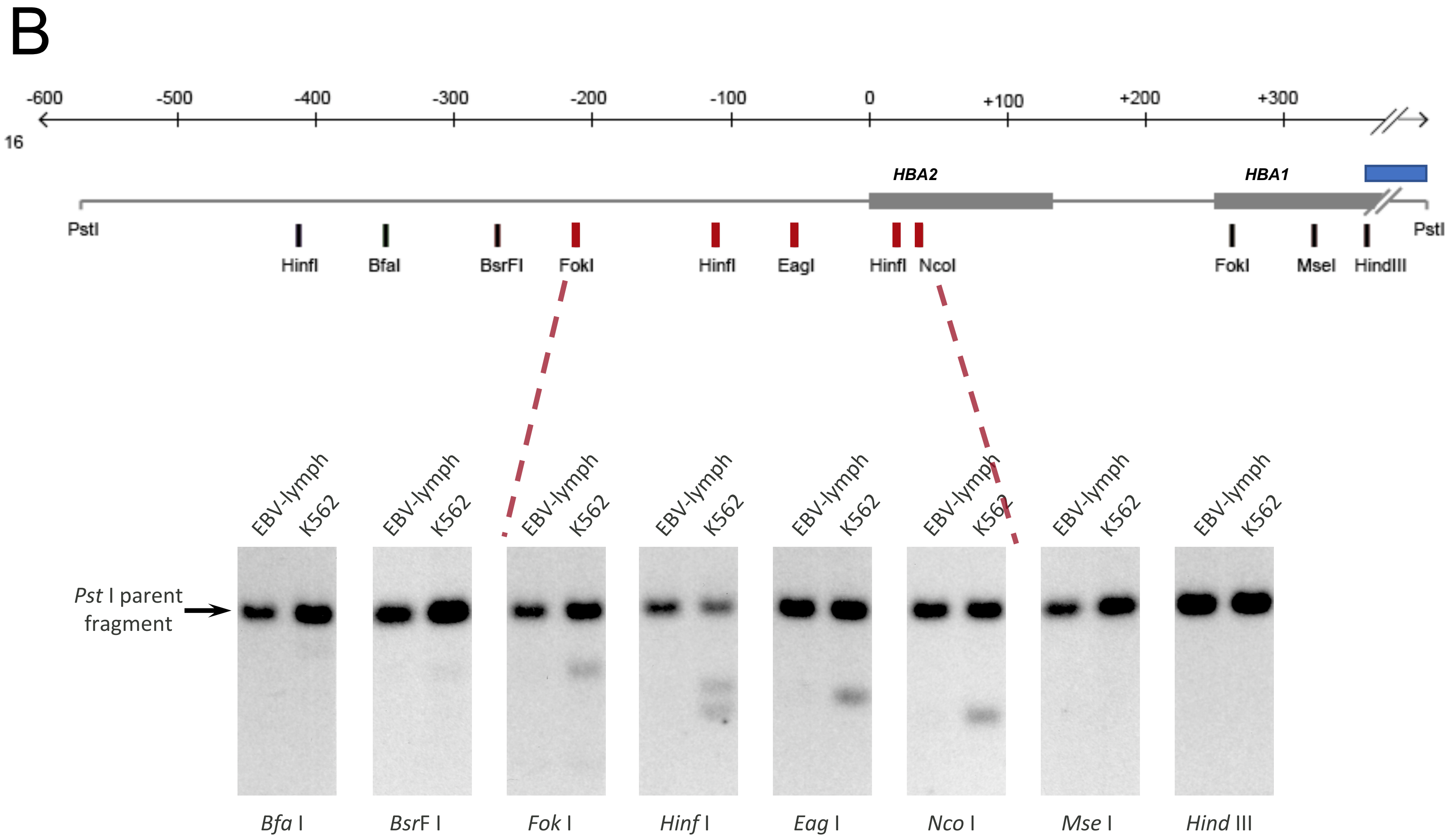
B
-600
-500
-400
-300
-200
-100
0
+100
+200
+300
16
HBA2
HBA1
PstI
HinfI
BfaI
BsrFI
FokI
HinfI
EagI
HinfI
NcoI
FokI
MseI
HindIII
PstI
EBV-lymph
K562
Pst I parent fragment
Bfa I
BsrF I
Fok I
Hinf I
Eag I
Nco I
Mse I
Hind III

Suppl Figure 2

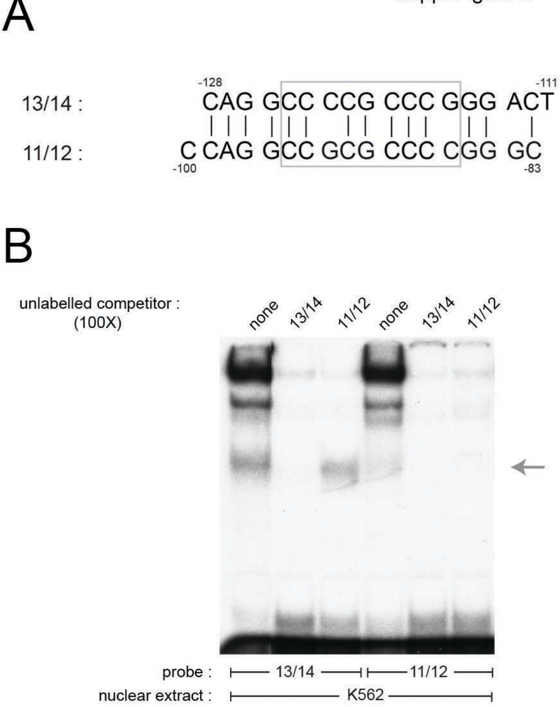

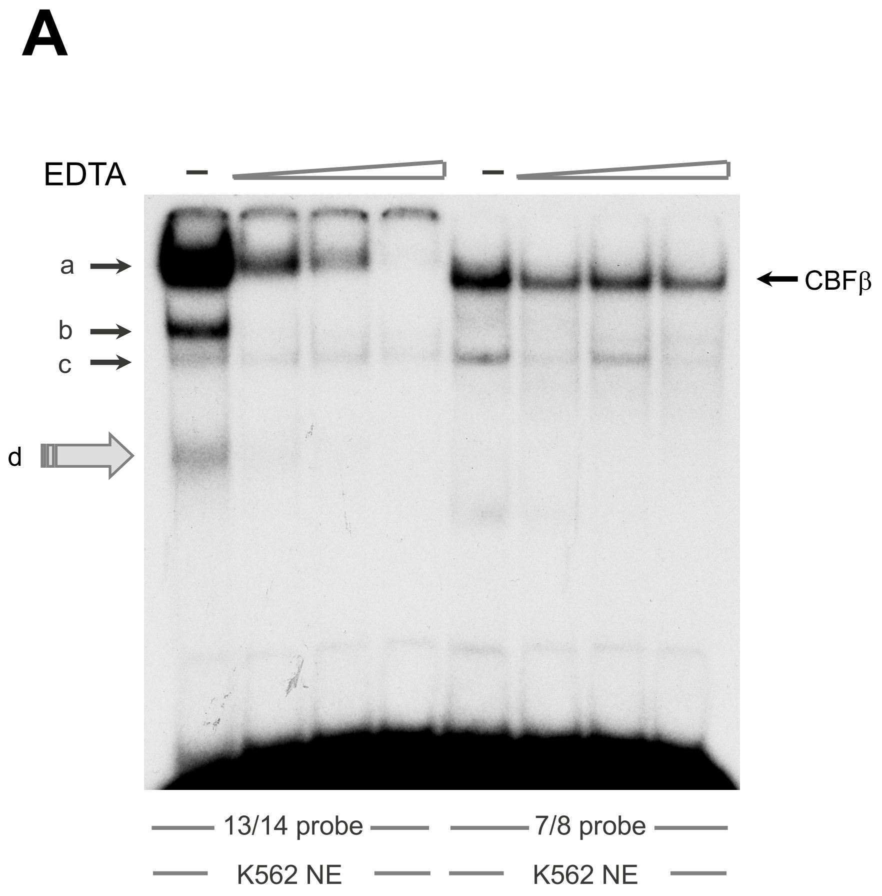
A
EDTA
a
b
c
d
CBFβ
13/14 probe
7/8 probe
K562 NE
K562 NE

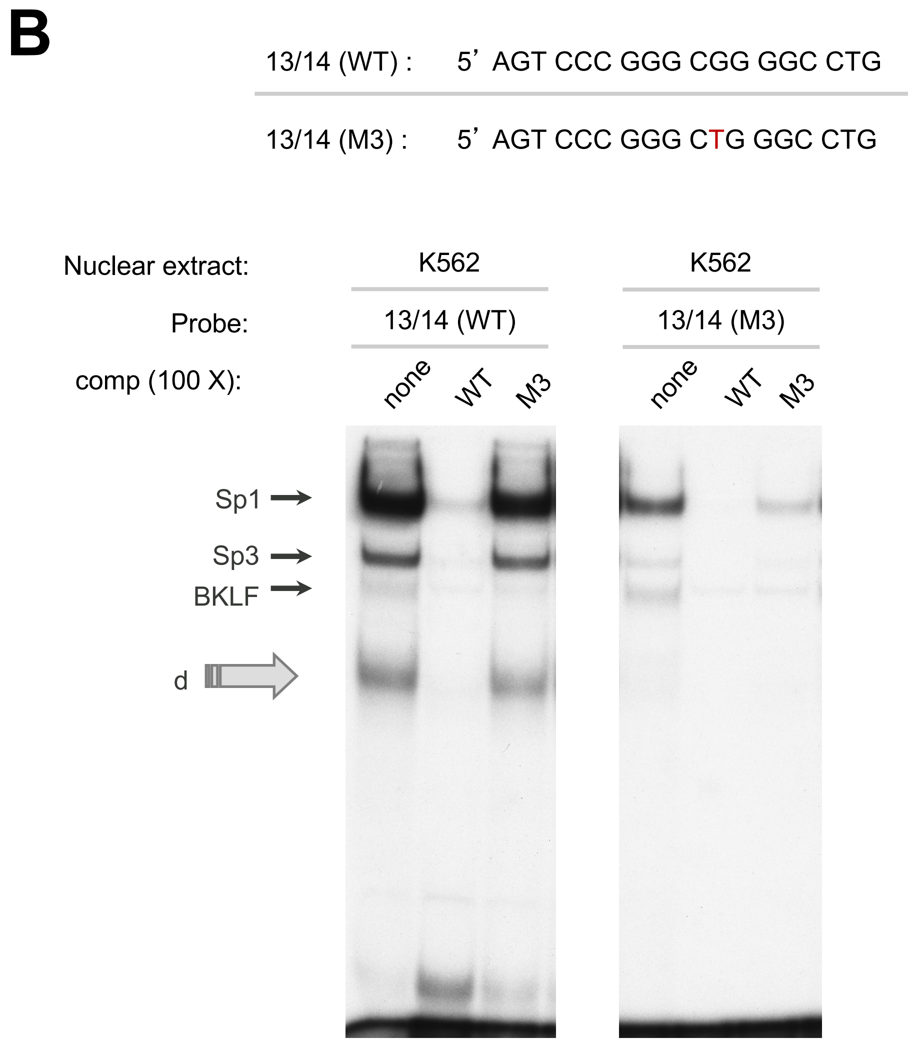
B
13/14 (WT) : 5' AGT CCC GGG CGG GGC CTG
13/14 (M3) : 5' AGT CCC GGG CTG GGC CTG
Nuclear extract:
K562
K562
Probe:
13/14 (WT)
13/14 (M3)
comp (100 X):
none
WT
M3
none
WT
M3
Sp1
Sp3
BKLF
d

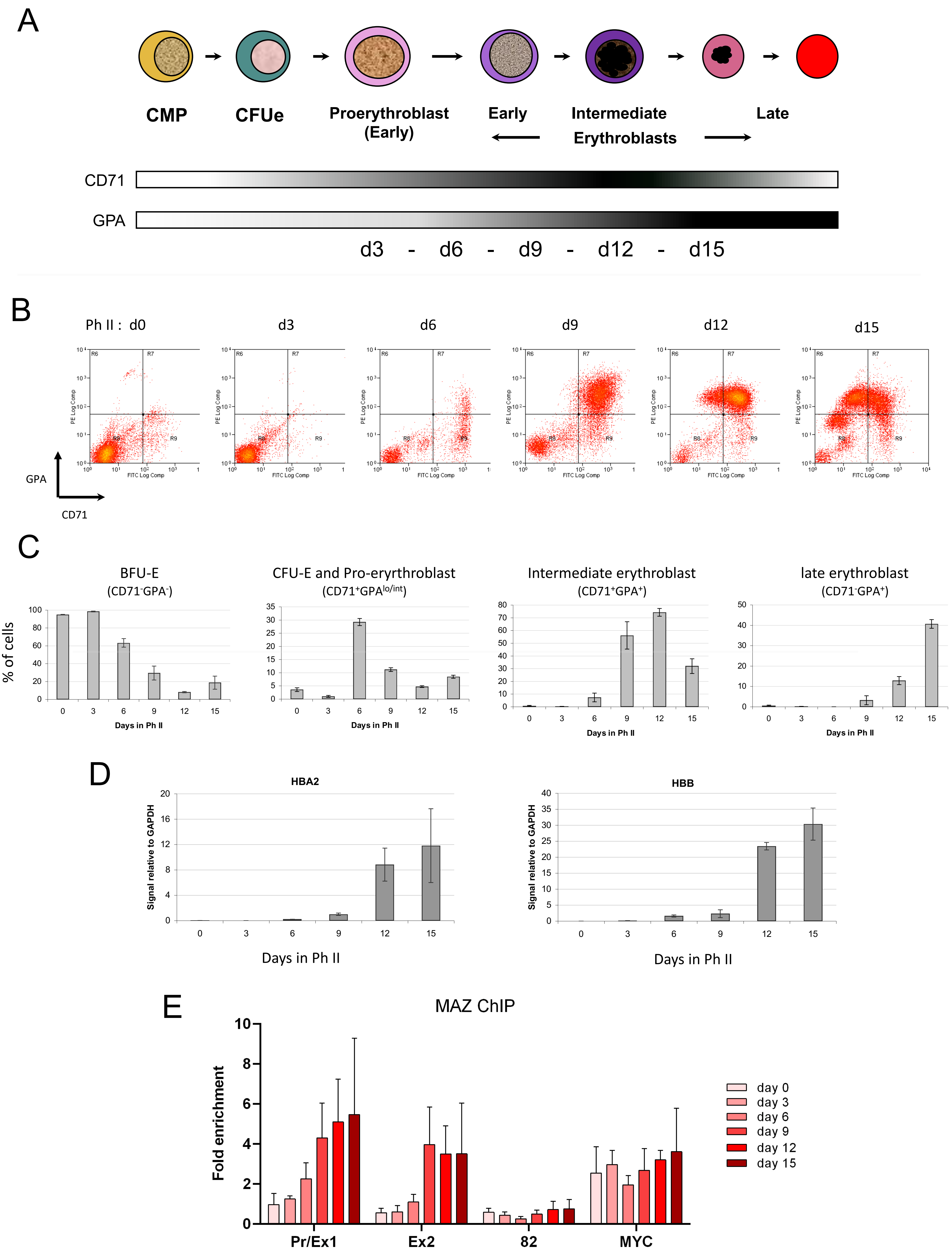
A
CMP
CFUe
Proerythroblast
(Early)
Early
Intermediate
Erythroblasts
Late
CD71
GPA
d3 - d6 - d9 - d12 - d15
B
Ph II : d0
d3
d6
d9
d12
d15
GPA
CD71
C
% of cells
BFU-E
(CD71-GPA-)
CFU-E and Pro-eryrthroblast
(CD71+GPAlo/int)
Intermediate erythroblast
(CD71+GPA+)
late erythroblast
(CD71-GPA+)
Days in Ph II
D
HBA2
HBB
Signal relative to GAPDH
Days in Ph II
E
MAZ ChIP
Fold enrichment
Pr/Ex1
Ex2
82
MYC
day 0
day 3
day 6
day 9
day 12
day 15

Deen et al.,

Suppl Figure 5

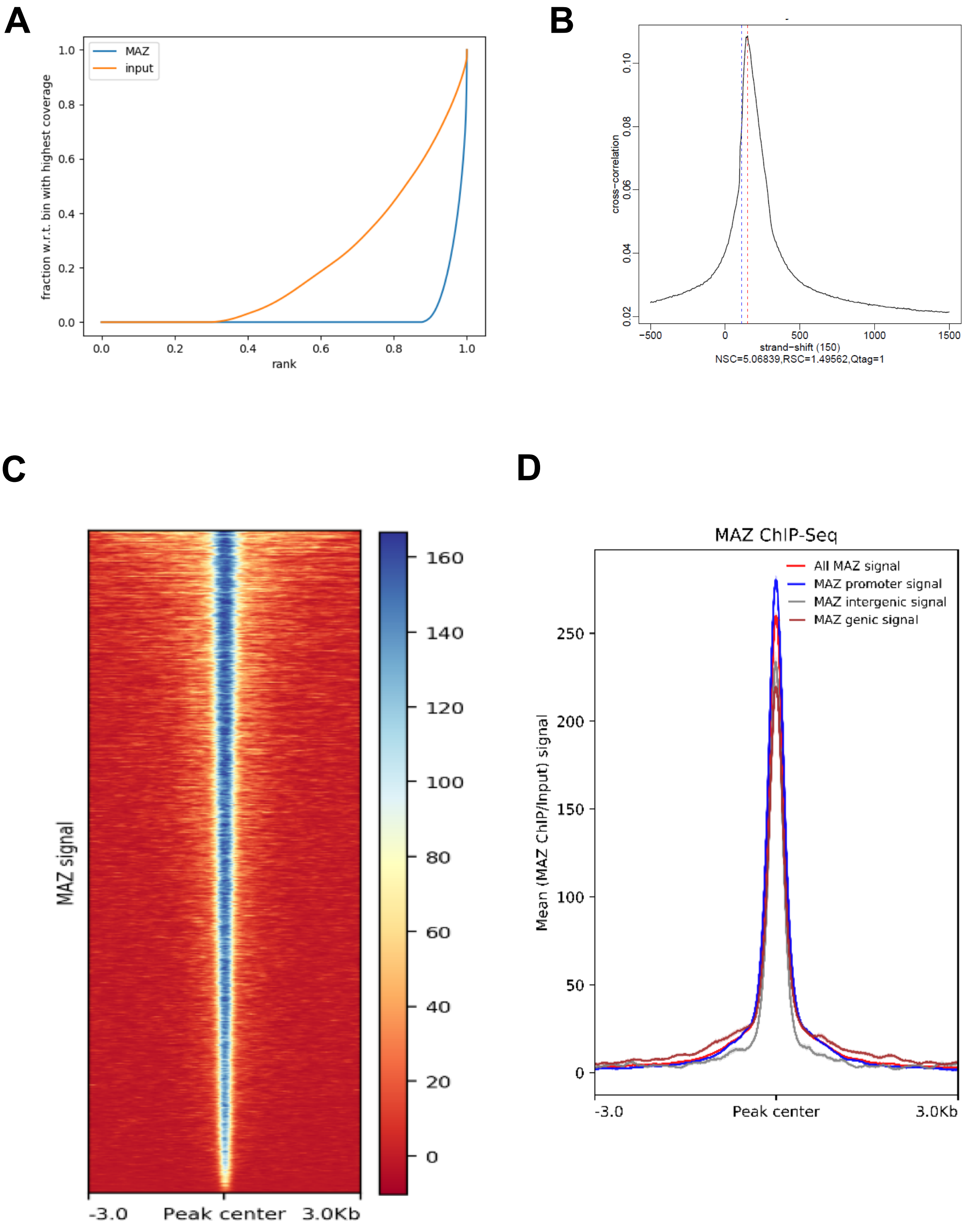

Deen et al.,

Suppl Figure 6

| Oligo | Sequence | Binding of MAZ |
|---|---|---|
| 13/14 (WT) : | 5' AGT CCC **GGG CGG GG**C CTG | + |
| 13/14 (M1) : | 5' CAC ACC **GGG CGG GG**C CTG | + |
| 13/14 (M2) : | 5' AGT CCC **GGG CG**T ATA CTG | - |
| 13/14 (M3) : | 5' AGT CCC **GGG C**T**G GG**C CTG | - (see also Sup. Fig.3B) |
| 13/14 (M4) : | 5' AGT CCC **G**T**G CGG GG**C CTG | - |
| 11/12 (WT): | 5' GC CCG **GGG CG**C **GG**C CTG | < |

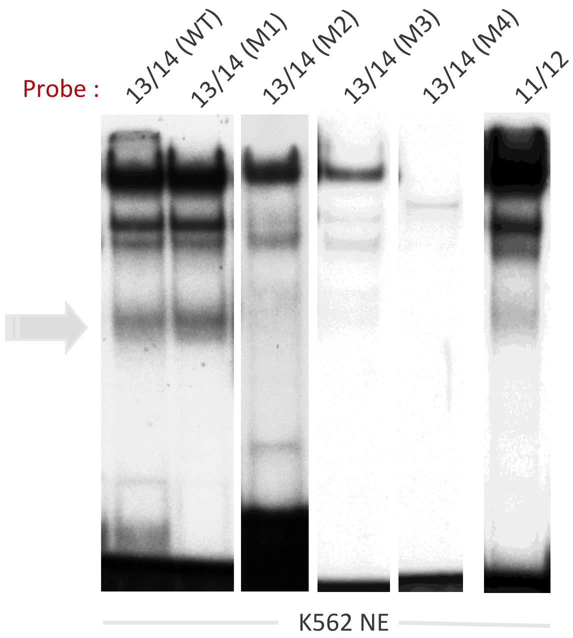

Suppl Figure 7

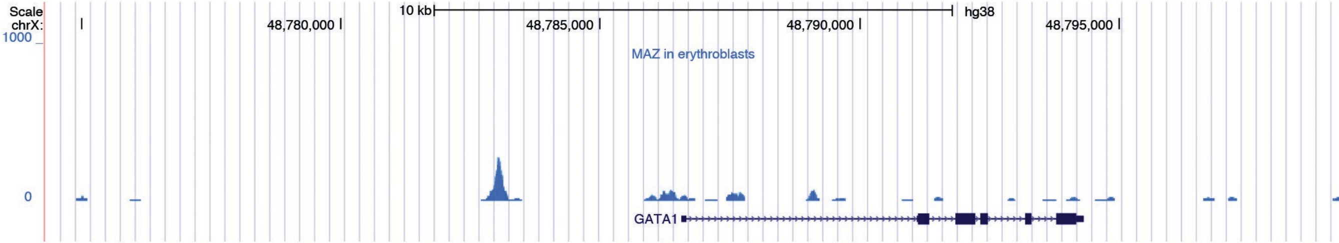


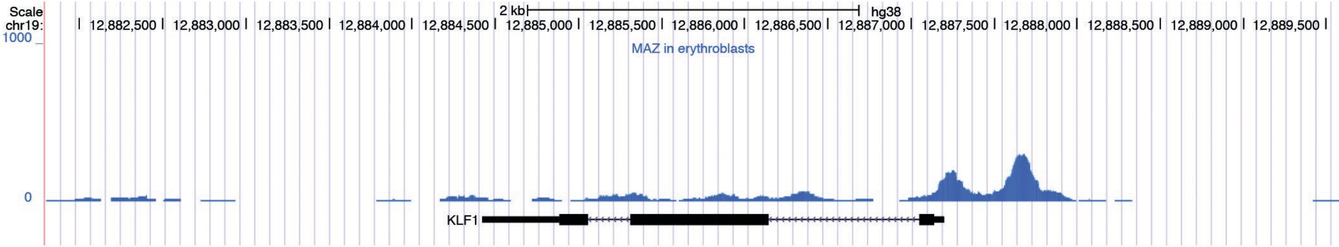

| Gene Names | Protein Names | Mol. Weight [kDa] | Uniprot | Pfam | Pfam Descriptions | ENSEMBL | Chrom | Strand | Position | MS/MS Count neg1 | MS/MS Count neg2 | MS/MS Count pos1 | MS/MS Count pos2 | NEG/POS | POS/NEG | NEG/POS Significance Down |
|---|---|---|---|---|---|---|---|---|---|---|---|---|---|---|---|---|
| FAM139A;FLJ40722;tcag7.730;FLJ00264 | Uncharacterized protein FAM139A;Family with sequence similarity 139, member A;Putative | 100.9 | A6NFQ2;Q17RQ4;Q14D25;Q8IWQ0;Q8N1I1;A6NGZ5;Q8NF84 | | | ENSG00000170379;ENSG0 | 7 | + | 143028667 | 0 | 0 | 0 | 1 | 0.034891007 | 28.66067999 | 1.28E-08 |
| GNL3;E2IG3;NS | Guanine nucleotide-binding protein-like 3;Nucleolar GTP-binding protein 3;Nucleostemin;E2 | 62.0 | Q9BVP2;Q5PU80 | PF08701;PF01926 | GNL3L/Grn1 putative GTPase;GTPase of unknown functio | ENSG00000163938 | 3 | + | 52694976 | 0 | 0 | 0 | 7 | 0.03628214 | 27.56177032 | 1.86E-08 |
| ANKK1;PKK2;SGK288 | Ankyrin repeat and protein kinase domain-containing protein 1;Protein kinase PKK2;X-kinase | 84.632 | Q8NFD2 | PF00023;PF00069 | Ankyrin repeat;Protein kinase domain | ENSG00000170209 | 11 | + | 112763723 | 0 | 0 | 0 | 1 | 0.046219752 | 21.63577185 | 1.68E-07 |
| ZCCHC17;PS1D;HSPC243;HSPC251;LDC4 | Nucleolar protein of 40 kDa;pNO40;Zinc finger CCHC domain-containing protein 17;Putative | 27.6 | Q9NP64 | PF00575 | S1 RNA binding domain | ENSG00000121766 | 1 | + | 31542429 | 0 | 0 | 0 | 6 | 0.046404896 | 21.54945043 | 1.74E-07 |
| NOL1 | Putative RNA methyltransferase NOL1;Proliferating-cell nucleolar antigen p120;Proliferation | 92.9 | P46087;Q3KQS4;A8MQ38 | PF01189;PF08062 | NOL1/NOP2/sun family;P120R (NUC006) repeat | ENSG00000111641 | 12 | - | 6536290 | 0 | 0 | 0 | 5 | 0.04829629 | 20.70552404 | 2.46E-07 |
| RPS8;OK/SW-cl.83;RP11-269F19.3-003 | 40S ribosomal protein S8;Ribosomal protein S8 | 25.2 | P62241;Q5JR94;Q5JR95;Q9BS10 | PF01201 | Ribosomal protein S8e | ENSG00000142937 | 1 | + | 45013826 | 0 | 0 | 0 | 2 | 0.068477656 | 14.60330352 | 4.30E-06 |
| MAZ | MAZ protein;Myc-associated zinc finger protein;MAZI;Purine-binding transcription factor;Pu | 51.1 | Q8IUI2;Q8NFN7;P56270;Q59GP8;Q9U | PF00096 | Zinc finger, C2H2 type | ENSG00000103495 | 16 | + | 29725356 | 0 | 1 | 10 | 19 | 0.070114301 | 14.26242555 | 5.15E-06 |
| SURF6;SURF-6 | Surfeit locus protein 6 | 41.5 | O75683;Q5T8U1 | PF04935 | Surfeit locus protein 6 | ENSG00000148296 | 9 | - | 135187390 | 0 | 0 | 0 | 12 | 0.071259165 | 14.03328268 | 5.83E-06 |
| HIST1H1D;H1F3 | Histone H1.3;Histone H1c | 22.4 | P16402 | PF00538 | linker histone H1 and H5 family | ENSG00000124575 | 6 | - | 26342475 | 3 | 0 | 1 | 3 | 0.091832125 | 10.88943545 | 3.70E-05 |
| HIST1H1E;H1F4 | Histone H1.4;Histone H1b | 21.9 | P10412;A3R0T7;A3R0T8;Q4VB24 | PF00538 | linker histone H1 and H5 family | ENSG00000168298 | 6 | + | 26264566 | 2 | 1 | 1 | 3 | 0.108359576 | 9.228533738 | 0.000112924 |
| H1FX | Histone H1x | 22.5 | Q92522 | PF00538 | linker histone H1 and H5 family | ENSG00000184897 | 3 | - | 130516794 | 2 | 3 | 4 | 21 | 0.111374125 | 8.978746219 | 0.000134919 |
| RSL1D1;CATX11;CSIG;PBK1;L12 | Ribosomal L1 domain-containing protein 1;Cellular senescence-inhibited gene protein;Prote | 55.0 | O76021;A0PJ87;Q32Q62;A0PJ61 | | | ENSG00000171490 | 16 | - | 11835554 | 1 | 2 | 5 | 18 | 0.113081888 | 8.843149138 | 0.000148787 |
| EBNA1BP2;EBP2 | Probable rRNA-processing protein EBP2;EBNA1-binding protein 2;Nucleolar protein p40 | 34.9 | Q99848;Q6IB29 | PF05890 | Eukaryotic rRNA processing protein EBP2 | ENSG00000117395 | 1 | - | 43402440 | 0 | 1 | 5 | 19 | 0.116081763 | 8.614617608 | 0.000175819 |
| HIST1H1B;H1F5 | Histone H1.5;Histone H1a | 22.6 | P16401 | PF00538 | linker histone H1 and H5 family | ENSG00000184357 | 6 | - | 27942606 | 6 | 5 | 5 | 8 | 0.121727315 | 8.215083005 | 0.00023691 |
| DDX24 | ATP-dependent RNA helicase DDX24;DEAD box protein 24 | 96.3 | Q9GZR7;Q4V9L5;Q59FS7;Q7Z4W0;Q7 | PF00270;PF00271 | DEAD/DEAH box helicase;Helicase conserved C-terminal d | ENSG00000089737 | 14 | - | 93587037 | 2 | 0 | 0 | 2 | 0.124198877 | 8.051602606 | 0.000268312 |
| RBM34;KIAA0117;RP11-739C15.1-008 | RNA-binding protein 34;RNA-binding motif protein 34;Novel protein | 48.6 | P42696;A8K8J7;A2A2V2 | PF00076 | RNA recognition motif. (a.k.a. RRM, RBD, or RNP domain) | ENSG00000188739 | 1 | - | 233361121 | 0 | 4 | 5 | 33 | 0.125066781 | 7.99572832 | 0.000280066 |
| RPL4;RPL1 | 60S ribosomal protein L4;L1 | 47.7 | P36578;A6NIF7;A8K502;Q53G74;Q59 | PF00573 | Ribosomal protein L4/L1 family | ENSG00000174444 | 15 | - | 64578708 | 0 | 0 | 0 | 2 | 0.133683948 | 7.480329626 | 0.000419441 |
| DHX36;DDX36;KIAA1488;MLEL1;RHAU | Probable ATP-dependent RNA helicase DHX36;DEAH box protein 36;MLE-like protein 1;RNA h | 114.8 | Q9H2U1 | PF00270;PF07717;PF04408;PF00271 | DEAD/DEAH box helicase;Domain of unknown function (D | ENSG00000174953 | 3 | - | 155476152 | 6 | 6 | 42 | 55 | 0.134661982 | 7.426000893 | 0.000438084 |
| DDX21 | Nucleolar RNA helicase 2;Nucleolar RNA helicase II;Nucleolar RNA helicase Gu;RH II/Gu;Gu-alp | 87.3 | Q9NR30;Q3SWU7;Q8NI92 | PF00270;PF08152;PF00271 | DEAD/DEAH box helicase;GUCT (NUC152) domain;Helicas | ENSG00000165732 | 10 | + | 70385898 | 2 | 1 | 16 | 50 | 0.136695825 | 7.315512374 | 0.000478862 |
| RRS1;KIAA0112;RRR | Ribosome biogenesis regulatory protein homolog | 41.2 | Q15050 | PF04939 | Ribosome biogenesis regulatory protein (RRS1) | ENSG00000179041 | 8 | + | 67503817 | 0 | 1 | 0 | 11 | 0.15305749 | 6.533492745 | 0.000919631 |
| SEMG2 | Semenogelin-2 precursor;Semenogelin II;SGII | 65.4 | Q02383;A0JLP3;A8K6Z6;Q53ZU2;Q6I | PF05474 | Semenogelin | ENSG00000124157 | 20 | + | 43283424 | 0 | 1 | 0 | 7 | 0.156822854 | 6.376621612 | 0.001053587 |
| 101F10.1 | Putative uncharacterized protein 101F10.1 | 51.6 | O43328;Q1ED39;Q5FWF3 | | | ENSG00000103550 | 16 | - | 19625224 | 2 | 8 | 8 | 44 | 0.157946096 | 6.331273939 | 0.001096196 |
| HLTF;HIP116A;RNF80;SMARCA3;SNF2L3;ZBU1 | cDNA FLJ76830, highly similar to Homo sapiens SWI/SNF related, matrix associated, actin dep | 113.9 | A8K5B6;Q14527;Q05BZ6;Q05DF7;Q5 | PF00271;PF08797;PF00176;PF00097 | Helicase conserved C-terminal domain;HIRAN domain;SN | ENSG00000071794 | 3 | - | 150230594 | 0 | 0 | 2 | 17 | 0.170312698 | 5.871552796 | 0.001653049 |
| RFC1;RFC140 | Replication factor C subunit 1;Replication factor C large subunit;RF-C 140 kDa subunit;Activa | 128.3 | P35251;A8K6E7;Q14756;A6NHS4 | PF00004;PF00533;PF08519 | ATPase family associated with various cellular activities (A | ENSG00000035928 | 4 | - | 38965471 | 0 | 2 | 2 | 35 | 0.173658827 | 5.758417338 | 0.001833606 |
| HIST1H1C;H1F2 | Histone H1.2;Histone H1d | 21.4 | P16403;A8K4I2 | PF00538 | linker histone H1 and H5 family | ENSG00000187837 | 6 | - | 26163994 | 4 | 11 | 13 | 28 | 0.179330879 | 5.576284492 | 0.002171482 |
| NOL5A;NOP56;RP4-686C3.1-010;RP4-686C3.1-016 | Nucleolar protein 5A;Nucleolar protein Nop56;56kDa with KKE/D repeat | 66.0 | O00567;A0PJ92;A8K9K6;Q9BSN3;Q5J | PF01798;PF08156;PF08060 | Putative snoRNA binding domain;NOP5NT (NUC127) dom | ENSG00000101361 | 20 | + | 2581178 | 0 | 0 | 3 | 13 | 0.182715717 | 5.472982937 | 0.002393115 |
| RPS17;LOC392101;hCG_1808625;tcag7.955 | Uncharacterized protein ENSP00000352682;40S ribosomal protein S17;Similar to dJ753D5. | 15.9 | A8MVC5;A6NH77;P08708;A4D1Q6 | PF00833 | Ribosomal S17 | ENSG00000197575;ENSG0 | 5 | + | 116079834 | 1 | 0 | 3 | 2 | 0.186869795 | 5.351319615 | 0.002686553 |
| NKRF;ITBA4;NRF | NF-kappa-B-repressing factor;NFkB-repressing factor;Transcription factor NRF;ITBA4 protein | 77.7 | O15226;A1Z1A7;A2I827;A2I828;A2I8 | PF00035;PF01585;PF01424 | Double-stranded RNA binding motif;G-patch domain;R3H | ENSG00000186416 | X | - | 118606328 | 2 | 1 | 1 | 26 | 0.189789398 | 5.268998224 | 0.002907444 |
| RBM28 | RNA-binding protein 28;RNA-binding motif protein 28 | 85.7 | Q9NW13;A4D100;Q53H65 | PF00076 | RNA recognition motif. (a.k.a. RRM, RBD, or RNP domain) | ENSG00000106344 | 7 | - | 127737673 | 0 | 1 | 1 | 13 | 0.1925073 | 5.194608209 | 0.003124297 |
| RPL31 | 60S ribosomal protein L31;RPL31 protein | 14.5 | P62899;Q76N53;Q6IRZ0;A6NF91 | PF01198 | Ribosomal protein L31e | ENSG00000071082 | 2 | + | 100985183 | 0 | 3 | 4 | 3 | 0.193102044 | 5.178609095 | 0.003173222 |
| SP3 | Transcription factor Sp3;SPR-2 | 81.9 | Q02447;Q59FX5;Q68DP2;Q86TP0;Q8 | PF00096 | Zinc finger, C2H2 type | ENSG00000172845 | 2 | - | 174481505 | 0 | 0 | 2 | 2 | 0 | #DIV/0! | 0.003507122 |
| NOL7;C6orf90;NOP27 | Nucleolar protein 7;Nucleolar protein of 27 kDa | 29.4 | Q9UMY1 | PF08157 | NUC129 domain | ENSG00000137420 | 6 | + | 13723555 | 0 | 0 | 0 | 1 | 0.200360096 | 4.991013773 | 0.003814223 |
| YBX1;NSEP1;YB1 | Nuclease sensitive element-binding protein 1;Y-box-binding protein 1;Y-box transcription fac | 35.9 | P67809;A0JLU4;Q05D43;Q6PKI6;A6N | PF00313 | 'Cold-shock' DNA-binding domain | ENSG00000065978 | 1 | + | 42920659 | 19 | 26 | 59 | 56 | 0.203669463 | 4.909916222 | 0.00413432 |
| HP1BP3;RP5-930J4.3-001;hCG_38870;RP5-930J4.3-005;RP5-930J4.3-002;DKFZp434I0612 | Heterochromatin protein 1, binding protein 3;Heterochromatin protein 1, binding protein 3 | 61.4 | A6NI71;A8K5D7;Q05BI0;Q5SSJ5;Q9UI | PF00538 | linker histone H1 and H5 family | ENSG00000127483 | 1 | - | 20941758 | 1 | 3 | 4 | 15 | 0.20422976 | 4.896446045 | 0.004190292 |
| ZNF512;KIAA1805;DKFZp666L156 | Zinc finger protein 512;Putative uncharacterized protein DKFZp666L156 | 64.6 | Q96ME7;Q53RZ7;Q658M0;Q86XK6 | PF00096 | Zinc finger, C2H2 type | ENSG00000198522 | 2 | + | 27659397 | 0 | 0 | 1 | 5 | 0.207506221 | 4.81913262 | 0.0045281 |
| C3orf37;DC12 | UPF0361 protein DC12 | 42.2 | A6NDM4;Q96FZ2;A6NJR9 | PF02586 | Uncharacterised ACR, COG2135 | ENSG00000183624 | 3 | + | 130480374 | 0 | 0 | 0 | 1 | 0.211952193 | 4.718045064 | 0.005015743 |
| NP60;HIBDL | Nuclear protein NP60;Nuclear protein of 60 kDa;3-hydroxyisobutyrate dehydrogenase-like pr | 60.5 | Q49A26 | PF02178;PF03446;PF00855 | AT hook motif;NAD binding domain of 6-phosphoglucona | ENSG00000140632 | 16 | - | 4793208 | 0 | 0 | 0 | 8 | 0.212985911 | 4.695146238 | 0.005134051 |
| ZNF579 | Zinc finger protein 579 | 60.5 | Q8NAF0 | PF00096 | Zinc finger, C2H2 type | ENSG00000179943 | 19 | - | 60780705 | 1 | 0 | 1 | 1 | 0.217221512 | 4.603595621 | 0.00563865 |
| ZNF646;hCG_18384;KIAA0296 | Zinc finger protein 646;Zinc finger protein 646, isoform CRA_a | 200.8 | Q8IVD8;O15015 | PF00096 | Zinc finger, C2H2 type | ENSG00000167395 | 16 | + | 30993263 | 0 | 0 | 1 | 7 | 0.220472558 | 4.53571188 | 0.006047989 |
| TOP1 | DNA topoisomerase 1;DNA topoisomerase I | 90.7 | P11387;A8KA78;Q6NWZ5;Q6PK95;Q9 | PF01028;PF02919 | Eukaryotic DNA topoisomerase I, catalytic core;Eukaryoti | ENSG00000198900 | 20 | + | 39090876 | 61 | 64 | 124 | 174 | 0.22392167 | 4.465847369 | 0.00650365 |
| hCG_2004593;RPL17 | Uncharacterized protein ENSP00000347049;HCG2004593;60S ribosomal protein L17;L23 | 21.5 | A8MVY1;A8MSU5;A6NNY1;P18621;Q6 | PF00237 | Ribosomal protein L22p/L17e | ENSG00000175333;ENSG0 | 15 | - | 69420551 | 0 | 0 | 0 | 2 | 0.228626112 | 4.373953571 | 0.007161423 |
| HIST2H2AB | Histone H2A type 2-B | 14.0 | Q8IUE6 | PF00125 | Core histone H2A/H2B/H3/H4 | ENSG00000184270 | 1 | - | 148125698 | 0 | 1 | 1 | 0 | 0.230344338 | 4.341326601 | 0.007412284 |
| NUMA1;NUMA;NUMA1 variant protein | Nuclear mitotic apparatus protein 1;NuMA protein;SP-H antigen;NUMA1 variant protein | 238.3 | Q14980;Q9UNL7;Q4LE64;A8K394;Q3SYK8;Q59HB8;Q7Z757;Q96HN5;Q9UPG3 | | | ENSG00000137497 | 11 | - | 71387841 | 0 | 0 | 0 | 3 | 0.237780209 | 4.20556447 | 0.008564947 |
| SPTAN1;SPTA2 | Uncharacterized protein SPTAN1;Spectrin alpha chain, brain;Spectrin, non-erythroid alpha c | 286.0 | A6NG51;A6NN88;Q13813;Q9UG16;Q | PF00036;PF08726;PF00018;PF00435 | EF hand;Ca2+ insensitive EF hand;SH3 domain;Spectrin re | ENSG00000197694 | 9 | + | 130354690 | 1 | 0 | 1 | 3 | 0.24187878 | 4.134302307 | 0.009247783 |
| MKI67IP;NIFK;NOPP34 | MKI67 FHA domain-interacting nucleolar phosphoprotein;Nucleolar protein interacting with | 34.2 | Q9BYG3;A8K788 | PF00076 | RNA recognition motif. (a.k.a. RRM, RBD, or RNP domain) | ENSG00000155438 | 2 | - | 122201027 | 1 | 0 | 0 | 5 | 0.244493798 | 4.090083303 | 0.00970145 |
| RPS25 | 40S ribosomal protein S25 | 13.7 | P62851;A6NL90;A6NP42;A8MW59 | PF03297 | S25 ribosomal protein | ENSG00000118181 | 11 | - | 118391635 | 2 | 1 | 3 | 5 | 0.244717089 | 4.086351314 | 0.009740844 |
| TCOF1 | Treacle protein;Treacher Collins syndrome protein;TCOF1 protein | 155.9 | Q13428;Q59FZ2;Q9UFD4;A0JLU0 | PF03546 | Treacher Collins syndrome protein Treacle | ENSG00000070814 | 5 | + | 149717428 | 0 | 2 | 4 | 18 | 0.248042458 | 4.031567853 | 0.010339812 |
| RPL11;RP11-223J15.3-005;RP11-223J15.3-004 | 60S ribosomal protein L11;CLL-associated antigen KW-12;Ribosomal protein L11 | 20.3 | P62913;Q08ES8;Q5VVD0;Q5VVC8;Q5 | PF00281;PF00673 | Ribosomal protein L5;ribosomal L5P family C-terminus | ENSG00000142676 | 1 | + | 23890881 | 1 | 4 | 1 | 2 | 0.251979501 | 3.968576788 | 0.011079027 |
| BXDC2;BRIX | Brix domain-containing protein 2;Ribosome biogenesis protein Brix | 41.4 | Q8TDN6;A0JLQ5;A8K0P5;Q9NUW4 | PF04427 | Brix domain | ENSG00000113460 | 5 | + | 34951238 | 1 | 1 | 6 | 16 | 0.252218815 | 3.964811268 | 0.011125021 |
| SEMG1;SEMG | Semenogelin-1 precursor;Semenogelin I;SGI;Alpha-inhibin-92;Alpha-inhibin-31;Seminal basi | 52.1 | P04279;Q53ZV1;Q53ZV2;A0JLP2;Q53 | PF05474 | Semenogelin | ENSG00000124233 | 20 | + | 43269088 | 3 | 7 | 0 | 24 | 0.256816024 | 3.893838028 | 0.012032364 |
| PDCD11;KIAA0185 | Protein RRP5 homolog;Programmed cell death protein 11 | 208.7 | Q14690;Q2TA92;Q5W093;Q6P2J3 | PF00575 | S1 RNA binding domain | ENSG00000148843 | 10 | + | 105146402 | 1 | 2 | 8 | 22 | 0.2648962 | 3.775063595 | 0.013738204 |
| MMTAG2;C1orf35 | Multiple myeloma tumor-associated protein 2;hMMTAG2 | 29.4 | Q9BU76 | | | ENSG00000143793 | 1 | - | 226355052 | 0 | 1 | 1 | 3 | 0.275914568 | 3.62431026 | 0.016297093 |
| RPS4X;CCG2;RPS4;SCAR;RPS4Y2;RPS4Y2P | 40S ribosomal protein S4, X isoform;Single copy abundant mRNA protein;SCR10;40S ribosom | 29.6 | P62701;Q53HV1;Q96IR1;Q8TD47;A6N | PF00467;PF00900;PF08071;PF01479 | KOW motif;Ribosomal family S4e;RS4NT (NUC023) doma | ENSG00000198034;ENSG0 | X | - | 71408650 | 0 | 1 | 1 | 3 | 0.277494165 | 3.603679376 | 0.016686269 |
| LYAR;PNAS-5 | Cell growth-regulating nucleolar protein | 43.6 | Q9NX58;A8K3Y5 | PF08790 | LYAR-type C2HC zinc finger | ENSG00000145220 | 4 | - | 4320329 | 25 | 25 | 49 | 83 | 0.279336023 | 3.579917796 | 0.017147192 |
| CCDC137 | Coiled-coil domain-containing protein 137 | 33.2 | Q6PK04 | | | ENSG00000185298 | 17 | + | 77244191 | 3 | 7 | 8 | 20 | 0.283547948 | 3.526740384 | 0.018230194 |
| ISG20L2;HSD38;HSD-38 | Interferon-stimulated 20 kDa exonuclease-like 2 | 39.2 | Q9H9L3 | PF00929 | Exonuclease | ENSG00000143319 | 1 | - | 154959019 | 0 | 0 | 5 | 10 | 0.284404838 | 3.516114584 | 0.018455475 |
| REXO4;PMC2;XPMC2H | RNA exonuclease 4;Exonuclease XPMC2;hPMC2;Prevents mitotic catastrophe 2 protein homo | 46.7 | Q9GZR2 | PF00929 | Exonuclease | ENSG00000148300 | 9 | - | 135261009 | 0 | 0 | 0 | 4 | 0.287660432 | 3.476321003 | 0.019326702 |
| RRP1B;KIAA0179 | RRP1-like protein B | 84.4 | Q14684;Q6PJM8 | PF05997 | Nucleolar protein,Nop52 | ENSG00000160208 | 21 | + | 43903860 | 2 | 6 | 9 | 40 | 0.294208631 | 3.398948552 | 0.021152765 |
| BAG2 | BAG family molecular chaperone regulator 2;Bcl-2-associated athanogene 2;BAG-2 | 23.8 | O95816 | PF02179 | BAG domain | ENSG00000112208 | 6 | + | 57145083 | 0 | 0 | 1 | 1 | 0.295437205 | 3.384814037 | 0.021506377 |
| PURB | Transcriptional activator protein Pur-beta;Purine-rich element-binding protein B | 33.2 | Q96QR8;A4D2L7 | PF04845 | PurA ssDNA and RNA-binding protein | ENSG00000146676 | 7 | - | 44889059 | 3 | 5 | 5 | 9 | 0.296259205 | 3.375422548 | 0.021744913 |
| ZBTB48;HKR3;RP11-58A11.4-004;RP11-58A11.4-005 | Zinc finger and BTB domain-containing protein 48;Krueppel-related zinc finger protein 3;Prot | 77.1 | P10074;A8K291;Q6LCP1;Q5SY20;Q5S | PF00651;PF00096 | BTB/POZ domain;Zinc finger, C2H2 type | ENSG00000204859 | 1 | + | 6562698 | 0 | 0 | 3 | 3 | 0.296456269 | 3.373178796 | 0.021802331 |
| MECP2 | Methyl-CpG-binding protein 2;MeCP-2 protein;MeCP2;Uncharacterized protein MECP2 | 53.3 | P51608;A8K079;A8MW75 | PF02178;PF01429 | AT hook motif;Methyl-CpG binding domain | ENSG00000169057 | X | - | 152940218 | 1 | 5 | 15 | 22 | 0.299994055 | 3.333399388 | 0.022848401 |
| RBMX;HNRPG;RBMXP1;CCBL2;DKFZp547N1117;RBMXL1;RP4-531M19.2-001;hCG_23341;R | Heterogeneous nuclear ribonucleoprotein G;hnRNP G;RNA-binding motif protein, X chromos | 42.3 | P38159;Q8N8Y7;Q2VIN3;Q96E39;Q5T | PF08081;PF00076 | RBM1CTR (NUC064) family;RNA recognition motif. (a.k.a | ENSG00000147274;ENSG0 | X | - | 135783288 | 0 | 0 | 2 | 1 | 0.301443158 | 3.317375011 | 0.023285238 |
| WRN;RECQ3;RECQL2 | Werner syndrome ATP-dependent helicase | 162.5 | Q14191;A1KYY9;Q59F09 | PF01612;PF00270;PF00271;PF00570 | 3'-5' exonuclease;DEAD/DEAH box helicase;Helicase cons | ENSG00000165392 | 8 | + | 31010320 | 6 | 2 | 15 | 16 | 0.304470314 | 3.284392443 | 0.024213478 |
| CSDA;DBPA | DNA-binding protein A;Cold shock domain-containing protein A;Single-strand DNA-binding p | 40.1 | P16989;Q59EB5 | PF00313 | 'Cold-shock' DNA-binding domain | ENSG00000060138 | 12 | - | 10742956 | 1 | 0 | 7 | 7 | 0.320982029 | 3.115439213 | 0.029650515 |
| MLF1IP;CENPU;ICEN24;KLIP1;PBIP1 | Centromere protein U;CENP-U;CENP-U(50);CENP-50;Interphase centromere complex protein | 47.5 | Q71F23;A2RRD9;A5D8X7;A8K8D2;Q09GN1;Q09GN2 | | | ENSG00000151725 | 4 | - | 185852232 | 0 | 1 | 1 | 1 | 0.324959647 | 3.077305163 | 0.031054621 |
| CIR | CBF1-interacting corepressor;Recepin | 52.5 | Q86X95;A0PJI7;A6NFI6;A8K8M4;A6NDY8 | | | ENSG00000138433 | 2 | - | 174921126 | 0 | 0 | 0 | 2 | 0 | #DIV/0! | 0.034830712 |
| RPL10A;NEDD6 | 60S ribosomal protein L10a;CSA-19;Protein NEDD6;Neural precursor cell expressed developm | 25.0 | P62906;Q1JQ76;A6NGU2 | PF00687 | Ribosomal protein L1p/L10e family | ENSG00000198755 | 6 | + | 35544156 | 10 | 4 | 3 | 5 | 0.335999913 | 2.976191243 | 0.035142697 |
| CCDC59;BR22;TAP26;HSPC128 | Thyroid transcription factor 1-associated protein 26;TTF-1-associated protein 26;TTF-1-associ | 28.7 | Q9P031 | | | ENSG00000133773 | 12 | - | 81270761 | 1 | 1 | 1 | 2 | 0.33672906 | 2.969746657 | 0.035422524 |
| UBTF;UBF;UBF1 | Nucleolar transcription factor 1;Upstream-binding factor 1;UBF-1;Autoantigen NOR-90;Ribos | 89.4 | P17480;A8K6R8;A8K962;Q05BZ1;Q9E | PF00505 | HMG (high mobility group) box | ENSG00000108312 | 17 | - | 39637928 | 1 | 0 | 2 | 11 | 0.338603382 | 2.953307774 | 0.036147407 |
| BLM;RECQ2;RECQL3 | Bloom syndrome protein;RecQ protein-like 3;DNA helicase, RecQ-like type 2 | 159.0 | P54132;Q3B7X0 | PF08072;PF00270;PF00271;PF00570 | BDHCT (NUC031) domain;DEAD/DEAH box helicase;Helic | ENSG00000197299 | 15 | + | 89061583 | 12 | 2 | 12 | 24 | 0.350074594 | 2.856534059 | 0.040757417 |
| RPS3A;MFTL;hCG_33299 | 40S ribosomal protein S3a;Uncharacterized protein ENSP00000343748;HCG33299, isoform | 29.9 | P61247;A8K4W0;Q6NXR8;A6NCR2 | PF01015 | Ribosomal S3Ae family | ENSG00000145425;ENSG0 | 4 | + | 152240204 | 6 | 7 | 5 | 6 | 0.353276822 | 2.830641403 | 0.042097196 |
| PARP1;ADPRT;PPOL | Poly [ADP-ribose] polymerase 1;PARP-1;ADPRT;NAD(+) ADP-ribosyltransferase 1;Poly[ADP-rib | 113.1 | P09874;Q05D33;Q5VX86;Q6PJL0;Q9 | PF00533;PF08063;PF00644;PF02877;PF0 | BRCA1 C Terminus (BRCT) domain;PADR1 (NUC008) doma | ENSG00000143799 | 1 | - | 224615016 | 123 | 119 | 200 | 304 | 0.353989237 | 2.824944647 | 0.04239837 |
| CTCF | Transcriptional repressor CTCF;CCCTC-binding factor;CTCFL paralog;11-zinc finger protein;CC | 82.8 | P49711;Q53XI7;Q59EL8 | PF00096 | Zinc finger, C2H2 type | ENSG00000102974 | 16 | + | 66153965 | 0 | 0 | 0 | 5 | 0 | #DIV/0! | 0.046394538 |
| DDX18 | ATP-dependent RNA helicase DDX18;DEAD box protein 18;Myc-regulated DEAD box protein;M | 75.4 | Q9NVP1;Q4ZG72;Q53TI6;Q8N254 | PF00270;PF00271 | DEAD/DEAH box helicase;Helicase conserved C-terminal d | ENSG00000088205 | 2 | + | 118288725 | 13 | 16 | 27 | 34 | 0.3648522 | 2.740835875 | 0.047129787 |
| XRCC1 | DNA-repair protein XRCC1;X-ray repair cross-complementing protein 1 | 69.5 | P18887;Q59HH7;Q6IBS4 | PF00533;PF01834 | BRCA1 C Terminus (BRCT) domain;XRCC1 N terminal dom | ENSG00000073050 | 19 | - | 48739304 | 1 | 2 | 3 | 20 | 0.37026312 | 2.700782085 | 0.049582853 |
| XPC;XPCC | DNA-repair protein complementing XP-C cells;Xeroderma pigmentosum group C-complemen | 106.0 | Q01831;A8K146 | PF03835 | DNA repair protein Rad4 | ENSG00000154767 | 3 | - | 14161651 | 0 | 0 | 0 | 2 | 0 | #DIV/0! | 0.052344994 |
| MKI67 | Antigen KI-67 | 358.7 | P46013 | PF00498;PF08065 | FHA domain;K167R (NUC007) repeat | ENSG00000148773 | 10 | - | 129784913 | 1 | 2 | 2 | 28 | 0.377629469 | 2.64809842 | 0.053023619 |
| RPL7A;SURF-3;SURF3;RP11-244N20.10-005 | 60S ribosomal protein L7a;Surfeit locus protein 3;PLA-X polypeptide;Uncharacterized protei | 30.1 | A6NH86;P62424;Q5T8U4;A6NH90;A8 | PF01248 | Ribosomal protein L7Ae/L30e/S12e/Gadd45 family | ENSG00000148303;ENSG0 | 9 | + | 135204890 | 0 | 0 | 0 | 3 | 0.379401182 | 2.635732434 | 0.053868379 |
| C19orf53;HSPC023 | Leydig cell tumor 10 kDa protein homolog | 10.6 | Q9UNZ5 | | | ENSG00000104979 | 19 | + | 13746257 | 5 | 7 | 9 | 9 | 0.403986203 | 2.475332058 | 0.066258886 |
| ARL6IP4;FLJ00169 | ADP-ribosylation factor-like protein 6-interacting protein 4;ARL-6-interacting protein 4;Aip-4 | 44.9 | Q66PJ3;Q8TEM4 | | | ENSG00000182196 | 12 | + | 122030560 | 0 | 0 | 0 | 8 | 0 | #DIV/0! | 0.066396871 |
| HNRNPA2B1;HNRPA2B1 | cDNA FLJ75945, highly similar to Homo sapiens heterogeneous nuclear ribonucleoprotein A2 | 37.5 | A8K064;P22626 | PF00076 | RNA recognition motif. (a.k.a. RRM, RBD, or RNP domain) | ENSG00000122566 | 7 | - | 26197992 | 0 | 0 | 1 | 2 | 0.405951551 | 2.463348142 | 0.067301725 |
| C16orf75 | OB DNA-binding domain-containing protein C16orf175 | 15.9 | Q96E14 | | | ENSG00000175643 | 16 | + | 11318137 | 7 | 6 | 5 | 11 | 0.411961354 | 2.427412159 | 0.070537011 |
| DDX41;ABS | Probable ATP-dependent RNA helicase DDX41;DEAD box protein 41;DEAD box protein abstra | 69.8 | Q9UJV9;Q53HI2 | PF00270;PF00271;PF00098 | DEAD/DEAH box helicase;Helicase conserved C-terminal d | ENSG00000183258 | 5 | - | 176871185 | 1 | 3 | 10 | 9 | 0.417762371 | 2.393705295 | 0.073725188 |
| XRN2 | 5'-3' exoribonuclease 2;DHM1-like protein;DHP protein | 108.6 | Q9H0D6 | PF03159 | XRN 5'-3' exonuclease N-terminus | ENSG00000088930 | 20 | + | 21231942 | 6 | 9 | 12 | 32 | 0.421146586 | 2.374470155 | 0.075614198 |
| DDX54 | ATP-dependent RNA helicase DDX54;DEAD box protein 54;ATP-dependent RNA helicase DP97 | 98.7 | Q8TDD1 | PF08147;PF00270;PF00271 | DBP10CT (NUC160) domain;DEAD/DEAH box helicase;He | ENSG00000123064 | 12 | - | 112079369 | 0 | 0 | 0 | 4 | 0 | #DIV/0! | 0.075770715 |
| MEN1;SCG2 | Menin;Menin 1243del1 variant | 68.0 | O00255;A5HBD0;A5HBC6;A5HBC7;A5 | PF05053 | Menin | ENSG00000133895 | 11 | - | 64327564 | 0 | 2 | 2 | 1 | 0.441276967 | 2.266150453 | 0.087277082 |
| HSPA1A;HSPA1;HSPA1B;HSP70-1 | Heat shock 70 kDa protein 1;HSP70.1;HSP70-1/HSP70-2;Heat shock 70kDa protein 1A varian | 70.1 | P08107;A8K5I0;Q59EJ3;Q9UQC1 | PF00012 | Hsp70 protein | ENSG00000204388;ENSG0 | 6 | + | 31903667 | 0 | 3 | 3 | 1 | 0.442633136 | 2.259207273 | 0.088088174 |
| WDR76 | WD repeat-containing protein 76;WDR76 protein | 69.8 | Q9H967;A0MNP5;Q05CI4 | PF00400 | WD domain, G-beta repeat | ENSG00000092470 | 15 | + | 41906470 | 0 | 0 | 2 | 6 | 0.443224442 | 2.256193264 | 0.088442792 |
| ZFX;ZFY;ZNF711;CMPX1;ZNF6 | Zinc finger X-chromosomal protein;Zinc finger Y-chromosomal protein;X-linked zinc finger pr | 90.5 | P17010;Q59EB9;Q8NHZ2;Q8NHZ3;Q8 | PF00096;PF04704 | Zinc finger, C2H2 type;Zfx / Zfy transcription activation re | ENSG00000005889;ENSG0 | X | + | 24077783 | 1 | 0 | 2 | 0 | 0.44391744 | 2.252671129 | 0.088859144 |
| RPS19 | 40S ribosomal protein S19 | 16.1 | P39019;Q8WVX7 | PF01090 | Ribosomal protein S19e | ENSG00000105372 | 19 | + | 47055828 | 2 | 4 | 6 | 1 | 0.445476269 | 2.244788487 | 0.089798624 |
| MDK;MK1;NEGF2 | Midkine precursor;MK;Neurite outgrowth-promoting protein;Midgestation and kidney prote | 15.6 | P21741;Q2LEK4 | PF01091;PF05196 | PTN/MK heparin-binding protein family, C-terminal dom | ENSG00000110492 | 11 | + | 46359194 | 0 | 1 | 1 | 2 | 0.45003059 | 2.22207117 | 0.092566493 |
| NOC3L;AD24;C10orf117;FAD24 | Nucleolar complex protein 3 homolog;NOC3 protein homolog;NOC3-like protein;Nucleolar c | 92.5 | Q8WTT2;A8K905;A6NJZ9 | PF03914;PF07540 | CBF/Mak21 family;Nucleolar complex-associated protein | ENSG00000173145 | 10 | - | 96082979 | 0 | 0 | 0 | 2 | 0 | #DIV/0! | 0.093975603 |
| RP9 | Retinitis pigmentosa 9 protein;Pim-1-associated protein;PAP-1 | 26.1 | Q8TA86 | | | ENSG00000164610 | 7 | - | 33100937 | 0 | 0 | 0 | 7 | 0 | #DIV/0! | 0.09496142 |
| POLR3E;KIAA1452 | DNA-directed RNA polymerase III subunit RPC5;RNA polymerase III subunit C5;DNA-directed F | 79.9 | Q9NVU0;Q53GZ2 | PF04801 | Sin-like protein conserved region | ENSG00000058600 | 16 | + | 22216231 | 1 | 1 | 1 | 1 | 0.463532676 | 2.157345214 | 0.100968178 |

| | | | | | | | | | | | | | | | | |
|---|---|---|---|---|---|---|---|---|---|---|---|---|---|---|---|---|
| RPS7 | 40S ribosomal protein S7;Uncharacterized protein ENSP00000352132 | 22.1 | P62081;Q57Z92;A6NE65;A8MX50;A8 | PF01251 | Ribosomal protein S7e | ENSG00000171863;ENSG0 | 2 | + | 3600728 | 2 | 4 | 6 | 6 | 0.471970402 | 2.118776931 | 0.106361001 |
| GPATCH4;GPATC4 | cDNA FLJ77219;G patch domain-containing protein 4;Uncharacterized protein GPATCH4 | 50.4 | A8KAG1;Q5T3I0;A8MRS9 | PF01585 | G-patch domain | ENSG00000160818 | 1 | - | 154830908 | 0 | 0 | 1 | 9 | 0.476424432 | 2.098968762 | 0.109249943 |
| AHCTF1P;AHCTF1;ELYS;TMBS62;MSTP108 | Uncharacterized protein AHCTF1P;AT-hook-containing transcription factor 1;Embryonic larg | 256.3 | A6NP09;A8MSG9;Q8WYP5;A6NGM0 | PF02178 | AT hook motif | ENSG00000153207 | 1 | - | 245069023 | 0 | 0 | 0 | 4 | 0.483110522 | 2.069919727 | 0.113639516 |
| DKFZp686M0665 | Putative uncharacterized protein DKFZp686M0665 | 61.1 | Q7Z335;Q8TBM5;Q9NXV6;Q9NYH0 | | | ENSG00000168564 | 4 | + | 184602794 | 2 | 0 | 2 | 8 | 0.487870245 | 2.049725334 | 0.116801951 |
| RPL23A;hCG_1988529 | Uncharacterized protein RPL23A;Ribosomal protein L23a, isoform CRA_a;Uncharacterized pr | 21.916 | A8MUS3;A8MXA8;A6NGD1;A6NN95;P | PF00276;PF03939 | Ribosomal protein L23;Ribosomal protein L23, N-termin | ENSG00000198242;ENSG0 | 17 | + | 24071127 | 0 | 0 | 0 | 1 | 0 | #DIV/0! | 0.116933838 |
| RPS14;PRO2640 | 40S ribosomal protein S14 | 16.3 | P62263 | PF00411 | Ribosomal protein S11 | ENSG00000164587 | 5 | - | 149802945 | 3 | 2 | 1 | 3 | 0.491104484 | 2.036226574 | 0.118968156 |
| PARP2;ADPRT2;ADPRTL2 | Poly [ADP-ribose] polymerase 2;PARP-2;NAD(+) ADP-ribosyltransferase 2;Poly[ADP-ribose] syn | 66.2 | Q9UGN5;A8K214 | PF00644;PF02877;PF05406 | Poly(ADP-ribose) polymerase catalytic domain;Poly(ADP- | ENSG00000129484 | 14 | + | 19881613 | 11 | 11 | 20 | 12 | 0.496281504 | 2.014985429 | 0.122464029 |
| BCL9 | B-cell CLL/lymphoma 9 protein;B-cell lymphoma 9 protein;Bcl-9;Protein legless homolog | 149.3 | O00512;Q1JQ81;Q5T489 | | | ENSG00000116128 | 1 | + | 145479806 | 0 | 0 | 0 | 1 | 0.500559895 | 1.997762926 | 0.125378848 |
| RSBN1 | Round spermatid basic protein 1 | 45.7 | Q5VWQ0 | | | ENSG00000081019 | 1 | - | 114105977 | 0 | 1 | 1 | 1 | 0.505996054 | 1.976299997 | 0.129115055 |
| MPG;AAG;ANPG;MID1;PIG16;Z69720.1-002;Z69720.1-004 | DNA-3-methyladenine glycosylase;3-methyladenine DNA glycosidase;ADPG;3-alkyladenine DN | 32.2 | P29372;Q5J9I4;Q1W6H1;A2IDA3 | PF02245 | Methylpurine-DNA glycosylase (MPG) | ENSG00000103152 | 16 | + | 67018 | 17 | 23 | 21 | 35 | 0.519188399 | 1.926083099 | 0.138326518 |
| PRR6 | Proline-rich protein 6;Nuclear protein p30 | 29.9 | Q7Z7K6 | PF04828 | Glutathione-dependent formaldehyde-activating enzyme | ENSG00000166582 | 17 | - | 16186574 | 0 | 1 | 1 | 1 | 0.526996164 | 1.897547018 | 0.143869119 |
| NTHL1;NTH1;OCTS3 | Endonuclease III-like protein 1 | 34.4 | P78549 | PF00633;PF00730 | Helix-hairpin-helix motif;HhH-GPD superfamily base exci | ENSG00000065057 | 16 | - | 2029817 | 2 | 6 | 3 | 3 | 0.532303234 | 1.878628452 | 0.147672777 |
| DIMT1L;DIMT1;HUSSY-05 | Probable dimethyladenosine transferase;S-adenosylmethionine-6-N',N'-adenosyl(rRNA) dimet | 35.2 | Q9UNQ2;A8K9K8 | PF00398 | Ribosomal RNA adenine dimethylase | ENSG00000086189 | 5 | - | 61720108 | 3 | 8 | 4 | 6 | 0.535220129 | 1.868390117 | 0.149775376 |
| RBM14;SIP | RNA-binding protein 14;RNA-binding motif protein 14;RRM-containing coactivator activator | 69.5 | Q96PK6 | PF00076 | RNA recognition motif. (a.k.a. RRM, RBD, or RNP domain) | ENSG00000173959 | 11 | + | 66140673 | 1 | 4 | 5 | 14 | 0.541923909 | 1.845277506 | 0.154638794 |
| HIST1H4A;H4/A;H4FA;HIST1H4B;H4/I;H4FI;HIST1H4C;H4/G;H4FG;HIST1H4D;H4/B;H4FB;HIS | Histone H4 | 11.4 | P62805;A2VCL0;Q0VAS5;Q6B823;Q6 | PF00125 | Core histone H2A/H2B/H3/H4 | ENSG00000124529;ENSG0 | 6 | - | 26135022 | 10 | 8 | 11 | 11 | 0.54336874 | 1.840370868 | 0.155692498 |
| TOP3A;TOP3 | DNA topoisomerase 3-alpha;DNA topoisomerase III alpha | 112.4 | Q13472;A6NCL8;A8K398;A8KA61 | PF01131;PF01751;PF01396;PF06839 | DNA topoisomerase;Toprim domain;Topoisomerase DNA | ENSG00000177302 | 17 | - | 18117960 | 17 | 16 | 11 | 17 | 0.546568431 | 1.829597069 | 0.158032765 |
| H2AFZ;H2AZ;H2AFV;H2AV;hCG_19242 | Histone H2A.Z;H2A/z;Histone H2AV;H2A.F/Z;Uncharacterized protein H2AFV;H2A histone fam | 13.6 | P0C0S5;Q71UI9;A6NN01;A6NKY0;A6N | PF00125 | Core histone H2A/H2B/H3/H4 | ENSG00000164032;ENSG0 | 4 | - | 101088271 | 3 | 2 | 6 | 2 | 0.547417414 | 1.826759571 | 0.158655254 |
| DDX52;ROK1;HUSSY-19 | Probable ATP-dependent RNA helicase DDX52;DEAD box protein 52;ATP-dependent RNA helic | 67.5 | Q9Y2R4;Q59H21;A8MTP9 | PF00270;PF00271 | DEAD/DEAH box helicase;Helicase conserved C-terminal d | ENSG00000141141 | 17 | - | 33046526 | 0 | 2 | 1 | 5 | 0.549697297 | 1.819183039 | 0.160330046 |
| SSB | Lupus La protein;Sjoegren syndrome type B antigen;SS-B;La ribonucleoprotein;La autoantige | 46.8 | P05455;Q14730;Q15367;Q2F832;Q5 | PF05383;PF00076;PF08777 | La domain;RNA recognition motif. (a.k.a. RRM, RBD, or R | ENSG00000138385 | 2 | + | 170363605 | 15 | 8 | 12 | 20 | 0.550009397 | 1.818150754 | 0.160559666 |
| NAT10;ALP;KIAA1709 | N-acetyltransferase 10 | 115.7 | Q9H0A0;Q05CW7 | PF00583;PF08351;PF05127 | Acetyltransferase (GNAT) family;Domain of unknown func | ENSG00000135372 | 11 | + | 34083725 | 43 | 25 | 37 | 40 | 0.55580698 | 1.799185753 | 0.164840188 |
| RPL8 | 60S ribosomal protein L8 | 28.0 | P62917;A8K094 | PF00181;PF03947 | Ribosomal Proteins L2, RNA binding domain;Ribosomal P | ENSG00000161016 | 8 | - | 145985958 | 0 | 0 | 0 | 4 | 0 | #DIV/0! | 0.167923115 |
| SERBP1;PAIRBP1;CGI-55 | Plasminogen activator inhibitor 1 RNA-binding protein;PAI1 RNA-binding protein 1;PAI-RBP1 | 45.0 | Q8NC51;Q5VU19;Q5VU22;Q63HR1;Q | PF04774 | Hyaluronan / mRNA binding family | ENSG00000142864 | 1 | - | 67646081 | 0 | 1 | 1 | 0 | 0.561758619 | 1.780124001 | 0.169263221 |
| MYH9;FLJ00279 | Myosin-9;Myosin heavy chain 9;Myosin heavy chain, non-muscle IIa;Non-muscle myosin heav | 226.53 | P35579;A8K6E4;Q60FE2;Q86XU5;Q9 | PF00612;PF00063;PF02736;PF01576 | IQ calmodulin-binding motif;Myosin head (motor domai | ENSG00000100345 | 22 | - | 35007273 | 0 | 0 | 2 | 1 | 0 | #DIV/0! | 0.173240676 |
| KRT8;CYK8;KRT7;SCL | Keratin, type II cytoskeletal 8;Cytokeratin-8;CK-8;Keratin-8;K8;Keratin, type II cytoskeletal 7; | 53.7 | P05787;A8K4H3;Q7L4M3;Q969I0;A8 | PF00038 | Intermediate filament protein | ENSG00000170421;ENSG0 | 12 | - | 51577238 | 2 | 1 | 4 | 1 | 0.570191964 | 1.753795325 | 0.175577581 |
| RPS12;RP5-1181K21.3-001;hCG_2030596 | 40S ribosomal protein S12 | 14.5 | P25398;Q76M58;A6NCP0;A6NJZ8 | PF01248 | Ribosomal protein L7Ae/L30e/S12e/Gadd45 family | ENSG00000112306 | 6 | + | 133177374 | 3 | 1 | 4 | 4 | 0.571951517 | 1.748399945 | 0.176901635 |
| | | 18.3 | | | | | | | | 0 | 4 | 3 | 2 | 0.574932346 | 1.739335085 | 0.179149687 |
| NOL5;NOP5;HSPC120 | Nucleolar protein 5;Nucleolar protein NOP5;NOP58 | 59.6 | Q9Y2X3 | PF01798;PF08156;PF08060 | Putative snoRNA binding domain;NOP5NT (NUC127) dom | ENSG00000055044 | 2 | + | 202838702 | 0 | 0 | 2 | 2 | 0 | #DIV/0! | 0.189294858 |
| APTX;AXA1 | Aprataxin;Forkhead-associated domain histidine triad-like protein;FHA-HIT | 29.1 | Q7Z2E3;Q0P662 | PF01230 | HIT domain | ENSG00000137074 | 9 | - | 32962609 | 1 | 2 | 2 | 2 | 0.589975177 | 1.694986566 | 0.190584245 |
| ZC3HAV1;ZC3HDC2;PRO1677 | Zinc finger CCCH type antiviral protein 1;Zinc finger CCCH domain-containing protein 2;Zinc f | 101.4 | Q7Z2W4;A4D1S4;A8K9U6;A4D1R2 | PF00644;PF00642 | Poly(ADP-ribose) polymerase catalytic domain;Zinc finge | ENSG00000105939 | 7 | - | 138378808 | 12 | 15 | 21 | 30 | 0.602872299 | 1.658726072 | 0.200494583 |
| PNKP | Bifunctional polynucleotide phosphatase/kinase;Polynucleotide kinase-3'-phosphatase;DNA | 57.1 | Q96T60;Q9BUL2 | PF08645 | Polynucleotide kinase 3 phosphatase | ENSG00000039650 | 19 | - | 55056274 | 32 | 53 | 61 | 50 | 0.615897193 | 1.623647602 | 0.210588225 |
| MSH2 | DNA mismatch repair protein Msh2;MutS protein homolog 2 | 104.7 | P43246;A5JTV2;Q0ZAI9;Q53FK0;Q53 | PF01624;PF05188;PF05192;PF05190;PF | MutS domain I;MutS domain II;MutS domain III;MutS fam | ENSG00000095002 | 2 | + | 47483767 | 11 | 13 | 10 | 13 | 0.615942548 | 1.623528043 | 0.210623507 |
| CT45-5;CT45-2;CT45-4;CT45-6;CT45-1;CT45-3 | Cancer/testis antigen 45-5;CT45-5;Cancer/testis antigen 45-2;CT45-2;Cancer/testis antigen 4 | 21.1 | Q6NSH3;A8K842;Q5DJT8;Q8N7B7;Q5HYN5;Q8NHU0 | | | ENSG00000213441;ENSG0 | X | - | 134773630 | 12 | 14 | 11 | 14 | 0.62220211 | 1.607194808 | 0.215500848 |
| MSH3;DUC1;DUG | MutS homolog 3;E. coli;DNA mismatch repair protein Msh3;Divergent upstream protein;DUP | 127.4 | A1L480;A1L482;A6NMM6;A8K1E1;P2 | PF01624;PF05188;PF05192;PF00488 | MutS domain I;MutS domain II;MutS domain III;MutS do | ENSG00000113318 | 5 | + | 79986050 | 7 | 2 | 14 | 7 | 0.628995814 | 1.5898357 | 0.220811412 |
| POLR3A | DNA-directed RNA polymerase III subunit RPC1;RNA polymerase III subunit C1;DNA-directed | 155.8 | O14802;Q7Z755;Q8IW34;Q8TCW2;Q | PF04997;PF00623;PF04983;PF05000;PF | RNA polymerase Rpb1, domain 1;RNA polymerase Rpb1, | ENSG00000148606 | 10 | - | 79405912 | 2 | 2 | 4 | 3 | 0.630610336 | 1.585765317 | 0.222075875 |
| ZFP64;ZNF338;RP5-831D17.3-008;RP5-831D17.3-009 | Zinc finger protein 64, isoforms 1 and 2;Zinc finger protein 338;Zinc finger protein 64 homolo | 74.6 | Q9NPA5;Q5JWM1;Q9NTW7;Q53XQ1; | PF00096 | Zinc finger, C2H2 type | ENSG00000020256 | 20 | - | 50101609 | 2 | 1 | 4 | 7 | 0.633307505 | 1.579011764 | 0.224190184 |
| RMI1;C9orf76 | Protein RMI1 homolog;75 kDa BLM-associated protein;BLAP75;FAAP75 | 70.1 | Q9H9A7;Q05BX1;Q05CW3 | PF08585;PF01336 | Domain of unknown function (DUF1767);OB-fold nucleic | ENSG00000178966 | 9 | + | 85785533 | 12 | 6 | 9 | 20 | 0.644407415 | 1.551813305 | 0.232914659 |
| H2AFX;H2AX | Histone H2A.x;H2a/x | 15.1 | P16104 | PF00125 | Core histone H2A/H2B/H3/H4 | ENSG00000188486 | 11 | - | 118469799 | 4 | 3 | 2 | 1 | 0.650117222 | 1.53818414 | 0.237415361 |
| CEBPB;TCF5;PP9092 | CCAAT/enhancer-binding protein beta;C/EBP beta;Nuclear factor NF-IL6;Transcription factor | 36.1 | P17676;A8K671;Q99557;Q9BSC0 | PF07716 | Basic region leucine zipper | ENSG00000172216 | 20 | + | 48240798 | 1 | 4 | 4 | 1 | 0.660444953 | 1.514130732 | 0.245574114 |
| SRP68 | Signal recognition particle 68 kDa protein;SRP68;CDNA FLJ14414 fis, clone HEMBA1004847, | 70.7 | Q9UHB9;Q96K97;Q96K98 | | | ENSG00000167881 | 17 | - | 71546786 | 1 | 3 | 8 | 7 | 0.662349162 | 1.509777709 | 0.247080573 |
| DHX15;DBP1;DDX15 | Putative pre-mRNA-splicing factor ATP-dependent RNA helicase DHX15;DEAH box protein 15; | 90.9 | O43143 | PF07717;PF04408;PF00271 | Domain of unknown function (DUF1605);Helicase associa | ENSG00000109606 | 4 | - | 24138188 | 22 | 26 | 29 | 30 | 0.665424567 | 1.502799942 | 0.249514805 |
| LACRT | Extracellular glycoprotein lacritin precursor | 14.2 | Q9GZZ8 | | | ENSG00000135413 | 12 | - | 53310864 | 3 | 0 | 1 | 1 | 0.672777601 | 1.486375287 | 0.255340135 |
| RPA3;REPA3;RPA14 | Replication protein A 14 kDa subunit;RP-A;RF-A;Replication factor-A protein 3;p14 | 13.6 | P35244;A4D105 | PF08661 | Replication factor A protein 3 | ENSG00000106399 | 7 | - | 7643100 | 6 | 10 | 4 | 7 | 0.674061186 | 1.483544848 | 0.25635769 |
| CSDA;DBPA | DNA-binding protein A;Cold shock domain-containing protein A;Single-strand DNA-binding p | 31.947 | P16989;Q96B76;Q96GD7 | PF00313 | 'Cold-shock' DNA-binding domain | ENSG00000060138 | 12 | - | 10742956 | 0 | 0 | 1 | 1 | 0 | #DIV/0! | 0.260961346 |
| RPS5 | 40S ribosomal protein S5 | 22.9 | P46782;Q53G25;A6NIM7 | PF00177 | Ribosomal protein S7p/S5e | ENSG00000083845 | 19 | + | 63590448 | 0 | 1 | 1 | 0 | 0.688238113 | 1.452985501 | 0.267605222 |
| SUB1;PC4;RPO2TC1 | Activated RNA polymerase II transcriptional coactivator p15;SUB1 homolog;Positive cofactor | 14.4 | P53999;Q59G24;Q6IBA2 | PF02229 | Transcriptional Coactivator p15 (PC4) | ENSG00000113387 | 5 | + | 32621445 | 10 | 8 | 12 | 16 | 0.691950617 | 1.445189838 | 0.27055227 |
| LIG3 | DNA ligase 3;DNA ligase III;Polydeoxyribonucleotide synthase [ATP] 3 | 112.9 | P49916;Q6NVK3;Q96DF0 | PF04679;PF01068;PF04675;PF00645 | ATP dependent DNA ligase C terminal region;ATP depend | ENSG00000005156 | 17 | + | 30331651 | 96 | 79 | 98 | 128 | 0.702929955 | 1.422616853 | 0.279268289 |
| KPRP;C1orf45 | Keratinocyte proline-rich protein;hKPRP | 64.1 | Q5T749 | | | ENSG00000203786 | 1 | + | 150997130 | 0 | 6 | 1 | 3 | 0.704016637 | 1.420420977 | 0.280130839 |
| PURA;PUR1 | Transcriptional activator protein Pur-alpha;Purine-rich single-stranded DNA-binding protein | 34.9 | Q00577;Q2NLC9;Q2NLD4;Q56A79;Q5 | PF04845 | PurA ssDNA and RNA-binding protein | ENSG00000185129 | 5 | + | 139473892 | 13 | 13 | 17 | 8 | 0.706730883 | 1.414965759 | 0.282285056 |
| HIST2H2AA3;H2AFO;HIST2H2AA;HIST2H2AA4;HIST2H2AC;H2AFQ | Histone H2A type 2-A;H2A.2;H2A/o;Histone H2A type 2-C;H2A-GL101;H2A/q | 14.1 | Q6FI13;Q16777 | PF00125 | Core histone H2A/H2B/H3/H4 | ENSG00000183558;ENSG0 | 1 | - | 148080129 | 19 | 22 | 18 | 19 | 0.708819614 | 1.410796174 | 0.283942578 |
| | | 16.6 | | | | | | | | 13 | 2 | 5 | 14 | 0.713928921 | 1.400699663 | 0.287995941 |
| TFAM;TCF6L2 | Transcription factor A, mitochondrial precursor;mtTFA;Mitochondrial transcription factor 1; | 29.1 | Q00059;Q6LES8;A8MRB2;Q05D31 | PF00505 | HMG (high mobility group) box | ENSG00000108064 | 10 | + | 59815182 | 12 | 9 | 11 | 18 | 0.713956327 | 1.400645896 | 0.288017678 |
| HIST2H2BE;H2BFQ;HIST1H2BO;H2BFH;H2BFN;HIST1H2BB;H2BFF;HIST1H2BJ;H2BFR;HIST3H2 | Histone H2B type 2-E;H2B.q;H2B/q;H2B-GL105;Histone H2B type 1-O;H2B.n;H2B/n;H2B.2;Hi | 13.9 | Q16778;A8K110;P23527;P33778;P06 | PF00125 | Core histone H2A/H2B/H3/H4 | ENSG00000184678;ENSG0 | 1 | - | 148080395 | 18 | 24 | 27 | 27 | 0.718372952 | 1.392034593 | 0.291519846 |
| CREB1 | cAMP response element-binding protein;CREB;Putative uncharacterized protein CREB1 | 36.7 | P16220;Q4ZG78;Q5U0J5;Q53X93;Q5 | PF00170;PF02173 | bZIP transcription factor;pKID domain | ENSG00000118260 | 2 | + | 208102931 | 3 | 1 | 1 | 3 | 0.735589957 | 1.359453035 | 0.305150637 |
| TTF1 | Transcription termination factor 1;TTF-1;TTF-I;RNA polymerase I termination factor | 103.1 | Q15361 | PF00249 | Myb-like DNA-binding domain | ENSG00000125482 | 9 | - | 134240756 | 0 | 0 | 0 | 2 | 0 | #DIV/0! | 0.305495129 |
| RAD23B | UV excision repair protein RAD23 homolog B;hHR23B;XP-C repair-complementing complex 5 | 43.2 | P54727;Q53F10;Q7Z5K8 | PF00627;PF00240;PF09280 | UBA/TS-N domain;Ubiquitin family;XPC-binding domain | ENSG00000119318 | 9 | + | 109085365 | 13 | 16 | 14 | 14 | 0.750250531 | 1.332888093 | 0.316719615 |
| PIP5K1A;RP11-429H9.6-001 | Phosphatidylinositol-4-phosphate 5-kinase type-1 alpha;Phosphatidylinositol-4-phosphate 5 | 62.6 | Q99755;A8K4Q0;P78517;A2A3N6;Q6 | PF01504;PF02809 | Phosphatidylinositol-4-phosphate 5-Kinase;Ubiquitin in | ENSG00000143398;ENSG0 | 1 | + | 149437696 | 4 | 6 | 8 | 6 | 0.752422725 | 1.329040135 | 0.31843001 |
| POLR3D;BN51;BN51T | DNA-directed RNA polymerase III subunit RPC4;RNA polymerase III subunit C4;DNA-directed | 44.4 | P05423 | PF05132 | RNA polymerase III RPC4 | ENSG00000168495 | 8 | + | 22158564 | 1 | 0 | 0 | 0 | 0.75351984 | 1.327105071 | 0.319293478 |
| VIM | Vimentin | 53.7 | P08670;Q53HU8;Q548L2;Q8N9S6 | PF00038;PF04732 | Intermediate filament protein;Intermediate filament hea | ENSG00000026025 | 10 | + | 17311250 | 1 | 0 | 0 | 0 | 0.75595916 | 1.322822783 | 0.321212311 |
| NOSTRIN | Nostrin;eNOS trafficking inducer;Nitric oxide synthase trafficker;Nitric oxide synthase traffic i | 57.7 | Q8IVI9;A8K2I9;Q53TH8 | PF00611;PF02185;PF00018 | Fes/CIP4 homology domain;Hr1 repeat;SH3 domain | ENSG00000163072 | 2 | + | 169367353 | 13 | 17 | 28 | 25 | 0.765503919 | 1.30632904 | 0.328706365 |
| POLR3B | DNA-directed RNA polymerase III subunit RPC2;RNA polymerase III subunit C2;DNA-directed | 127.8 | Q9NW08;A8K6H0;Q7Z3R8 | PF04563;PF04561;PF04565;PF04566;PF | RNA polymerase beta subunit;RNA polymerase Rpb2, do | ENSG00000013503 | 12 | + | 105275619 | 1 | 1 | 2 | 2 | 0.766324799 | 1.304929712 | 0.329349768 |
| DSP | Desmoplakin;DP;250/210 kDa paraneoplastic pemphigus antigen | 331.8 | P15924;Q4LE79 | PF00681 | Plectin repeat | ENSG00000096696 | 6 | + | 7486869 | 71 | 77 | 97 | 17 | 0.773614532 | 1.292633422 | 0.335055165 |
| GNB2L1 | Guanine nucleotide binding protein (G protein), beta polypeptide 2-like 1 variant;Guanine nu | 37.9 | Q53HU2;Q5VLR4;P63244;Q6FH47;Q5 | PF00400 | WD domain, G-beta repeat | ENSG00000204628 | 5 | - | 180596531 | 5 | 6 | 7 | 5 | 0.773930342 | 1.292105951 | 0.33530199 |
| CD3EAP;ASE1;CAST;PAF49 | DNA-directed RNA polymerase I subunit RPA34;RNA polymerase I-associated factor PAF49;An | 55.2 | O15446;A8K818;A0JLT1;Q05BI1 | | | ENSG00000117877 | 19 | + | 50601307 | 9 | 10 | 6 | 8 | 0.782750403 | 1.277546451 | 0.342183117 |
| CETN2;CALT;CEN2 | Centrin-2;Caltractin isoform 1 | 19.7 | P41208 | PF00036 | EF hand | ENSG00000147400 | X | - | 151746529 | 0 | 1 | 1 | 2 | 0.786077776 | 1.272138751 | 0.344772594 |
| SRP14 | Signal recognition particle 14 kDa protein;SRP14;18 kDa Alu RNA-binding protein | 14.6 | P37108;Q96Q14;A8MVS4 | PF02290 | Signal recognition particle 14kD protein | ENSG00000140319 | 15 | - | 38115535 | 50 | 45 | 73 | 45 | 0.787603546 | 1.269674324 | 0.345958774 |
| ZNF512B;KIAA1196 | Zinc finger protein 512B | 97.3 | Q96KM6 | PF00096 | Zinc finger, C2H2 type | ENSG00000196700 | 20 | - | 62058499 | 1 | 1 | 0 | 5 | 0 | #DIV/0! | 0.349001963 |
| RPS28 | Uncharacterized protein ENSP00000331275;40S ribosomal protein S28 | 7.8 | A6NNT3;P62857 | PF01200 | Ribosomal protein S28e | ENSG00000182953;ENSG0 | 17 | + | 18416760 | 2 | 3 | 1 | 0 | 0.794506577 | 1.258642821 | 0.351315422 |
| | | 68.1 | P15636 | | | | | | | 152 | 200 | 190 | 170 | 0.807936951 | 1.237720343 | 0.361687604 |
| EIF6;EIF3A;ITGB4BP;OK/SW-cl.27 | Eukaryotic translation initiation factor 6;eIF-6;B4 integrin interactor;CAB;p27(BBP);B(2)GCN | 26.6 | P56537;Q6IBN8;Q96TD5 | PF01912 | eIF-6 family | ENSG00000126005 | 20 | - | 33330139 | 5 | 7 | 3 | 1 | 0.810743988 | 1.233434987 | 0.363846712 |
| RPS10;RP11-375E1__A.4-005;LOC376693;RPS10L;RP11-371L19.1-001 | Ribosomal protein S10;40S ribosomal protein S10;Ribosomal protein S10 variant;LOC37669 | 19.9 | Q5TZB9;P46783;Q59GE4;A6NJ00;Q6P | PF03501 | Plectin/S10 domain | ENSG00000124614;ENSG0 | 6 | - | 34493209 | 2 | 3 | 2 | 2 | 0.812422348 | 1.230886868 | 0.365136166 |
| POLR3H;KIAA1665 | DNA-directed RNA polymerase III subunit RPC8;RNA polymerase III subunit C8;DNA-directed | 22.9 | Q9Y535;A6NEV8;Q6ZPA8 | PF08292;PF03876 | RNA polymerase III subunit Rpc25;RNA polymerase Rpb7 | ENSG00000100413 | 22 | - | 40251754 | 1 | 1 | 1 | 0 | 0.814466604 | 1.227797427 | 0.366705189 |
| RSBN1 | Round spermatid basic protein 1 | 90.1 | Q5VWQ0;A8K937 | | | ENSG00000081019 | 1 | - | 114105977 | 10 | 14 | 7 | 8 | 0.83529394 | 1.197183353 | 0.382589673 |
| ARG1 | Arginase-1;Type I arginase;Liver-type arginase | 35.7 | P05089;A6NEA0 | PF00491 | Arginase family | ENSG00000118520 | 6 | + | 131936058 | 11 | 5 | 5 | 1 | 0.837841469 | 1.193543214 | 0.384519389 |
| POLR1C;POLR1E;RP3-337H4.4;RP3-337H4.4-004;RP3-337H4.4-007 | DNA-directed RNA polymerases I and III subunit RPAC1;RNA polymerases I and III subunit AC1 | 39.2 | O15160;Q96HT3;Q5JTE4;Q5JTE0;Q5J | PF01000;PF01193 | RNA polymerase Rpb3/RpoA insert domain;RNA polymer | ENSG00000171453 | 6 | + | 43592769 | 3 | 1 | 0 | 1 | 0.841111907 | 1.188902441 | 0.386992287 |
| POLB | DNA polymerase beta | 38.2 | P06746;Q53EV2;Q6LBJ9;Q9BQF2 | | | ENSG00000070501 | 8 | + | 42315131 | 41 | 56 | 33 | 43 | 0.842796008 | 1.186526741 | 0.38826374 |
| SHROOM3;KIAA1481;SHRML;MSTP013 | Protein Shroom3;Shroom;Shroom-related protein;hShrmL;CDNA FLJ42424 fis, clone BLADE2 | 216.9 | Q8TF72;Q6ZVL3 | PF08688;PF08687;PF00595 | Apx/Shroom domain ASD1;Apx/Shroom domain ASD2;P | ENSG00000138771 | 4 | + | 77575277 | 1 | 1 | 1 | 1 | 0.850256048 | 1.176116304 | 0.393879598 |
| RPL38 | 60S ribosomal protein L38 | 8.2 | P63173 | PF01781 | Ribosomal L38e protein family | ENSG00000172809 | 17 | + | 69711390 | 1 | 4 | 9 | 3 | 0.851567507 | 1.174305021 | 0.394864068 |
| RFC5 | Replication factor C subunit 5;Replication factor C 36 kDa subunit;RF-C 36 kDa subunit;RFC3 | 38.5 | P40937;A8K3S0;Q59GW7;A8K4Z2;A8 | PF00004;PF08542 | ATPase family associated with various cellular activities ( | ENSG00000111445 | 12 | + | 116938893 | 24 | 31 | 27 | 25 | 0.872617286 | 1.145977757 | 0.410546924 |
| SRP9;hCG_1984300;DKFZp564M2223 | Signal recognition particle 9 kDa protein;SRP9;SRP9 protein;Signal recognition particle 9kDa | 10.1 | P49458;A8K0N0;Q8WVW9;Q6NVX0; | PF05486 | Signal recognition particle 9 kDa protein (SRP9) | ENSG00000143742;ENSG0 | 1 | + | 224032151 | 46 | 39 | 43 | 37 | 0.881765818 | 1.134087962 | 0.41729075 |
| HMGB1;HMG1;RP11-550P23.1-002;RP11-550P23.1-003;HMG1L10;RP11-550P23.1-005 | High mobility group protein B1;High mobility group protein 1;HMG-1;High-mobility group b | 24.9 | P09429;A5D8W9;Q14321;Q5T7C3;Q | PF00505 | HMG (high mobility group) box | ENSG00000189403;ENSG0 | 13 | - | 29930877 | 1 | 2 | 4 | 2 | 0.885864676 | 1.128840587 | 0.42029751 |
| FBL;FIB1;FLRN | rRNA 2'-O-methyltransferase fibrillarin;34 kDa nucleolar scleroderma antigen | 33.8 | P22087;Q96BS4 | PF01269 | Fibrillarin | ENSG00000105202 | 19 | - | 45016938 | 13 | 9 | 15 | 17 | 0.891037606 | 1.122287088 | 0.424078945 |
| LIN28B;CSDD2 | Lin-28 homolog B | 29.9 | Q6ZN17;A7E2T3 | PF00313;PF00098 | 'Cold-shock' DNA-binding domain;Zinc knuckle | ENSG00000187772 | 6 | + | 105511616 | 9 | 15 | 16 | 11 | 0.895337305 | 1.116897502 | 0.427210678 |
| | cDNA FLJ76546, highly similar to Homo sapiens cAMP responsive element modulator (CREM) | 12.1 | A8K3J7 | | | | | | | 4 | 3 | 1 | 1 | 0.898700027 | 1.112718337 | 0.429652682 |
| RSBN1L | Round spermatid basic protein 1-like protein | 94.9 | Q6PCB5 | | | ENSG00000187257 | 7 | + | 77163696 | 34 | 29 | 31 | 38 | 0.911549611 | 1.097032996 | 0.438924257 |
| | | 24.4 | P00761 | | | | | | | 71 | 110 | 74 | 78 | 0.913156858 | 1.095102108 | 0.440077202 |
| DYNLL1;DLC1;DNCL1;DNCLC1;HDLC1 | Dynein light chain 1, cytoplasmic;Dynein light chain LC8-type 1;8 kDa dynein light chain;DLC | 10.4 | P63167;Q6FGH9 | PF01221 | Dynein light chain type 1 | ENSG00000088986 | 12 | + | 119392043 | 6 | 6 | 0 | 5 | 0.914144864 | 1.093918524 | 0.440785186 |
| POLR2H | DNA-directed RNA polymerases I, II, and III subunit RPABC3;RNA polymerases I, II, and III subu | 17.1 | P52434 | PF03870 | RNA polymerase Rpb8 | ENSG00000163882 | 3 | + | 185563888 | 2 | 2 | 1 | 1 | 0.91831229 | 1.088954173 | 0.443765122 |
| RPSA;LAMBR;LAMR1;LAMR1P15 | Ribosomal protein SA;40S ribosomal protein SA;p40;34/67 kDa laminin receptor;Colon carci | 33.3 | Q6NSD1;Q6NXQ8;Q86VC0;P08865;A6 | PF00318 | Ribosomal protein S2 | ENSG00000168028;ENSG0 | 3 | + | 39423208 | 5 | 7 | 12 | 7 | 0.927573376 | 1.078081828 | 0.45035019 |
| XRCC5;G22P2 | ATP-dependent DNA helicase 2 subunit 2;ATP-dependent DNA helicase II 80 kDa subunit;Lupu | 82.7 | P13010;A8K3X5;Q0Z7V0;Q53HH7;Q5 | PF02735;PF03730;PF03731;PF08785 | Ku70/Ku80 beta-barrel domain;Ku70/Ku80 C-terminal a | ENSG00000079246 | 2 | + | 216675717 | 184 | 219 | 178 | 191 | 0.928739032 | 1.076728732 | 0.451175365 |
| BANF1;BAF;BCRG1 | Barrier-to-autointegration factor;Breakpoint cluster region protein 1 | 10.1 | O75531;A6NCY2 | PF02961 | Barrier to autointegration factor | ENSG00000175334 | 11 | + | 65526126 | 10 | 19 | 13 | 12 | 0.9342928 | 1.070328274 | 0.455095563 |
| HMGN2;HMG17;HMGN3;TRIP7;PNAS-24;TRMT5 | Non-histone chromosomal protein HMG-17;High-mobility group nucleosome-binding domai | 32.7 | A6NDH4;A6NG37;A6NH59;P05204;Q | PF01101 | HMG14 and HMG17 | ENSG00000198830;ENSG0 | 1 | + | 26671593 | 1 | 3 | 2 | 2 | 0.93831973 | 1.065734811 | 0.457926217 |
| RPA1;REPA1;RPA70 | Replication protein A 70 kDa DNA-binding subunit;RP-A;RF-A;Replication factor-A protein 1;S | 68.1 | P27694;A8K0Y9 | PF04057;PF08646;PF01336 | Replication factor-A protein 1, N-terminal domain;Replic | ENSG00000132383 | 17 | + | 1680095 | 24 | 22 | 24 | 21 | 0.940009923 | 1.063818557 | 0.45911133 |
| | | 63.2 | | | | | | | | 20 | 13 | 12 | 2 | 0.9416833 | 1.061928145 | 0.460282912 |
| APTX;AXA1;RP11-562M8.1-005 | Aprataxin;Forkhead-associated domain histidine triad-like protein;FHA-HIT;Aprataxin variant | 40.7 | Q7Z2E3;Q5T784;Q6JV85;Q5T782;Q5T | PF01230 | HIT domain | ENSG00000137074 | 9 | - | 32962609 | 32 | 25 | 30 | 40 | 0.949401536 | 1.053295115 | 0.465664144 |
| NOLA2;NHP2;HSPC286 | H/ACA ribonucleoprotein complex subunit 2;Nucleolar protein family A member 2;snoRNP p | 17.2 | Q9NX24;Q96P96 | PF01248 | Ribosomal protein L7Ae/L30e/S12e/Gadd45 family | ENSG00000145912 | 5 | - | 177509074 | 4 | 3 | 3 | 5 | 0.952275145 | 1.050116666 | 0.467658131 |
| HNRPD;AUF1 | Heterogeneous nuclear ribonucleoprotein D0;hnRNP D0;AU-rich element RNA-binding prote | 38.4 | Q14103;A8K9J2;Q12771;Q9UCE8 | PF00076 | RNA recognition motif. (a.k.a. RRM, RBD, or RNP domain) | ENSG00000138668 | 4 | - | 83493491 | 3 | 1 | 1 | 3 | 0.953903251 | 1.048324344 | 0.468785562 |
| POF1B | Protein POF1B;Premature ovarian failure protein 1B;Uncharacterized protein POF1B | 68.7 | Q8WVV4;A8K2U5;A8MVC4 | | | ENSG00000124429 | X | - | 84419058 | 5 | 2 | 1 | 0 | 0.961976907 | 1.039525993 | 0.474351619 |

| | | | | | | | | | | | | | | | | |
|---|---|---|---|---|---|---|---|---|---|---|---|---|---|---|---|---|
| C1orf77;HT031;PP7704;RP1-178F15.2-002 | Uncharacterized protein C1orf77;Chromosome 1 open reading frame 77 | 26.5 | Q9Y3Y2;Q0VAQ8;Q5T7Y7;Q5T7Y9 | | | ENSG00000160679 | 1 | + | 151873149 | 4 | 4 | 2 | 3 | 0.963306977 | 1.038090686 | 0.475264605 |
| PRPF19;NMP200;PRP19;SNEV | Pre-mRNA-processing factor 19;PRP19/PSO4 homolog;hPso4;Nuclear matrix protein 200;Ser | 55.2 | Q9UMS4 | PF08606;PF04564;PF00400 | Prp19/Pso4-like;U-box domain;WD domain, G-beta repe | ENSG00000110107 | 11 | - | 60414768 | 1 | 1 | 1 | 2 | 0.968817682 | 1.03218595 | 0.479035201 |
| CENPQ;C6orf139 | Centromere protein Q;CENP-Q | 30.6 | Q7L2Z9;A8KAF1 | | | ENSG00000031691 | 6 | + | 49539055 | 0 | 1 | 1 | 0 | 0.972825315 | 1.027933776 | 0.481765092 |
| WDR5;BIG3 | WD repeat-containing protein 5;BMP2-induced 3-kb gene protein | 36.6 | P61964 | PF00400 | WD domain, G-beta repeat | ENSG00000196363 | 9 | + | 135991031 | 8 | 5 | 6 | 7 | 0.973051349 | 1.027694994 | 0.481918751 |
| PKP1 | Plakophilin-1;Band-6 protein;B6P | 82.9 | Q13835;Q14BN3;Q14CA0 | PF00514 | Armadillo/beta-catenin-like repeat | ENSG00000081277 | 1 | + | 199519203 | 5 | 3 | 10 | 0 | 0.974570512 | 1.02609302 | 0.482950634 |
| TFAP2B | Transcription factor AP-2 beta;AP2-beta;Activating enhancer-binding protein 2 beta;cDNA FL | 51.5 | Q92481;A8K557 | PF03299 | Transcription factor AP-2 | ENSG00000008196 | 6 | + | 50894398 | 28 | 31 | 29 | 17 | 0.975717469 | 1.024886847 | 0.483728708 |
| TGM3 | Protein-glutamine gamma-glutamyltransferase E precursor;TGase E;TGE;TG(E);Transglutamina | 76.6 | Q08188;A8K5N6;Q32MM0 | PF00927;PF01841;PF00868 | Transglutaminase family, C-terminal ig like domain;Trans | ENSG00000125780 | 20 | + | 2224675 | 7 | 5 | 3 | 3 | 0.979071464 | 1.021375902 | 0.485999101 |
| NHP2L1 | NHP2-like protein 1;High mobility group-like nuclear protein 2 homolog 1;U4/U6.U5 tri-snRN | 14.2 | P55769;Q6FHM6 | PF01248 | Ribosomal protein L7Ae/L30e/S12e/Gadd45 family | ENSG00000100138 | 22 | - | 40399886 | 11 | 6 | 6 | 12 | 0.979538287 | 1.02088914 | 0.486314525 |
| RFC3 | Replication factor C subunit 3;Replication factor C 38 kDa subunit;RFC38;Activator 1 38 kDa | 40.6 | P40938;Q53HK8;Q5U0C2;Q5W0E8;A8MX32 | | | ENSG00000133119 | 13 | + | 33290206 | 64 | 57 | 55 | 50 | 0.987446056 | 1.012713549 | 0.491636072 |
| PWP1 | Periodic tryptophan protein 1 homolog;Keratinocyte protein IEF SSP 9502 | 55.8 | Q13610;A8K3R6;Q05BL3;Q53FW9;Q6 | PF00400 | WD domain, G-beta repeat | ENSG00000136045 | 12 | + | 106603720 | 11 | 11 | 9 | 12 | 0.988126415 | 1.012016261 | 0.49209201 |
| RFC4 | Replication factor C subunit 4;Replication factor C 37 kDa subunit;RF-C 37 kDa subunit;RFC37 | 39.7 | P35249 | PF00004;PF08542 | ATPase family associated with various cellular activities (A | ENSG00000163918 | 3 | - | 187990378 | 42 | 36 | 42 | 42 | 0.994693283 | 1.005335028 | 0.496477129 |
| HMGN4;HMG17L3;NHC | High mobility group nucleosome-binding domain-containing protein 4;Non-histone chromo | 9.5 | O00479 | PF01101 | HMG14 and HMG17 | ENSG00000182952 | 6 | + | 26646551 | 2 | 2 | 3 | 3 | 0.999735218 | 1.000264852 | 0.499824666 |
| CDC5L;KIAA0432;PCDC5RP | Cell division cycle 5-like protein;Cdc5-like protein;Pombe cdc5-related protein | 92.25 | Q99459 | PF00249 | Myb-like DNA-binding domain | ENSG00000096401 | 6 | + | 44463457 | 0 | 0 | 0 | 1 | 0 | #DIV/0! | 0.5 |
| NCL | Nucleolin;Protein C23;CDNA FLJ45706 fis, clone FEBRA2028457, highly similar to Nucleolin; | 76.613 | P19338;Q6ZS99;Q9BQ02;Q9UCF0 | PF00076 | RNA recognition motif. (a.k.a. RRM, RBD, or RNP domain) | ENSG00000115053 | 2 | - | 232027706 | 0 | 0 | 1 | 1 | 0 | #DIV/0! | 0.5 |
| KLF1;EKLF | Krueppel-like factor 1;Erythroid krueppel-like transcription factor;EKLF;Erythroid transcriptio | 38.22 | Q13351;Q6PIJ5 | PF00096 | Zinc finger, C2H2 type | ENSG00000105610 | 19 | - | 12856241 | 0 | 0 | 0 | 1 | 0 | #DIV/0! | 0.5 |
| RPL29 | cDNA FLJ78093, highly similar to Homo sapiens ribosomal protein L29 (RPL29), mRNA;60S ri | 27.349 | A6NJU4;A8K0H3;P47914;Q5T1D1;Q6 | PF01779 | Ribosomal L29e protein family | ENSG00000162244 | 3 | - | 52002685 | 0 | 0 | 0 | 1 | 0 | #DIV/0! | 0.5 |
| NGDN;C14orf120 | Neuroguidin;EIF4E-binding protein;Uncharacterized protein NGDN | 35.9 | Q8NEJ9;A8K760;A8MT00 | PF04000 | Sas10/Utp3/C1D family | ENSG00000129460 | 14 | + | 22994554 | 0 | 0 | 0 | 3 | 0 | #DIV/0! | 0.5 |
| DDB1;XAP1 | DNA damage-binding protein 1;Damage-specific DNA-binding protein 1;UV-damaged DNA-bir | 127.0 | Q16531;A6NG77 | PF03178 | CPSF A subunit region | ENSG00000167986 | 11 | - | 60823510 | 1 | 0 | 2 | 2 | 1.00465569 | 0.995365885 | 0.503075334 |
| JUP;hCG_1771506;CTNNG;DP3 | JUP protein;Junction plakoglobin, isoform CRA_a;Junction plakoglobin;Desmoplakin-3;Desm | 81.7 | Q7L3S5;P14923;Q15151;Q7KZ86;Q8 | PF00514;PF02985 | Armadillo/beta-catenin-like repeat;HEAT repeat | ENSG00000173801 | 17 | - | 37164382 | 18 | 19 | 28 | 2 | 1.014260028 | 0.985940461 | 0.509373973 |
| ASH2L;ASH2L1 | Set1/Ash2 histone methyltransferase complex subunit ASH2;ASH2-like protein | 68.7 | Q9UBL3;A8K6T5;A8K7C3 | PF00622 | SPRY domain | ENSG00000129691 | 8 | + | 38082223 | 3 | 5 | 6 | 5 | 1.014921263 | 0.985298108 | 0.509805353 |
| POLR2E | DNA-directed RNA polymerases I, II, and III subunit RPABC1;RNA polymerases I, II, and III subu | 24.6 | P19388 | PF01191;PF03871 | RNA polymerase Rpb5, C-terminal domain;RNA polymera | ENSG00000099817 | 19 | - | 1039155 | 2 | 5 | 4 | 7 | 1.015449698 | 0.984785364 | 0.510149887 |
| KRT9 | Keratin, type I cytoskeletal 9;Cytokeratin-9;CK-9;Keratin-9;K9 | 62.1 | P35527 | PF00038 | Intermediate filament protein | ENSG00000171403 | 17 | - | 36975622 | 129 | 163 | 164 | 87 | 1.031578145 | 0.969388509 | 0.520575336 |
| RPA2;REPA2;RPA32;RPA34;RP4-547C9.3-003 | Replication protein A 32 kDa subunit;RP-A;RF-A;Replication factor-A protein 2;p32;p34;Repli | 38.8 | P15927;Q5TEJ7 | PF08784;PF01336 | Replication protein A C terminal;OB-fold nucleic acid bin | ENSG00000117748 | 1 | - | 28090637 | 5 | 3 | 3 | 8 | 1.036187112 | 0.965076663 | 0.5235224 |
| HMGA1;HMGIY | High mobility group protein HMG-I/HMG-Y;HMG-I(Y);High mobility group AT-hook protein 1; | 36.8 | P17096;Q5T6U9;Q6IPL9;A6NM05;Q5T | PF02178 | AT hook motif | ENSG00000137309 | 6 | + | 34312628 | 0 | 1 | 1 | 0 | 1.043678624 | 0.958149354 | 0.528281976 |
| RPL5;MSTP030 | 60S ribosomal protein L5 | 34.4 | P46777;A2RUM7;Q32LZ3;Q59GX9;Q8 | PF00861 | Ribosomal L18p/L5e family | ENSG00000122406 | 1 | + | 93070182 | 1 | 1 | 0 | 1 | 1.046761773 | 0.955327206 | 0.530229749 |
| RPL7L1 | Ribosomal protein L7-like 1 | 28.7 | Q6DKI1;A8K5J5;A8K7D4 | PF00327 | Ribosomal protein L30p/L7e | ENSG00000146223 | 6 | + | 42955351 | 0 | 0 | 3 | 4 | 1.053829024 | 0.948920533 | 0.534670129 |
| HSPB1;HSP27 | Heat shock protein beta-1;HspB1;Heat shock 27 kDa protein;HSP 27;Stress-responsive protein | 22.8 | P04792;Q96C20;Q96EI7;Q9UC35 | PF00011 | Hsp20/alpha crystallin family | ENSG00000106211 | 7 | + | 75769859 | 4 | 2 | 4 | 1 | 1.059179242 | 0.944127264 | 0.538009115 |
| TAF15;RBP56;TAF2N;FUS;TLS | TATA-binding protein-associated factor 2N;RNA-binding protein 56;TAFII68;TAF(II)68;RNA-bir | 65.0 | Q92804;Q86X94;P35637;Q13344;Q6 | PF00076;PF00641 | RNA recognition motif. (a.k.a. RRM, RBD, or RNP domain) | ENSG00000172660;ENSG0 | 17 | + | 31160596 | 5 | 5 | 5 | 2 | 1.066763031 | 0.937415312 | 0.542708658 |
| XRCC6;G22P1 | ATP-dependent DNA helicase 2 subunit 1;ATP-dependent DNA helicase II 70 kDa subunit;Lupu | 69.8 | P12956;Q6FG89;Q6IC76 | PF02735;PF03730;PF03731;PF02037 | Ku70/Ku80 beta-barrel domain;Ku70/Ku80 C-terminal ar | ENSG00000196419 | 22 | + | 40347241 | 238 | 303 | 215 | 250 | 1.069440694 | 0.935068215 | 0.544358599 |
| | | 54.4 | | | | | | | | 5 | 2 | 0 | 3 | 1.083480941 | 0.922951168 | 0.552929982 |
| NPM1;NPM | Nucleophosmin;NPM;Nucleolar phosphoprotein B23;Numatrin;Nucleolar protein NO38;Nuc | 32.6 | P06748;A4ZU86;A8K3N7;Q9BTI9;Q9B | PF03066 | Nucleoplasmin | ENSG00000181163 | 5 | + | 170747403 | 1 | 1 | 0 | 0 | 1.102816463 | 0.906769198 | 0.564513867 |
| DCD;AIDD;DSEP | Dermcidin precursor;Preproteolysin;Survival-promoting peptide;DCD-1;Dermcidin isoform 2 | 11.3 | P81605;Q53YJ2;A5JHP3 | | | ENSG00000161634 | 12 | - | 53324642 | 11 | 10 | 9 | 13 | 1.106780332 | 0.903521658 | 0.566857108 |
| POLR1E;PAF53;PRAF1 | DNA-directed RNA polymerase I subunit RPA49;RNA polymerase I-associated factor 53;RNA po | 47.3 | Q9GZS1 | PF06870 | A49-like RNA polymerase I associated factor | ENSG00000137054 | 9 | + | 37475932 | 6 | 11 | 8 | 7 | 1.113291792 | 0.898237109 | 0.570683108 |
| PPP1CC;PPP1CA;PPP1A | Serine/threonine-protein phosphatase PP1-gamma catalytic subunit;PP-1G;Protein phospha | 38.5 | P36873;Q9UPN1;Q07161;P62136;A6 | PF00149 | Calcineurin-like phosphoesterase | ENSG00000186298;ENSG0 | 12 | - | 109642127 | 12 | 12 | 8 | 5 | 1.127507038 | 0.886912424 | 0.578935384 |
| PRKDC;HYRC;HYRC1 | DNA-dependent protein kinase catalytic subunit;DNA-PK catalytic subunit;DNA-PKcs;DNPK1; | 469.1 | P78527 | PF02259;PF02260;PF02985;PF08163;PF0 | FAT domain;FATC domain;HEAT repeat;NUC194 domain;I | ENSG00000121031 | 8 | - | 48848222 | 301 | 324 | 245 | 268 | 1.130226944 | 0.884778058 | 0.580498693 |
| CETN3;CEN3 | Centrin-3 | 19.6 | O15182;Q6ICP7 | PF00036 | EF hand | ENSG00000153140 | 5 | - | 89725287 | 1 | 2 | 3 | 2 | 1.132161718 | 0.883266042 | 0.581607678 |
| SSBP1;SSBP | Single-stranded DNA-binding protein, mitochondrial precursor;Mt-SSB;MtSSB;PWP1-interac | 17.3 | Q04837;A4D1U3;Q567R6;Q75MV9 | PF00436 | Single-strand binding protein family | ENSG00000106028 | 7 | + | 141084628 | 8 | 7 | 5 | 3 | 1.135814759 | 0.880425256 | 0.583694623 |
| CNBP;RNF163;ZNF9 | Cellular nucleic acid-binding protein;CNBP;Zinc finger protein 9 | 19.0 | P62633;Q4JGY0;Q4JGY1;Q6T598;A8K | PF00098 | Zinc knuckle | ENSG00000169714 | 3 | - | 130371105 | 2 | 3 | 8 | 3 | 1.137818112 | 0.878875094 | 0.584835277 |
| HNRPAB;ABBP1 | Heterogeneous nuclear ribonucleoprotein A/B;hnRNP A/B;APOBEC1-binding protein 1;ABBP- | 36.2 | Q99729;Q53F64 | PF08143;PF00076 | CBFNT (NUC161) domain;RNA recognition motif. (a.k.a. R | ENSG00000197451 | 5 | + | 177564114 | 1 | 2 | 1 | 1 | 1.138609461 | 0.878264264 | 0.5852851 |
| MLL;ALL1;CXXC7;HRX;HTRX;KMT2A;TRX1 | Zinc finger protein HRX;ALL-1;Trithorax-like protein;Lysine N-methyltransferase 2A;CXXC-type | 431.8 | Q03164;Q59FF2;Q9HBJ3;Q9HBJ4;Q9U | PF02178;PF05965;PF05964;PF00628;PF0 | AT hook motif;F/Y rich C-terminus;F/Y-rich N-terminus;PH | ENSG00000118058 | 11 | + | 117812415 | 0 | 0 | 0 | 3 | 1.142153851 | 0.875538789 | 0.587294618 |
| C19orf57 | Uncharacterized protein C19orf57;Pre-T/NK cell-associated protein 3B3 | 84.2 | Q0VDD7 | | | ENSG00000132016 | 19 | - | 13854168 | 3 | 5 | 1 | 1 | 1.14704646 | 0.871804268 | 0.590054554 |
| DPY30 | Protein dpy-30 homolog;Dpy-30-like protein | 11.3 | Q9C005 | PF05186 | Dpy-30 motif | ENSG00000162961 | 2 | - | 32102465 | 3 | 2 | 1 | 1 | 1.162924778 | 0.859900845 | 0.598900291 |
| KRT6A;K6A;KRT6D | Keratin, type II cytoskeletal 6A;Cytokeratin-6A;CK 6A;K6a keratin;Cytokeratin-6D;CK 6D | 60.0 | P02538;A4QPC1;A8K2I0 | PF00038 | Intermediate filament protein | ENSG00000205420 | 12 | - | 51167231 | 70 | 138 | 133 | 53 | 1.165213729 | 0.858211653 | 0.600161464 |
| RFC2 | Replication factor C subunit 2;Replication factor C 40 kDa subunit;RF-C 40 kDa subunit;RFC40 | 39.2 | P35250;Q75MT5 | PF00004;PF08542 | ATPase family associated with various cellular activities (A | ENSG00000049541 | 7 | - | 73283770 | 35 | 28 | 19 | 24 | 1.170613734 | 0.854252749 | 0.603122851 |
| EPPK1;EPIPL;PLEC1;hCG_1994702 | Epiplakin;450 kDa epidermal antigen;Plectin-1;PLTN;PCN;Hemidesmosomal protein 1;HD1;P | 555.6 | P58107;Q9NSU9;Q15149;Q6S380;Q6 | PF00681;PF00307;PF03501 | Plectin repeat;Calponin homology (CH) domain;Plectin/S | ENSG00000178209 | 8 | - | 145061309 | 0 | 1 | 1 | 1 | 1.173078321 | 0.852458001 | 0.60446795 |
| RAD23A | UV excision repair protein RAD23 homolog A;hHR23A | 39.6 | P54725;A8K1J3;Q59EU8;Q5M7Z1 | PF00627;PF00240;PF09280 | UBA/TS-N domain;Ubiquitin family;XPC-binding domain | ENSG00000179262 | 19 | + | 12917654 | 2 | 2 | 2 | 4 | 1.1771682 | 0.849496274 | 0.606691123 |
| APOBEC3C;APOBEC1L;PBI | Apolipoprotein B mRNA editing enzyme, catalytic polypeptide-like 3C variant;Probable DNA d | 25.2 | Q59GY0;Q9NRW3 | PF05240;PF08210 | APOBEC-like C-terminal domain;APOBEC-like N-terminal d | ENSG00000198904 | 22 | + | 37740208 | 15 | 12 | 7 | 8 | 1.180291363 | 0.847248426 | 0.60838129 |
| RPL22 | 60S ribosomal protein L22;Epstein-Barr virus small RNA-associated protein;EBER-associated p | 14.8 | P35268;Q6IBD1;Q7Z4W8;Q8WYP0;Q5 | PF01776 | Ribosomal L22e protein family | ENSG00000116251 | 1 | - | 6167668 | 6 | 9 | 10 | 14 | 1.183556801 | 0.844910864 | 0.610141499 |
| KRT6B;K6B;KRTL1 | Keratin, type II cytoskeletal 6B;Cytokeratin-6B;CK 6B;K6b keratin | 60.0 | P04259 | PF00038 | Intermediate filament protein | ENSG00000185479 | 12 | - | 51126704 | 5 | 7 | 11 | 8 | 1.190324391 | 0.840107123 | 0.613766936 |
| SCGB1D2;LIPHB | Secretoglobin family 1D member 2 precursor;Lipophilin-B | 9.9 | O95969 | PF01099 | Uteroglobin family | ENSG00000124935 | 11 | + | 61766300 | 5 | 1 | 1 | 1 | 1.194040314 | 0.837492661 | 0.615744652 |
| HSPA8;HSC70;HSP73;HSPA10 | Heat shock cognate 71 kDa protein;Heat shock 70 kDa protein 8;Heat shock 70kDa protein 8 i | 70.9 | P11142;A8K7Q2;Q53GZ6;Q96BE0;Q9 | PF00012 | Hsp70 protein | ENSG00000109971 | 11 | - | 122433411 | 26 | 29 | 27 | 18 | 1.197713132 | 0.834924468 | 0.617690444 |
| RPL15;EC45;TCBAP0781;RP11-511D14.1-001 | 60S ribosomal protein L15;Ribosomal protein L15 pseudogene 3 | 24.1 | P61313;Q5U0C0;Q8N6E1;Q5T6E0;Q6 | PF00827 | Ribosomal L15 | ENSG00000174748;ENSG0 | 3 | + | 23933643 | 1 | 1 | 0 | 1 | 1.208203886 | 0.827674875 | 0.62319924 |
| POLR3K;RPC11;My010 | DNA-directed RNA polymerase III subunit RPC10;RNA polymerase III subunit C10;DNA-directe | 12.3 | Q9Y2Y1 | PF02150;PF01096 | RNA polymerases M/15 Kd subunit;Transcription factor S | ENSG00000161980 | 16 | - | 37000 | 1 | 1 | 2 | 2 | 1.226427814 | 0.815376159 | 0.63259725 |
| SOD1 | Superoxide dismutase [Cu-Zn];Uncharacterized protein SOD1 | 16.1 | P00441;A6NHJ0;Q6NR85;A8MST3 | PF00080 | Copper/zinc superoxide dismutase (SODC) | ENSG00000142168 | 21 | + | 31953806 | 2 | 0 | 0 | 0 | 1.238341904 | 0.807531423 | 0.638624576 |
| DNAJC9 | DnaJ homolog subfamily C member 9;DnaJ protein SB73;RcDNAJ9 | 29.9 | Q8WXX5;Q2VIL4 | PF00226 | DnaJ domain | ENSG00000213551 | 10 | - | 74672596 | 6 | 11 | 6 | 6 | 1.251360962 | 0.799129932 | 0.645106388 |
| RPS3;OK/SW-cl.26 | 40S ribosomal protein S3 | 26.7 | P23396;Q53G83;Q6IPX9;Q9NQS8 | PF07650;PF00189 | KH domain;Ribosomal protein S3, C-terminal domain | ENSG00000149273 | 11 | + | 74788210 | 1 | 3 | 1 | 1 | 1.256535794 | 0.795838849 | 0.647652687 |
| KRT16;KRT16A | Keratin, type I cytoskeletal 16;Cytokeratin-16;CK-16;Keratin-16;K16 | 51.3 | P08779;A8K488 | PF00038 | Intermediate filament protein | ENSG00000186832 | 17 | - | 37019559 | 41 | 99 | 76 | 31 | 1.267557971 | 0.788918553 | 0.653019531 |
| TXN;TRDX;TRX;TRX1;TXN delta 3 | Thioredoxin;Trx;ATL-derived factor;ADF;Surface-associated sulphydryl protein;SASP;Thioredo | 11.7 | P10599;Q5T937;Q9UDG5;O60744;Q5 | PF00085 | Thioredoxin | ENSG00000136810 | 9 | - | 112046129 | 6 | 3 | 1 | 2 | 1.280429728 | 0.780987803 | 0.659190059 |
| KRT5 | Keratin, type II cytoskeletal 5;Cytokeratin-5;CK-5;Keratin-5;K5;58 kDa cytokeratin | 62.4 | P13647 | PF00038 | Intermediate filament protein | ENSG00000186081 | 12 | - | 51194630 | 44 | 54 | 46 | 24 | 1.283166799 | 0.779321909 | 0.660488804 |
| CSTA;STF1;STFA | Cystatin-A;Stefin-A;Cystatin-AS | 11.0 | P01040;Q6IB90 | PF00031 | Cystatin domain | ENSG00000121552 | 3 | + | 123526701 | 18 | 10 | 8 | 6 | 1.289921716 | 0.775240844 | 0.663674086 |
| DYNLL2;DLC2 | Dynein light chain 2, cytoplasmic;Dynein light chain LC8-type 2 | 10.4 | Q96FJ2 | PF01221 | Dynein light chain type 1 | ENSG00000121083 | 17 | + | 53515798 | 2 | 3 | 1 | 0 | 1.292000916 | 0.773993259 | 0.664648839 |
| KRT1;KRTA | Keratin, type II cytoskeletal 1;Cytokeratin-1;CK-1;Keratin-1;K1;67 kDa cytokeratin;Hair alpha | 66.0 | P04264 | PF00038 | Intermediate filament protein | ENSG00000167768 | 12 | - | 51354787 | 232 | 261 | 263 | 147 | 1.292030519 | 0.773975525 | 0.664662698 |
| KRT14 | Keratin, type I cytoskeletal 14;Cytokeratin-14;CK-14;Keratin-14;K14 | 51.6 | P02533 | PF00038 | Intermediate filament protein | ENSG00000186847 | 17 | - | 36992059 | 108 | 203 | 175 | 68 | 1.310289319 | 0.763190225 | 0.673108076 |
| TFAP2A;RP1-290I10.1-001;hCG_37170;AP2TF;TFAP2;RP1-290I10.1-003;RP1-290I10.1-005 | Transcription factor AP-2 alpha;Activating enhancer binding protein 2 alpha;Transcription fac | 48.3 | Q96SH1;P05549;Q5TAV5;Q8N1C6;Q9 | PF03299 | Transcription factor AP-2 | ENSG00000137203 | 6 | - | 10504902 | 3 | 2 | 3 | 3 | 1.312292442 | 0.762025268 | 0.674022191 |
| COL1A1 | Collagen alpha-1(I) chain precursor;Alpha-1 type I collagen | 138.9 | P02452;Q14042;Q6LAN8;Q9UMA6 | PF01410;PF01391;PF00093 | Fibrillar collagen C-terminal domain;Collagen triple helix | ENSG00000108821 | 17 | - | 45616456 | 1 | 0 | 0 | 0 | 1.313014257 | 0.761606353 | 0.674350991 |
| UBC;RPS27A;UBA80;UBCEP1;UBA52;UBCEP2;UBB;hCG_1998947 | cDNA FLJ75516, highly similar to Xenopus tropicalis ubiquitin C, mRNA;UBC protein;Ubiquit | 79.0 | A8K674;Q59EM9;Q66K58;Q96H31;Q9 | PF00240;PF01599;PF01020 | Ubiquitin family;Ribosomal protein S27a;Ribosomal L40 | ENSG00000150991;ENSG0 | 12 | - | 123962147 | 12 | 5 | 8 | 7 | 1.323345576 | 0.755660515 | 0.679022535 |
| IGKV1-5;IGKC;IGKV2-24;IGK@ | IGKV1-5 protein;Ig kappa chain V-II region RPMI 6410 precursor;Immunoblobulin light chain | 26.2 | Q6PIH6;P06310;Q8TCD0;Q0KKI6;P01 | PF07654;PF07686 | Immunoglobulin C1-set domain;Immunoglobulin V-set d | ENSG00000211614;ENSG0 | 2 | - | 89302292 | 9 | 0 | 0 | 0 | 1.329482521 | 0.752172356 | 0.681767078 |
| SERPINB3;SCCA;SCCA1;SERPINB4;SCCA2 | Serpin B3;Squamous cell carcinoma antigen 1;SCCA-1;Protein T4-A;SCCA1b;SCCA2/SCCA1 fus | 44.6 | P29508;Q53H28;Q5K684;Q86VF3;Q8 | PF00079 | Serpin (serine protease inhibitor) | ENSG00000206073;ENSG0 | 18 | - | 59473412 | 6 | 2 | 3 | 0 | 1.337856764 | 0.747464173 | 0.685475882 |
| DKFZp686P03159;ANXA2;ANX2;ANX2L4;CAL1H;LPC2D | Putative uncharacterized protein DKFZp686P03159;Annexin A2;Annexin-2;Annexin II;Lipoco | 40.4 | Q6N0B3;Q8TBV2;P07355;A6NK55;A6 | PF00191 | Annexin | ENSG00000182718;ENSG0 | 15 | - | 58426625 | 10 | 7 | 8 | 3 | 1.338963536 | 0.746846328 | 0.685962933 |
| EWSR1;hCG_2010995;EWS | CDNA FLJ31747 fis, clone NT2RI2007377, highly similar to RNA-BINDING PROTEIN EWS;Ewing | 69.0 | Q96MX4;Q01844;Q5THL0;Q96FE8;Q9 | PF00076;PF00641 | RNA recognition motif. (a.k.a. RRM, RBD, or RNP domain) | ENSG00000182944;ENSG0 | 22 | + | 27994017 | 3 | 3 | 0 | 1 | 1.343104875 | 0.744543496 | 0.687778957 |
| PIP;GCDFP15;GPIP4 | Prolactin-inducible protein precursor;Prolactin-induced protein;Secretory actin-binding pro | 16.6 | P12273;A0A963;A0A9C3;A0A9F3;A4D | PF05326 | Seminal vesicle autoantigen (SVA) | ENSG00000159763 | 7 | + | 142539288 | 16 | 6 | 3 | 7 | 1.368012799 | 0.730987313 | 0.698489303 |
| DSG1 | Desmoglein-1 precursor;Desmosomal glycoprotein 1;DG1;DGI;Pemphigus foliaceus antigen | 113.7 | Q02413;Q712V6 | PF00028;PF01049 | Cadherin domain;Cadherin cytoplasmic region | ENSG00000134760 | 18 | + | 27152050 | 22 | 19 | 31 | 4 | 1.393289409 | 0.717725975 | 0.708993064 |
| ACTG1;ACTB;ACTG;ACTA1 | Beta actin variant;Actin, cytoplasmic 2;Gamma-actin;Actin, cytoplasmic 1;Beta-actin;Unchar | 45.1 | Q53G76;Q53G99;Q53GK6;P63261;A1 | PF00022 | Actin | ENSG00000075624;ENSG0 | 7 | - | 5533315 | 8 | 4 | 2 | 11 | 1.442794093 | 0.693099594 | 0.728535596 |
| POLDIP3;KIAA1649;PDIP46 | Polymerase delta-interacting protein 3;46 kDa DNA polymerase delta interaction protein;p46 | 46.1 | Q9BY77;A8K6V9;Q659C6;Q96DI9;A8K | PF00076 | RNA recognition motif. (a.k.a. RRM, RBD, or RNP domain) | ENSG00000100227 | 22 | - | 41309671 | 0 | 2 | 0 | 0 | 1.485833865 | 0.673022754 | 0.744465535 |
| HNRPM;NAGR1 | Heterogeneous nuclear ribonucleoprotein M;hnRNP M | 77.5 | P52272;Q59ES8;Q7KYM9;Q6P2D7;Q8 | PF00076 | RNA recognition motif. (a.k.a. RRM, RBD, or RNP domain) | ENSG00000099783 | 19 | + | 8415651 | 38 | 50 | 36 | 34 | 1.496257531 | 0.668334147 | 0.748180951 |
| KRT17 | Keratin, type I cytoskeletal 17;Cytokeratin-17;CK-17;Keratin-17;K17;39.1;Radiated keratinoc | 48.1 | Q04695;Q14666;A8MW45 | PF00038 | Intermediate filament protein | ENSG00000186831;ENSG0 | 17 | - | 37029218 | 19 | 40 | 42 | 19 | 1.511951287 | 0.66139697 | 0.753672801 |
| LGALS7;PIG1 | Galectin-7;Gal-7;HKL-14;PI7;p53-induced gene 1 protein | 15.1 | P47929;Q6IB87 | PF00337 | Galactoside-binding lectin | ENSG00000178934;ENSG0 | 19 | + | 43971694 | 2 | 1 | 2 | 0 | 1.527872032 | 0.654505076 | 0.759120899 |
| DSC1;hCG_25474 | Desmocollin-1 precursor;Desmosomal glycoprotein 2/3;DG2/DG3;Desmocollin 1b;Desmoco | 100.0 | Q08554;Q9HB01;Q9HB00 | PF00028;PF08758 | Cadherin domain;Cadherin prodomain like | ENSG00000134765 | 18 | - | 26963214 | 15 | 10 | 22 | 2 | 1.528270755 | 0.654334317 | 0.759255771 |
| S100A7;PSOR1 | Protein S100-A7;S100 calcium-binding protein A7;Psoriasin | 11.5 | P31151;Q5SY67 | PF01023 | S-100/ICaBP type calcium binding domain | ENSG00000143556 | 1 | - | 151696844 | 13 | 9 | 3 | 1 | 1.539561681 | 0.649535522 | 0.763043512 |
| CASP14 | Caspase-14 precursor;CASP-14;Caspase-14 subunit p19;Caspase-14 subunit p10 | 27.7 | P31944 | PF00656 | Caspase domain | ENSG00000105141 | 19 | + | 15024015 | 5 | 3 | 2 | 1 | 1.550507319 | 0.644950196 | 0.766657843 |
| SYNGAP1;KIAA1938;DAB2IP;AF9Q34;AIP1;KIAA1743 | Ras GTPase-activating protein SynGAP;Synaptic Ras GTPase-activating protein 1;Synaptic Ras- | 148.3 | Q96PV0;A8MQC4;Q5VWQ8 | PF00168;PF00169;PF00616 | C2 domain;PH domain;GTPase-activator protein for Ras-l | ENSG00000197283;ENSG0 | 6 | + | 33495919 | 0 | 2 | 2 | 1 | 1.552413039 | 0.644158465 | 0.76728139 |
| RPL30 | Uncharacterized protein RPL30;60S ribosomal protein L30 | 12.9 | A8MYX5;P62888 | PF01248 | Ribosomal protein L7Ae/L30e/S12e/Gadd45 family | ENSG00000156482 | 8 | - | 99123130 | 1 | 2 | 2 | 0 | 1.553881988 | 0.643549515 | 0.767760874 |
| PRSS3;PRSS4;TRY3;TRY4 | Trypsin-3 precursor;Trypsin III;Brain trypsinogen;Mesotrypsinogen;Trypsin IV;Serine protease | 32.5 | P35030;A8CED1;A6NN76;Q5VXV0;Q8 | PF00089 | Trypsin | ENSG00000010438 | 9 | + | 33740515 | 1 | 0 | 0 | 0 | 1.564925966 | 0.639007865 | 0.771333792 |
| DEFA1;DEF1;DEFA2;MRS;DEFA3;DEF3 | Neutrophil defensin 1 precursor;HNP-1;HP-1;HP1;Defensin, alpha 1;HP 1-56;Neutrophil defe | 10.2 | P59665;Q6EZF6;P59666;Q6EZE9 | PF00323;PF00879 | Mammalian defensin;Defensin propeptide | ENSG00000206047;ENSG0 | 8 | - | 6822581 | 3 | 0 | 1 | 2 | 1.61143765 | 0.62056388 | 0.78577715 |
| KRT36;HHA6;HKA6;KRTHA6;hCG_1771519 | Keratin, type I cuticular Ha6;Hair keratin, type I Ha6;Keratin-36;KRT36 protein;Keratin, hair, a | 52.2 | O76013;Q86XG4 | PF00038 | Intermediate filament protein | ENSG00000126337 | 17 | - | 36895916 | 1 | 0 | 0 | 0 | 1.624536894 | 0.615560043 | 0.789674013 |
| HRNR;S100A18 | Hornerin | 282.4 | Q86YZ3;Q5DT20;Q5U1F4;Q5W8V9 | PF01023 | S-100/ICaBP type calcium binding domain | ENSG00000197915 | 1 | - | 150451176 | 38 | 39 | 36 | 19 | 1.629722071 | 0.613601557 | 0.791196376 |
| PRDX1;PAGA;PAGB;TDPX2 | Peroxiredoxin-1;Thioredoxin peroxidase 2;Thioredoxin-dependent peroxide reductase 2;Pro | 22.1 | Q06830 | PF00578 | AhpC/TSA family | ENSG00000117450 | 1 | - | 45749295 | 6 | 7 | 5 | 1 | 1.664802647 | 0.600671798 | 0.801202629 |
| THOC4;ALY;BEF | THO complex subunit 4;Tho4;Ally of AML-1 and LEF-1;Transcriptional coactivator Aly/REF;bZI | 27.6 | Q86V81;A8MU19 | PF00076 | RNA recognition motif. (a.k.a. RRM, RBD, or RNP domain) | ENSG00000183684 | 17 | - | 77439017 | 9 | 15 | 8 | 5 | 1.751612729 | 0.570902451 | 0.823888048 |
| RUVBL2;TIP48;TIP49B;CGI-46 | RuvB-like 2;48 kDa TATA box-binding protein-interacting protein;48 kDa TBP-interacting prot | 51.2 | Q9Y230 | PF00004;PF06068 | ATPase family associated with various cellular activities (A | ENSG00000183207 | 19 | + | 54188967 | 1 | 1 | 1 | 0 | 1.765354598 | 0.566458433 | 0.827224433 |
| GAPDH;GAPD;CDABP0047;OK/SW-cl.12 | Glyceraldehyde-3-phosphate dehydrogenase;GAPDH;Glyceraldehyde 3-phosphate dehydroge | 36.1 | P04406;Q2TSD0;Q53X65;A4UCT1;Q1 | PF02800;PF00044 | Glyceraldehyde 3-phosphate dehydrogenase, C-terminal | ENSG00000111640 | 12 | + | 6513872 | 6 | 12 | 6 | 1 | 1.837840733 | 0.54411679 | 0.843761204 |
| CALML5;CLSP | Calmodulin-like protein 5;Calmodulin-like skin protein | 15.9 | Q9NZT1;Q53H37;Q5SQI3 | PF00036 | EF hand | ENSG00000178372 | 10 | - | 5530660 | 8 | 8 | 10 | 3 | 1.847717372 | 0.541208312 | 0.845882373 |
| S100A9;CAGB;CFAG;MRP14 | Protein S100-A9;S100 calcium-binding protein A9;Calgranulin-B;Migration inhibitory factor- | 13.2 | P06702 | PF00036;PF01023 | EF hand;S-100/ICaBP type calcium binding domain | ENSG00000163220 | 1 | + | 151596954 | 12 | 10 | 7 | 7 | 1.870755157 | 0.534543495 | 0.85071311 |
| SKP1A;EMC19;OCP2;SKP1;TCEB1L | S-phase kinase-associated protein 1A;Cyclin A/CDK2-associated protein p19;p19A;p19skp1;R | 18.7 | P63208 | PF01466;PF03931 | Skp1 family, dimerisation domain;Skp1 family, tetrameri | ENSG00000113558 | 5 | - | 133520468 | 1 | 0 | 0 | 0 | 1.872177342 | 0.534137433 | 0.851006046 |
| PRDX2;NKEFB;TDPX1;hCG_172488 | Peroxiredoxin-2;Thioredoxin peroxidase 1;Thioredoxin-dependent peroxide reductase 1;Thic | 21.9 | P32119;A6NIW5 | PF00578 | AhpC/TSA family | ENSG00000167815 | 19 | - | 12768634 | 4 | 0 | 1 | 0 | 1.892509363 | 0.528398971 | 0.855128257 |
| C7orf24 | Uncharacterized protein C7orf24 | 21.0 | O75223 | | | ENSG00000006625 | 7 | - | 30502764 | 3 | 1 | 0 | 1 | 1.952351468 | 0.512202857 | 0.866575938 |
| S100A8;CAGA;CFAG;MRP8 | Protein S100-A8;S100 calcium-binding protein A8;Calgranulin-A;Migration inhibitory factor- | 10.8 | P05109;A8K5L3;Q9UCM6 | PF00036;PF01023 | EF hand;S-100/ICaBP type calcium binding domain | ENSG00000143546 | 1 | - | 151629132 | 8 | 6 | 1 | 10 | 1.966325809 | 0.508562719 | 0.869108547 |
| AZGP1;ZAG;ZNGP1 | Zinc-alpha-2-glycoprotein precursor;Zn-alpha-2-glycoprotein;Zn-alpha-2-GP | 34.3 | P25311;Q5XKQ4;Q8N4N0 | PF07654;PF00129 | Immunoglobulin C1-set domain;Class I Histocompatibilit | ENSG00000160862 | 7 | - | 99402290 | 15 | 7 | 2 | 1 | 2.017801078 | 0.495588991 | 0.878005601 |

| | | | | | | | | | | | | | | | | |
|---|---|---|---|---|---|---|---|---|---|---|---|---|---|---|---|---|
| KRT10;KPP | Keratin, type I cytoskeletal 10;Cytokeratin-10;CK-10;Keratin-10;K10 | 59.5 | P13645;A6NK80 | PF00038 | Intermediate filament protein | ENSG00000186395 | 17 | - | 36227896 | 180 | 164 | 154 | 65 | 2.035764895 | 0.491215858 | 0.880957003 |
| NOLA1;GAR1 | H/ACA ribonucleoprotein complex subunit 1;Nucleolar protein family A member 1;snoRNP p | 25.6 | Q9NY12 | PF04410 | Gar1 protein RNA binding region | ENSG00000109534 | 4 | + | 110956115 | 21 | 22 | 16 | 10 | 2.038870435 | 0.490467655 | 0.881459468 |
| ANXA1;ANX1;LPC1;RP11-71A24.1-004 | Annexin A1;Annexin-1;Annexin I;Lipocortin I;Calpactin II;Chromobindin-9;p35;Phospholipa | 38.7 | P04083;Q05BR2;Q5TZZ9;Q5T3N1 | PF00191 | Annexin | ENSG00000135046 | 9 | + | 74956495 | 5 | 0 | 0 | 0 | 2.098818024 | 0.476458649 | 0.890727753 |
| CAT | Catalase | 59.8 | P04040;A8K6C0;Q8TAK2;Q9UC85 | PF00199 | Catalase | ENSG00000121691 | 11 | + | 34417054 | 4 | 3 | 2 | 1 | 2.128612667 | 0.469789556 | 0.895043001 |
| KRT80;KB20 | Keratin, type II cytoskeletal 80;Cytokeratin-80;Keratin-80;Type II keratin-20 | 50.5 | Q6KB66;Q6P1A5;Q7Z3Q0 | PF00038 | Intermediate filament protein | ENSG00000167767 | 12 | - | 50849047 | 3 | 4 | 8 | 0 | 2.14333626 | 0.466562349 | 0.897107719 |
| HIST1H2BN;H2BFD;HIST2H2BF;HIST1H2BK;H2BFT;HIRIP1;HIST1H2BC;H2BFL;HIST1H2BE;H2E | Histone H2B type 1-N;H2B.d;H2B/d;Histone H2B type 2-F;Histone H2B type 1-K;H2B K;HIRA-i | 18.8 | Q99877;A8K9J7;Q5QNW6;A8K0U9;Q | PF00125 | Core histone H2A/H2B/H3/H4 | ENSG00000196501;ENSG0 | 6 | + | 27913517 | 2 | 4 | 3 | 4 | 2.160557301 | 0.462843545 | 0.899467532 |
| NIP7;CGI-37;HSPC031;HSPC180;OK/SW-cl.76;OK/SW-cl.78 | 60S ribosome subunit biogenesis protein NIP7 homolog;KD93 | 20.5 | Q9Y221 | | | ENSG00000132603 | 16 | + | 67931047 | 2 | 2 | 1 | 0 | 2.221541349 | 0.450137919 | 0.907367868 |
| HBA1;HBA2 | Hemoglobin subunit alpha;Hemoglobin alpha chain;Alpha-globin;Alpha-globin gene;alpha-th | 15.3 | P69905;Q1HDT5;Q4ZGM8;Q7Z6G4;Q | PF00042 | Globin | ENSG00000188536;ENSG0 | 16 | + | 162875 | 20 | 4 | 6 | 4 | 2.236160401 | 0.447195112 | 0.909160773 |
| IFPS;FLG2;RP1-14N1.3-001 | Ifapsoriasin;Filaggrin 2 | 248.1 | Q5D862 | PF00036;PF01023 | EF hand;S-100/ICaBP type calcium binding domain | ENSG00000143520 | 1 | - | 150587837 | 12 | 6 | 4 | 1 | 2.25290303 | 0.443871745 | 0.911168334 |
| FABP5 | Fatty acid-binding protein, epidermal;E-FABP;Fatty acid-binding protein 5;Psoriasis-associate | 15.2 | Q01469;A8MTQ5 | PF00061 | Lipocalin / cytosolic fatty-acid binding protein family | ENSG00000164687 | 8 | + | 82355325 | 2 | 6 | 3 | 0 | 2.267660659 | 0.440983088 | 0.91289827 |
| S100A14;S100A15 | Protein S100-A14;S100 calcium-binding protein A14;S114 | 11.7 | Q9HCY8 | PF01023 | S-100/ICaBP type calcium binding domain | ENSG00000189334 | 1 | - | 151853355 | 2 | 1 | 0 | 0 | 2.283397853 | 0.43794383 | 0.914703102 |
| SERPINB12 | SERPINB12 protein;Serpin B12 | 48.4 | Q3SYB4;Q96P63;Q3SYB5 | PF00079 | Serpin (serine protease inhibitor) | ENSG00000166634 | 18 | + | 59374373 | 7 | 9 | 7 | 1 | 2.338740633 | 0.427580547 | 0.920736532 |
| FLG | Filaggrin | 435.2 | P20930;O75370;Q03838 | PF03516;PF01023 | Filaggrin;S-100/ICaBP type calcium binding domain | ENSG00000143631 | 1 | - | 150541275 | 21 | 11 | 6 | 5 | 2.364858648 | 0.422858255 | 0.923422329 |
| SBSN;UNQ698/PRO1343 | Suprabasin precursor | 60.5 | A8K5J0;Q6UWP8 | | | ENSG00000189001 | 19 | - | 40706109 | 11 | 6 | 2 | 0 | 2.452331307 | 0.407775245 | 0.931723878 |
| KRT78;K5B;KB40;hCG_2039607 | Keratin, type II cytoskeletal 78;Cytokeratin-78;Keratin-78;Type II keratin-40;Keratin-5b;cDNA | 57.0 | Q7RTT2;A8K4D6;Q5HYM7;Q8N1N4 | PF00038 | Intermediate filament protein | ENSG00000170423 | 12 | - | 51517863 | 13 | 10 | 11 | 0 | 2.523116186 | 0.396335296 | 0.937728286 |
| KRT81;KRTHB1;MLN137;KRT86;KRTHB6;KRT85;KRTHB5;KRT83;KRTHB3 | Keratin type II cuticular Hb1;Type II hair keratin Hb1;ghHKb1;ghHb1;Keratin-81;K81;K2.9;M | 55.0 | Q14533;Q17R48;Q8N120;O43790;A8 | PF00038 | Intermediate filament protein | ENSG00000205426;ENSG0 | 12 | - | 50965966 | 0 | 9 | 0 | 1 | 2.568085518 | 0.389395132 | 0.941243517 |
| THYN1;THY28;HSPC144;MDS012;My0054 | Thymocyte nuclear protein 1;Thymocyte protein Thy28 | 25.7 | Q9P016 | PF01878 | Protein of unknown function DUF55 | ENSG00000151500 | 11 | - | 133623383 | 5 | 1 | 0 | 0 | 2.583259469 | 0.387107843 | 0.942380513 |
| HBB;HBD | Hemoglobin subunit beta;Hemoglobin beta chain;Beta-globin;LVV-hemorphin-7;Beta globin; | 16.0 | P68871;A1YZ08;A4GX73;Q14473;Q14 | PF00042 | Globin | ENSG00000188170 | 11 | - | 5203272 | 21 | 11 | 13 | 10 | 2.853676047 | 0.3504252 | 0.959097013 |
| DDX59;ZNHIT5;RP11-92G12.2-004 | Probable ATP-dependent RNA helicase DDX59;DEAD box protein 59;Zinc finger HIT domain-c | 68.8 | Q5T1V6;Q5T1W1 | PF00270;PF00271;PF04438 | DEAD/DEAH box helicase;Helicase conserved C-terminal d | ENSG00000118197 | 1 | - | 198859666 | 2 | 1 | 1 | 0 | 2.871374161 | 0.348265306 | 0.959989409 |
| GSDM1;GSDM;GSDMA;FKSG9 | Gasdermin;Gasdermin family member A | 49.4 | Q96QA5 | PF04598 | Gasdermin family | ENSG00000167914 | 17 | + | 35372752 | 1 | 0 | 0 | 0 | 2.896716659 | 0.345218438 | 0.96123044 |
| CDSN;S | Corneodesmosin precursor;S protein;Corneodesmosin;cDNA FLJ78714, highly similar to Hon | 51.5 | Q15517;Q7Z559;Q2L6G8;A8K9C1;Q0EFA5;Q7Z560 | | | ENSG00000206460;ENSG0 | c6_QBL | - | 31218141 | 4 | 3 | 3 | 1 | 2.931929894 | 0.341072275 | 0.962885557 |
| ACTA2;ACTSA;ACTVS;GIG46;ACTA1;ACTA;ACTC1;ACTC;ACTG2;ACTA3;ACTL3;ACTSG;RP5-1068 | Actin, aortic smooth muscle;Alpha-actin-2;Cell growth-inhibiting gene 46 protein;Actin, alph | 42.0 | P62736;A8K3K1;P68133;Q5T8M9;P6 | PF00022 | Actin | ENSG00000107796;ENSG0 | 10 | - | 90684814 | 1 | 1 | 0 | 0 | 3.224418967 | 0.310133395 | 0.973992969 |
| LYZ;LZM | Lysozyme C precursor;1,4-beta-N-acetylmuramidase C | 16.5 | P61626 | PF00062 | C-type lysozyme/alpha-lactalbumin family | ENSG00000090382 | 12 | + | 68028401 | 7 | 7 | 3 | 3 | 3.275297433 | 0.305315783 | 0.975523476 |
| ERH | Enhancer of rudimentary homolog | 12.3 | P84090 | PF01133 | Enhancer of rudimentary | ENSG00000100632 | 14 | - | 68916601 | 5 | 8 | 13 | 15 | 3.333558183 | 0.299979765 | 0.977155883 |
| LOC342897 | CDNA FLJ41947 fis, clone PLACE6019932, moderately similar to Ictalurus punctatus NCC rec | 30.8 | Q6ZVX7 | PF04300 | F-box associated region | ENSG00000188505 | 19 | + | 44379441 | 1 | 1 | 1 | 0 | 3.355949495 | 0.297978263 | 0.977751187 |
| PTGDS;RP11-229P13.6-003;RP11-229P13.6-005;PDS;RP11-229P13.6-004 | Prostaglandin D2 synthase 21kDa;Brain;Prostaglandin-H2 D-isomerase precursor;Lipocalin-ty | 24.4 | Q5SQ11;Q5SQ09;P41222;Q5SQ12 | PF00061 | Lipocalin / cytosolic fatty-acid binding protein family | ENSG00000107317 | 9 | + | 138991777 | 2 | 0 | 0 | 0 | 3.392810324 | 0.294740909 | 0.978694461 |
| GIG12;LTF;LF | cDNA FLJ78440, highly similar to Human lactoferrin;Lactotransferrin precursor;Lactoferrin;T | 78.4 | A8K494;A8K9U8;Q19KS2;Q5DSM0;Q5 | PF00405 | Transferrin | ENSG00000012223 | 3 | - | 46452503 | 20 | 5 | 4 | 1 | 3.41215184 | 0.293070193 | 0.979171795 |
| CTSD;CPSD | Cathepsin D precursor;Cathepsin D light chain;Cathepsin D heavy chain | 44.6 | P07339 | PF07966;PF00026 | A1 Propeptide;Eukaryotic aspartyl protease | ENSG00000117984 | 11 | - | 1730561 | 3 | 3 | 0 | 1 | 3.495746709 | 0.286061916 | 0.981104108 |
| DKFZp686K18196;IGHA1 | Putative uncharacterized protein DKFZp686K18196;IGHA1 protein;CDNA FLJ25298 fis, clone | 56.4 | Q6N092;Q9BRV0;Q8N5K4;Q96DK0;Q | PF07654;PF07686;PF00047 | Immunoglobulin C1-set domain;Immunoglobulin V-set d | ENSG00000211895 | 14 | - | 105244550 | 19 | 11 | 10 | 2 | 3.522480859 | 0.283890826 | 0.981679863 |
| DSC3;DSC4 | Desmocollin-3 precursor;Desmocollin-4;HT-CP | 100.0 | Q14574;A6NN35;A8K6T3;Q9HAX1 | PF00028;PF08758 | Cadherin domain;Cadherin prodomain like | ENSG00000134762 | 18 | - | 26825029 | 1 | 1 | 0 | 0 | 3.551339574 | 0.281583887 | 0.982279846 |
| PLUNC;LUNX;NASG;SPURT;UNQ787/PRO1606 | Protein Plunc precursor;Palate lung and nasal epithelium clone protein;Lung-specific protein | 26.7 | Q9NP55;A6XMV5;A8K9R3 | PF01273 | LBP / BPI / CETP family, N-terminal domain | ENSG00000198183 | 20 | + | 31287461 | 6 | 0 | 0 | 0 | 3.669221629 | 0.272537367 | 0.984516095 |
| DKFZp686C02218;DKFZp686M08189;IGH@;DKFZp686L19235;DKFZp686K04218;IGHA2;D | Putative uncharacterized protein DKFZp686C02218;CDNA FLJ41981 fis, clone SMINT201188 | 53.8 | Q7Z374;Q6ZVX0;Q6MZX9;Q6P089;Q6 | PF07654;PF07686 | Immunoglobulin C1-set domain;Immunoglobulin V-set d | ENSG00000211890 | 14 | - | 105121536 | 3 | 1 | 1 | 1 | 3.829285925 | 0.261145294 | 0.987071418 |
| MUC5AC;MUC5B;MUC5 | Mucin 5AC, oligomeric mucus/gel-forming;Mucin-5B precursor;Mucin-5 subtype B, tracheob | 648.8 | A7Y9J9;Q9HC84;A6NLP9;O00446;Q9 | PF00007;PF08742;PF01826;PF00093;PF | Cystine-knot domain;C8 domain;Trypsin Inhibitor like cy | ENSG00000117983 | 11 | + | 1200930 | 7 | 0 | 1 | 0 | 3.867832792 | 0.258542717 | 0.987615072 |
| ALB;GIG20;GIG42;PRO0903;PRO1708;PRO2044;PRO2619;PRO2675;UNQ696/PRO1341 | Serum albumin precursor;Uncharacterized protein ALB;ALB protein | 69.4 | P02768;A8K9P0;A6NBZ8;Q56G89;Q8 | PF00273 | Serum albumin family | ENSG00000163631 | 4 | + | 74488870 | 58 | 20 | 18 | 9 | 4.022289988 | 0.248614596 | 0.989554798 |
| APOD | Apolipoprotein D precursor;Apo-D;ApoD | 21.3 | P05090 | PF00061 | Lipocalin / cytosolic fatty-acid binding protein family | ENSG00000189058 | 3 | - | 196776867 | 3 | 2 | 3 | 3 | 4.069959626 | 0.245702683 | 0.990084038 |
| IGHG1;DKFZp686N02209;DKFZp686H20196;DKFZp686O01196;IGHM;IGHV4-31;DKFZp686 | Uncharacterized protein ENSP00000346720;IGHG1 protein;Putative uncharacterized protei | 51.6 | A6NG04;A8K008;Q6PJ95;Q7Z351;Q6 | PF07686;PF07654;PF00008;PF00594;PF | Immunoglobulin V-set domain;Immunoglobulin C1-set d | ENSG00000198229;ENSG0 | 16 | + | 33537182 | 6 | 2 | 1 | 1 | 4.167092888 | 0.239975452 | 0.991073575 |
| RPS21;RP5-908M14.8-003;hCG_41768 | Ribosomal protein S21;Ribosomal protein S21, isoform CRA_e;40S ribosomal protein S21 | 9.9 | Q9BYK1;P63220;Q6FGH5;Q8WVC2 | PF01249 | Ribosomal protein S21e | ENSG00000171858 | 20 | + | 60395500 | 1 | 1 | 0 | 1 | 4.342530517 | 0.230280477 | 0.992597143 |
| KRT13 | Keratin, type I cytoskeletal 13;Cytokeratin-13;CK-13;Keratin-13;K13 | 49.6 | P13646;A1A4E9;A8K2H9 | PF00038 | Intermediate filament protein | ENSG00000171401 | 17 | - | 36898973 | 21 | 4 | 0 | 0 | 4.406457194 | 0.226939683 | 0.993079242 |
| KRT2;KRT2A;KRT2E | Keratin, type II cytoskeletal 2 epidermal;Cytokeratin-2e;K2e;CK 2e;keratin-2 | 65.9 | P35908 | PF00038 | Intermediate filament protein | ENSG00000172867 | 12 | - | 51324610 | 190 | 84 | 77 | 41 | 4.472741379 | 0.223576531 | 0.99354295 |
| TMSB4X;TB4X;THYB4;TMSB4;TMSL2;RP11-339B21.6-001;hCG_1780554 | Thymosin beta-4;T beta 4;Fx;Hematopoietic system regulatory peptide;Seraspenide;Thymosi | 5.1 | A6NI30;P62328;A2VCK8;Q2KHM7;Q5 | PF01290 | Thymosin beta-4 family | ENSG00000205542;ENSG0 | X | + | 12903146 | 6 | 0 | 0 | 0 | 4.531867671 | 0.220659576 | 0.993927873 |
| MUCL1;SBEM;UNQ590/PRO1160 | Mucin-like protein 1 precursor;Small breast epithelial mucin;Protein BS106 | 9.0 | Q96DR8 | | | ENSG00000172551 | 12 | + | 53534566 | 3 | 1 | 1 | 0 | 4.78339276 | 0.209056636 | 0.995305718 |
| | | 23.0 | P02662 | | | | | | | 5 | 1 | 2 | 1 | 5.352833155 | 0.186816957 | 0.997317331 |
| KRT4;CYK4 | Keratin, type II cytoskeletal 4;Cytokeratin-4;CK-4;Keratin-4;K4 | 63.9 | P19013;Q6PIN2 | PF00038 | Intermediate filament protein | ENSG00000170477 | 12 | - | 51486600 | 32 | 21 | 17 | 18 | 5.514428674 | 0.181342449 | 0.997698658 |
| | | 69.3 | P02769 | | | | | | | 6 | 4 | 2 | 0 | 6.058340096 | 0.165061714 | 0.998603826 |
| | | 14.3 | P00698 | | | | | | | 1 | 0 | 0 | 0 | 6.739420852 | 0.148380702 | 0.999228659 |
| KRT77;KRT1B | Keratin, type II cytoskeletal 1b;K1B;Keratin-77 | 61.8 | Q7Z794;Q0IIN1;Q147W7 | PF00038 | Intermediate filament protein | ENSG00000189182 | 12 | - | 51369910 | 13 | 2 | 0 | 0 | 6.745117227 | 0.148255392 | 0.999232369 |
| KRT31;HHA1;HKA1;KRTHA1;hCG_41569;KRT33B;HHA3-II;HKA3B;KRTHA3B;KRT35;HHA5;HK | Keratin, type I cuticular Ha1;Hair keratin, type I Ha1;Keratin-31;Type I hair keratin 1;Keratin, | 50.7 | Q15323;Q16275;Q9UE12;Q14525;Q9 | PF00038 | Intermediate filament protein | ENSG00000094796;ENSG0 | 17 | - | 36803505 | 0 | 2 | 0 | 0 | 7.033987364 | 0.142166875 | 0.999397156 |
| PIGR | Polymeric immunoglobulin receptor precursor;Poly-Ig receptor;PIGR;Hepatocellular carcino | 83.3 | P01833;Q8TE27 | PF07686 | Immunoglobulin V-set domain | ENSG00000162896 | 1 | - | 205168495 | 11 | 1 | 0 | 0 | 8.614654456 | 0.116081267 | 0.999824186 |
| | | 18.3 | P02754 | | | | | | | 9 | 16 | 3 | 1 | 9.013794948 | 0.110941064 | 0.999868432 |
| KRT37;HHA7;HKA7;KRTHA7;KRT38;HHA8;HKA8;KRTHA8 | Keratin, type I cuticular Ha7;Hair keratin, type I Ha7;Keratin-37;Keratin, type I cuticular Ha8; | 53.1 | O76014;O76015;A2RRM5;Q6A164;Q | PF00038 | Intermediate filament protein | ENSG00000108417;ENSG0 | 17 | - | 36830335 | 3 | 0 | 0 | 0 | 9.591345757 | 0.104260656 | 0.999912353 |
| XP32;C1orf68 | Skin-specific protein 32 | 26.2 | Q5T750 | | | ENSG00000198854 | 1 | + | 150957706 | 2 | 0 | 2 | 0 | 10.92940007 | 0.09149633 | 0.999963903 |
| UMOD | Uromodulin precursor;Tamm-Horsfall urinary glycoprotein;THP | 73.6 | P07911 | PF00008;PF07645;PF00100 | EGF-like domain;Calcium binding EGF domain;Zona pellu | ENSG00000169344 | 16 | - | 20251874 | 3 | 0 | 0 | 0 | 11.4087274 | 0.087652195 | 0.999973294 |
| | | 68.6 | Q3SZ57 | | | | | | | 2 | 2 | 3 | 3 | 12.63164687 | 0.079166241 | 0.999987184 |
| IGL@;V1-19;IGLC1;IGLC2;IGLC3;IGLV3-25;V2-17;V1-4;IGLV4-3;V5-1;IGLV2-14 | IGL@ protein;V1-19 protein;Putative uncharacterized protein;Ig lambda chain C regions;V2- | 24.8 | Q6IPQ0;Q6PIQ7;A2MYD2;A2NUT2;A2 | PF07654;PF07686 | Immunoglobulin C1-set domain;Immunoglobulin V-set d | ENSG00000211671;ENSG0 | 22 | + | 21495315 | 6 | 1 | 2 | 0 | 12.77628305 | 0.078270026 | 0.999988214 |
| IGL@;DKFZp781M0386;V<lambda>3;IGLC1;C1 segment;hCG_2040025;C7;hCG_2010697 | Putative uncharacterized protein;IGL@ protein;Putative uncharacterized protein DKFZp781 | 25.1 | Q6DHW4;Q96E61;Q6GMV7;Q6P5S3;Q | PF07654;PF07686 | Immunoglobulin C1-set domain;Immunoglobulin V-set d | ENSG00000211675;ENSG0 | 22 | + | 21567555 | 2 | 0 | 0 | 0 | 14.72411695 | 0.067915788 | 0.999995969 |
| AMBP;HCP;ITIL | AMBP protein precursor;Alpha-1-microglobulin;Protein HC;Complex-forming glycoprotein h | 39.0 | P02760;P78492;Q5TBD7 | PF00014;PF00061 | Kunitz/Bovine pancreatic trypsin inhibitor domain;Lipoc | ENSG00000106927 | 9 | - | 115862231 | 2 | 1 | 4 | 0 | 23.12172091 | 0.043249376 | 0.999999907 |
| HNRPR;HNRNPR;RP5-1057J7.2-002;DKFZp686A13234;SYNCRIP;HNRPQ;NSAP1 | HNRPR protein;Heterogeneous nuclear ribonucleoprotein R;hnRNP R;Putative uncharacteriz | 71.3 | Q9BV64;O43390;Q6MZS5;Q0VGD6;Q | PF00076 | RNA recognition motif. (a.k.a. RRM, RBD, or RNP domain | ENSG00000125944;ENSG0 | 1 | - | 23508863 | 6 | 7 | 0 | 0 | Infinity | #VALUE! | 1 |
| AMY1A;AMY1;AMY1B;AMY1C;AMY2B;AMY2A | Alpha-amylase 1 precursor;1,4-alpha-D-glucan glucanohydrolase 1;Salivary alpha-amylase;Al | 57.8 | P04745;A8K8H6;Q53F26;Q6NSB3;P1 | PF00128;PF02806 | Alpha amylase, catalytic domain;Alpha amylase, C-termi | ENSG00000051415;ENSG0 | 1 | + | 103999825 | 5 | 2 | 0 | 0 | Infinity | #VALUE! | 1 |
| C9orf114 | Uncharacterized protein C9orf114 | 42.0 | Q5T280 | PF02598 | Uncharacterized ACR, COG2106 | ENSG00000198917 | 9 | - | 130621753 | 1 | 1 | 0 | 0 | Infinity | #VALUE! | 1 |
| HSPA5;GRP78 | 78 kDa glucose-regulated protein precursor;GRP 78;Heat shock 70 kDa protein 5;Immunoglo | 72.4 | P11021;Q2KHP4 | PF00012 | Hsp70 protein | ENSG00000044574 | 9 | - | 127036955 | 3 | 1 | 2 | 0 | Infinity | #VALUE! | 1 |
| PKM2;PK2;PK3;PKM | Pyruvate kinase isozymes M1/M2;Pyruvate kinase muscle isozyme;Pyruvate kinase 2/3;Cytos | 57.9 | P14618;A6NFK3;Q504U3;Q8WUW7;Q | PF00224;PF02887 | Pyruvate kinase, barrel domain;Pyruvate kinase, alpha/be | ENSG00000067225 | 15 | - | 70278424 | 0 | 2 | 0 | 0 | Infinity | #VALUE! | 1 |
| SCGB2A1;LIPHC;MGB2;UGB3 | Mammaglobin-B precursor;Mammaglobin-2;Lipophilin-C;Lacryglobin;Secretoglobin family 2 | 10.9 | O75556 | PF01099 | Uteroglobin family | ENSG00000124939 | 11 | + | 61732716 | 2 | 0 | 0 | 0 | Infinity | #VALUE! | 1 |
| LCN1;VEGP;RP11-430N14.1-001 | Lipocalin-1 precursor;Von Ebner gland protein;VEG protein;Tear prealbumin;TP;Tear lipocali | 19.3 | P31025;Q5T8A1;Q5VSP4 | PF00061 | Lipocalin / cytosolic fatty-acid binding protein family | ENSG00000160349;ENSG0 | 9 | + | 137553105 | 3 | 2 | 0 | 0 | Infinity | #VALUE! | 1 |
| | | 19.0 | P02668 | | | | | | | 4 | 3 | 0 | 0 | Infinity | #VALUE! | 1 |
| PPP1CB | Serine/threonine-protein phosphatase PP1-beta catalytic subunit;PP-1B | 41.4 | P62140 | PF00149 | Calcineurin-like phosphoesterase | ENSG00000213639 | 2 | + | 28828118 | 1 | 4 | 0 | 0 | Infinity | #VALUE! | 1 |
| ATF1 | Cyclic AMP-dependent transcription factor ATF-1;Activating transcription factor 1;TREB36 pr | 29.2 | P18846 | PF00170;PF02173 | bZIP transcription factor;pKID domain | ENSG00000123268 | 12 | + | 49444128 | 0 | 2 | 0 | 0 | Infinity | #VALUE! | 1 |
| | Uncharacterized protein ENSP00000374816;Ig kappa chain V-III region CLL precursor;Rheum | 14.1 | A6NGI0;A6NK76;P04207;P04434;Q9 | PF07686 | Immunoglobulin V-set domain | ENSG00000211636;ENSG0 | 2 | + | 89911009 | 1 | 0 | 0 | 0 | Infinity | #VALUE! | 1 |
| | | 23.6 | P02666 | | | | | | | 2 | 0 | 0 | 0 | Infinity | #VALUE! | 1 |
| CPVL;VLP;PSEC0124;UNQ197/PRO223 | Probable serine carboxypeptidase CPVL precursor;Carboxypeptidase, vitellogenic-like;Vitello | 54.2 | Q9H3G5;A4D1A4;O75225;Q75MM4;Q | PF00450 | Serine carboxypeptidase | ENSG00000106066 | 7 | - | 29001374 | 2 | 1 | 0 | 0 | Infinity | #VALUE! | 1 |
| | | 15.2 | P01966 | | | | | | | 0 | 0 | 0 | 1 | NaN | #VALUE! | 1 |
| HCFC2 | Host cell factor 2;HCF-2;C2 factor | 86.8 | Q9Y5Z7;Q53FE7 | PF00041;PF01344;PF07646 | Fibronectin type III domain;Kelch motif;Kelch motif | ENSG00000111727 | 12 | + | 102982439 | 0 | 0 | 1 | 0 | NaN | #VALUE! | 1 |
| ACT | Actin-like protein | 42.0 | Q562R1;Q562R2;Q562S9;Q562X8 | PF00022 | Actin | ENSG00000169067 | 5 | - | 56813161 | 1 | 0 | 0 | 0 | NaN | #VALUE! | 1 |
| SDNK1 | Kappa 1 light chain variable region;BRE;Ig kappa chain V-I region WEA;Ig kappa chain V-I regio | 12.7 | Q96PF6;A2NI60;P01610;P80362;P01 | PF07686 | Immunoglobulin V-set domain | | | | | 1 | 0 | 0 | 0 | NaN | #VALUE! | 1 |
| NDRG3;RP3-460J8.1-003;hCG_1785125 | NDRG family member 3;NDRG family member 3, isoform CRA_c;Protein NDRG3 | 42.9 | Q5TH30;Q9UGV2;A2A2S8;A6NP49 | PF03096 | Ndr family | ENSG00000101079 | 20 | - | 34713583 | 0 | 0 | 0 | 1 | NaN | #VALUE! | 1 |
| CCDC72;HSPC016;HSPC330 | Coiled-coil domain-containing protein 72 | 7.6 | Q9Y2S6 | PF09072 | Translation machinery associated TMA7 | ENSG00000186012 | | | | 0 | 0 | 1 | 0 | NaN | #VALUE! | 1 |
| IPO7;RANBP7 | Importin-7;Imp7;Ran-binding protein 7;RanBP7 | 119.7 | O95373;A6NNM5 | PF03810 | Importin-beta N-terminal domain | ENSG00000205339 | 11 | + | 9362780 | 1 | 0 | 0 | 0 | NaN | #VALUE! | 1 |
| NOL12 | Nucleolar protein 12 | 24.7 | Q9UGY1 | | | ENSG00000100101 | 22 | + | 36412303 | 0 | 0 | 0 | 1 | NaN | #VALUE! | 1 |
| CREM;RP11-324I22.1-014;hCREM-1;RP11-324I22.1-005;RP11-324I22.1-015 | cAMP-responsive element modulator;Inducible cAMP early repressor;ICER;cAMP responsive e | 37.0 | Q03060;Q59FC7;Q5T9H8;Q14501;Q5 | PF00170;PF02173 | bZIP transcription factor;pKID domain | ENSG00000095794 | 10 | + | 35455807 | 0 | 1 | 0 | 0 | NaN | #VALUE! | 1 |
| S100P;S100E | Protein S100-P;S100 calcium-binding protein P | 10.4 | P25815;Q5J7W2 | PF01023 | S-100/ICaBP type calcium binding domain | ENSG00000163993 | 4 | + | 6745697 | 0 | 0 | 0 | 1 | NaN | #VALUE! | 1 |
| YWHAG;YWHAB;YWHAH;YWHA1;YWHAQ;LL22NC03-44A4.1-004;RP1-148E22.1-004;RP1-14 | 14-3-3 protein gamma;Protein kinase C inhibitor protein 1;KCIP-1;14-3-3 protein beta/alpha | 28.3 | P61981;P31946;A8K9K2;Q04917;Q9 | PF00244 | 14-3-3 protein | ENSG00000170027;ENSG0 | 7 | - | 75794053 | 0 | 1 | 0 | 0 | NaN | #VALUE! | 1 |
| SPP1;BNSP;OPN;PSEC0156 | Osteopontin precursor;Bone sialoprotein 1;Secreted phosphoprotein 1;SPP-1;Urinary stone | 35.4 | P10451;A6XMV6;Q4W597;Q3LGB0;Q | PF00865 | Osteopontin | ENSG00000118785 | 4 | + | 89115826 | 1 | 0 | 0 | 0 | NaN | #VALUE! | 1 |
| RBBP5;hCG_16408;RBQ3 | cDNA FLJ75510, highly similar to Homo sapiens retinoblastoma binding protein 5, mRNA;Re | 59.2 | A8K272;Q7Z6D8;Q15291;A8MTG7;A8 | PF00400 | WD domain, G-beta repeat | ENSG00000117222 | 1 | - | 203314400 | 0 | 0 | 0 | 2 | NaN | #VALUE! | 1 |
| FAM32A;CGI-144 | Protein FAM32A | 13.2 | Q9Y421 | PF08555 | Eukaryotic family of unknown function (DUF1754) | ENSG00000105058 | 19 | + | 16157235 | 0 | 0 | 1 | 0 | NaN | #VALUE! | 1 |
| DDX17 | Probable ATP-dependent RNA helicase DDX17;DEAD box protein 17;RNA-dependent helicase | 80.3 | Q92841;Q59F66 | PF00270;PF00271 | DEAD/DEAH box helicase;Helicase conserved C-terminal d | ENSG00000100201 | 22 | - | 37209389 | 0 | 0 | 1 | 0 | NaN | #VALUE! | 1 |
| C7orf50;FP15621 | Uncharacterized protein C7orf50 | 22.1 | Q9BRJ6 | | | ENSG00000146540 | 7 | - | 1003148 | 0 | 0 | 1 | 0 | NaN | #VALUE! | 1 |
| POLR1D | DNA-directed RNA polymerases I and III subunit RPAC2;RNA polymerases I and III subunit AC2 | 15.2 | Q9Y2S0 | PF01193 | RNA polymerase Rpb3/Rpb11 dimerisation domain | ENSG00000186184 | 13 | + | 27092903 | 0 | 0 | 1 | 0 | NaN | #VALUE! | 1 |
| UNQ773/PRO1567;LOC124220;hCG_1788537 | Uncharacterized protein UNQ773/PRO1567 precursor;Uncharacterized protein ENSP000003 | 22.7 | Q96DA0;A6NIY1 | | | ENSG00000162078 | 16 | + | 2820174 | 1 | 0 | 0 | 0 | NaN | #VALUE! | 1 |
| CDKN2AIPNL;MGC13017;hCG_16596 | CDKN2AIPNL protein;Similar to RIKEN cDNA A430101B06 | 13.2 | Q8WVE3;Q96HQ2 | | | ENSG00000145835 | 5 | - | 133756202 | 1 | 0 | 0 | 0 | NaN | #VALUE! | 1 |
| MEF2D;DAZAP1 | MEF2D protein;DAZ-associated protein 1;Deleted in azoospermia-associated protein 1;MEF2D | 50.4 | Q05BX2;Q5IRN4;Q96EP5;Q5IRM7 | PF00319;PF00076 | SRF-type transcription factor (DNA-binding and dimerisat | ENSG00000116604;ENSG0 | 1 | - | 154700146 | 0 | 0 | 0 | 1 | NaN | #VALUE! | 1 |
| APITD1;CENPS | Centromere protein S;CENP-S;Apoptosis-inducing TAF9-like domain-containing protein 1;Un | 19.0 | Q8N2Z9;A6NIJ4 | | | ENSG00000175279 | 1 | + | 10412746 | 0 | 0 | 1 | 0 | NaN | #VALUE! | 1 |
| HMGB2;HMG2 | High mobility group protein B2;High mobility group protein 2;HMG-2 | 24.0 | P26583;Q5U071 | PF00505 | HMG (high mobility group) box | ENSG00000164104 | 4 | - | 174489434 | 1 | 0 | 0 | 0 | NaN | #VALUE! | 1 |
| CCT3;CCTG;TRIC5;RP11-443G18.6-002 | T-complex protein 1 subunit gamma;TCP-1-gamma;CCT-gamma;hTRiC5;Chaperonin containi | 60.5 | P49368;Q2TU64;Q59H77;Q5SZY0;A6 | PF00118 | TCP-1/cpn60 chaperonin family | ENSG00000163468 | 1 | - | 154545376 | 1 | 0 | 0 | 0 | NaN | #VALUE! | 1 |
| DKFZp686I0180;PSMB5;LMPX | Putative uncharacterized protein DKFZp686I0180;Proteasome subunit beta type-5 precursor | 28.8 | Q7Z3B5;P28074;Q9TNN9 | PF00227 | Proteasome A-type and B-type | ENSG00000100804 | 14 | - | 22564907 | 1 | 0 | 0 | 0 | NaN | #VALUE! | 1 |
| ZCCHC9 | Zinc finger CCHC domain-containing protein 9 | 30.9 | Q8N567;Q05DH8 | PF00098 | Zinc knuckle | ENSG00000131732 | 5 | + | 80633178 | 0 | 1 | 0 | 0 | NaN | #VALUE! | 1 |
| IGK@;IGKV1-5;IGKC | IGK@ protein;IGKV1-5 protein;IGKC protein | 25.8 | Q6P5S8;Q6GMW0;Q5I69I9 | PF07654;PF07686 | Immunoglobulin C1-set domain;Immunoglobulin V-set d | ENSG00000211610;ENSG0 | 2 | - | 89223170 | 1 | 0 | 0 | 0 | NaN | #VALUE! | 1 |
| ALDOA;ALDA | Fructose-bisphosphate aldolase A;Muscle-type aldolase;Lung cancer antigen NY-LU-1 | 45.3 | P04075;A4UCS9;A4UCT0;Q6FI10;Q9 | PF00274 | Fructose-bisphosphate aldolase class-I | ENSG00000149925 | 16 | + | 29971945 | 0 | 1 | 0 | 0 | NaN | #VALUE! | 1 |
| IGHM | IGHM protein;Ig mu chain C region membrane-bound segment;Ig mu chain C region | 68.6 | Q96AA6;Q96EY0;P20769;A2J1N3;Q5 | PF07654;PF07686 | Immunoglobulin C1-set domain;Immunoglobulin V-set d | ENSG00000211899 | 14 | - | 105389522 | 2 | 0 | 0 | 0 | NaN | #VALUE! | 1 |

| | | | | | | | | | | | | | | | | |
|---|---|---|---|---|---|---|---|---|---|---|---|---|---|---|---|---|
| KRT19 | Keratin, type I cytoskeletal 19;Cytokeratin-19;CK-19;Keratin-19;K19 | 44.1 | P08727;A6NFZ1 | PF00038 | Intermediate filament protein | ENSG00000171345 | 17 | - | 36933396 | 1 | 0 | 0 | 0 | NaN | #VALUE! | 1 |
| HIST2H2BA;HIST2H3PS2 | Histone H2B | 7.6 | A6NG02;Q3KP43;Q6GMR5 | PF00125 | Core histone H2A/H2B/H3/H4 | ENSG00000158373;ENSG0 | 6 | + | 26266328 | 0 | 0 | 1 | 0 | NaN | #VALUE! | 1 |
| V<kappa>3 | Rheumatoid factor D5 light chain;Uncharacterized protein ENSP00000374812;Ig kappa chai | 12.8 | A0N5G5;A6NJ14;P04433 | PF07686 | Immunoglobulin V-set domain | ENSG00000211632;ENSG0 | 2 | + | 89849045 | 1 | 0 | 0 | 0 | NaN | #VALUE! | 1 |
| RPAP3 | RNA polymerase II-associated protein 3 | 75.7 | Q9H6T3 | PF00515 | Tetratricopeptide repeat | ENSG00000005175 | 12 | - | 46343344 | 0 | 0 | 1 | 0 | NaN | #VALUE! | 1 |
| | | 84.1 | A6NFJ0 | | | | | | | 0 | 0 | 1 | 0 | NaN | #VALUE! | 1 |

**Table S2.** Erythroid-related traits used to search for MAZ variants in the GeneATLAS database

| ID of the trait | Description of the trait | Category |
|---|---|---|
| selfReported_n_1504 | anaemia | Binary |
| clinical_c_D50 | D50 Iron deficiency anaemia | Binary |
| clinical_c_Block_D50-D53 | D50-D53 Nutritional anaemias | Binary |
| clinical_c_D51 | D51 Vitamin B12 deficiency anaemia | Binary |
| clinical_c_Block_D60-D64 | D60-D64 Aplastic and other anaemias | Binary |
| clinical_c_D63 | D63 Anaemia in chronic diseases classified elsewhere | Binary |
| clinical_c_D64 | D64 Other anaemias | Binary |
| clinical_c_Block_D70-D77 | D70-D77 Other diseases of blood and blood-forming organs | Binary |
| clinical_c_D75 | D75 Other diseases of blood and blood-forming organs | Binary |
| 30030-0.0 | Haematocrit percentage | Non-Binary |
| selfReported_n_1502 | haematology | Binary |
| 30020-0.0 | Haemoglobin concentration | Non-Binary |
| 30300-0.0 | High light scatter reticulocyte count | Non-Binary |
| 30290-0.0 | High light scatter reticulocyte percentage | Non-Binary |
| 30280-0.0 | Immature reticulocyte fraction | Non-Binary |
| 30050-0.0 | Mean corpuscular haemoglobin | Non-Binary |
| 30060-0.0 | Mean corpuscular haemoglobin concentration | Non-Binary |
| 30260-0.0 | Mean reticulocyte volume | Non-Binary |
| 30270-0.0 | Mean sphered cell volume | Non-Binary |

| | | |
|---|---|---|
| 30170-0.0 | Nucleated red blood cell count | Non-Binary |
| 30230-0.0 | Nucleated red blood cell percentage | Non-Binary |
| 30010-0.0 | Red blood cell (erythrocyte) count | Non-Binary |
| 30070-0.0 | Red blood cell (erythrocyte) distribution width | Non-Binary |
| 30250-0.0 | Reticulocyte count | Non-Binary |
| 30240-0.0 | Reticulocyte percentage | Non-Binary |

| **Cell line** | **Unique peaks** | **Total number of peaks** | **% unique peaks** |
|---|---|---|---|
| HepG2 | 3636 | 15481 | 23.5% |
| Gm12878 | 9480 | 23951 | 39.6% |
| A549 | 335 | 4323 | 7.7% |
| MCF-7 | 6512 | 20419 | 31.9% |
| IMR90 | 11383 | 28208 | 40.4% |
| Ery | 1780 | 10088 | 17.6% |

**Suppl Table 3.** Tissue specific MAZ peaks in ENCODE human cell lines and human primary erythroid cells.

**Table S4. Genomic loci associated with erythroid traits and erythropoiesis which are bound by MAZ**

| *Genomic loci* | *Erythroid trait*[1] | *Location of the MAZ peak* | *Position of the MAZ peak* | *Fold change relative to the input* | *$log_{10}$ q-value of the MAZ peak* |
|---|---|---|---|---|---|
| ITFG3(FAM234A) | Hb, MCH, MCHC, MCV, RBC | ITFG3 promoter | chr16:234379-235136 | 36.8 | 81.46 |
| HBS1L-MYB | Hct, MCH, MCHC, MCV, RBC | HBSL1 promoter,<br>MYB first intron | chr6:135054682-135055128<br>chr6:135183043-135183281 | 26.95<br>7.15 | 57.41<br>8.84 |
| G6PD | Hct, Hb, MCV, RBC, RDW | G6PD promoter,<br>G6PD second intron | chrX:154547246-154547697<br>chrX:154541913-154542409 | 13.49<br>16.38 | 21.39<br>29.99 |
| CCND3 | MCH, MCV, RBC | CCND3 promoter,<br>CCND3 first intron,<br>CCND3 second intron | chr6:42045709-42046059<br>chr6:42030970-42031193<br>chr6:42017103-42017413<br>chr6:41941500-41942068<br>chr6:41938239-41938465 | 20.64<br>9.47<br>18.51<br>35.9<br>6.48 | 38.02<br>12.15<br>32.58<br>78.1<br>6.64 |
| TFR2 | Hct, MCV, RBC | TFR2 promoter | chr7:100641422-100642132 | 78.49 | 205.83 |
| TFRC | MCH, MCV | TFRC promoter | chr8:20197105-20197542 | 13.49 | 21.39 |
| PDGFRA-KIT | MCV, RBC | KIT promoter | chr4:54657841-54658018 | 6.68 | 7.18 |
| CITED2 | MCH, MCV | CITED2 promoter | chr6:139374652-139374972<br>chr6:139375180-139375707 | 6.16<br>8.01 | 5.72<br>9.22 |
| HMOX2 | MCH, MCV | HMOX2 promoter | chr16:4476227-4476485 | 9.47 | 12.15 |
| PRKCE | Hct | PRKCE first intron, | chr2:45650249-45650426 | 6.01 | 6.17 |

| | | | | | |
|---|---|---|---|---|---|
| | | PRKCE second intron | chr2:45778683-45778969 | 6.55 | 6.46 |
| CD164 | MCH | CD164 promoter | chr6:109381892-109382875 | 7.91 | 9.22 |
| SH2B3 | Hb | SH2B3 promoter, SH2B3 first intron, SH2B3 second intron, SH2B3 third intron | chr12:111405191-111405978 chr12:111406092-111406300 chr12:111406596-111406773 chr12:111429666-111429889 chr12:111437838-111438136 | 13.18 8.68 7.91 6.17 6.55 | 21.95 10.7 9.22 5.88 6.46 |
| CDT1 | MCHC | CDT1 promoter | chr16:88803519-88804082 | 22.76 | 43.54 |
| FBXO7 | MCV, MCH | FBXO7 promoter | chr22:32474726-32475349 | 6.33 | 6.25 |
| RPS19 | Hb, RBC, MCV | RPS19 promoter | chr19:41859549-41860329 | 10.86 | 18.19 |
| EPOR | RBC, Hb | EPOR promoter, EPOR first intron | chr19:11384246-11384607 chr19:11383132-11383772 | 21.92 19.35 | 41.42 35.01 |
| KLF1 | MCH | KLF1 promoter, KLF1 first intron, KLF1 second intron | chr19:12887097-12887963 chr19:12886160-12886497 chr19:12885126-12885353 | 17.2 14.2 6.01 | 29.7 22.61 5.55 |
| GATA1 | RBC | GATA1 promoter | chrX:48786184-48786580 | 11.57 | 16.63 |
| SBDS | Hb | SBDS promoter | chr7:66995478-66995773 | 12.9 | 19.59 |
| CDAN1 | Hb | CDAN1 promoter | chr15:42736973-42737392 | 14.33 | 23.29 |
| PTPLAD1(HACD3) | MCH | PTPLAD1 promoter | chr15:65530326-65530535 | 7.7 | 8.62 |

[1] The erythroid traits are based on [28].
MCH: mean corpuscular hemoglobin; MCV: mean corpuscular volume; RBC: red blood cell count; Hb: hemoglobin; Hct: hematocrit; MCHC: MCH concentration; RDW: RBC distribution width.

**Table S5 Sequences of oligonucleotides used in this study**

The boxes denote the consensus MAZ binding sites. The substituted bases in the mutated oligonucleotides are given in bold and underlined. WT, wild type.

| Oligo | Sequence |
|---|---|
| 13/14 (WT) | 5′-CAG GCC CCG CCC GGG ACT-3′ |
| 13/14 (M1) | 5′-CAG GCC CCG CCC GG**<u>T GTG</u>**-3′ |
| 13/14 (M2) | 5′-CAG **<u>TAT A</u>**CG CCC GGG ACT-3′ |
| 13/14 (M3) | 5′-CAG GCC C**<u>A</u>**G CCC GGG ACT-3′ |
| 13/14 (M4) | 5′-CAG GCC CCG C**<u>A</u>**C GGG ACT-3′ |
| 11/12 (WT) | 5′-C CAG GCC **<u>G</u>**CG CCC CGG GC-3′ |

**Table S6: Datasets used in the study**

| Cell type | Data set | Crosslinking/ Reference | Mapped reads | GEO accession |
|---|---|---|---|---|
| Erythroblasts | MAZ | EGS + $CH_2O$ | 12 058 863 | GSE |
| Erythroblasts, default input data set | Input 1 | 20 | 13 158 630 | GSE114084 |
| Erythroblasts | ATAC-Seq | 8 | 70 014 192 | GSE74912 |
| Erythroblasts | H3K4me3 | 18 | 5 349 306 | GSE36985 |
| Erythroblasts | H3K27ac | 19 | 56 439 919 | GSE70660 |
| Erythroblasts | Pol2 | 18 | 97 237 763 | GSE36985 |
| Erythroblasts | H3K27me3 | 18 | 4 990 983 | GSE36985 |
| Erythroblasts | H3K4me1 | 19 | 27 531 447 | GSE70660 |
| Erythroblasts | GATA1 | 19 | 118 938 490 | GSE36985 |
| HepG2 | MAZ | 28 | 51 465 720 | GSE31477 |

| | | | | |
|---|---|---|---|---|
| (hepatocellular carcinoma) | | | | |
| A549 (lung carcinoma) | MAZ | 28 | 39 848 427 | GSE91939 |
| GM12878 (B lymphoblastoid) | MAZ | 28 | 40 269 745 | GSE106046 |
| MCF-7 (mammary adenocarcinoma) | MAZ | 28 | 128 288 726 | GSE91633 |
| IMR90 (fetal lung fibroblast) | MAZ | 28 | 34 117 266 | GSE31477 |
